# Multisite Assembly of Gateway Induced Clones (MAGIC): a flexible cloning toolbox with diverse applications in vertebrate model systems

**DOI:** 10.1101/2024.07.13.603267

**Authors:** William Gillespie, Yuwen Zhang, Oscar E. Ruiz, Juan Cerda, Joshua Ortiz-Guzman, Williamson D. Turner, Gabrielle Largoza, Michelle Sherman, Lili E. Mosser, Esther Fujimoto, Chi-Bin Chien, Kristen M. Kwan, Benjamin R. Arenkiel, W. Patrick Devine, Joshua D. Wythe

**Author notes:** Corresponding author To whom correspondence should be addressed: Joshua D. Wythe, Ph.D., University of Virginia School of Medicine Charlottesville, VA 22903.

## Abstract

Here we present the Multisite Assembly of Gateway Induced Clones (MAGIC) system, which harnesses site-specific recombination-based cloning via Gateway technology for rapid, modular assembly of between 1 and 3 “Entry” vector components, all into a fourth, standard high copy “Destination” plasmid backbone. The MAGIC toolkit spans a range of in vitro and in vivo uses, from directing tunable gene expression, to driving simultaneous expression of microRNAs and fluorescent reporters, to enabling site-specific recombinase-dependent gene expression. All MAGIC system components are directly compatible with existing multisite gateway Tol2 systems currently used in zebrafish, as well as existing eukaryotic cell culture expression Destination plasmids, and available mammalian lentiviral and adenoviral Destination vectors, allowing rapid cross-species experimentation. Moreover, herein we describe novel vectors with flanking *piggyBac* transposon elements for stable genomic integration in vitro or in vivo when used with *piggyBac* transposase. Collectively, the MAGIC system facilitates transgenesis in cultured mammalian cells, electroporated mouse and chick embryos, as well as in injected zebrafish embryos, enabling the rapid generation of innovative DNA constructs for biological research due to a shared, common plasmid platform.

## INTRODUCTION

Modern biologists have an arsenal of available techniques to study cellular and molecular biological mechanisms in vitro and in vivo, yet most current investigative methods rely upon tools generated by recombinant DNA technologies. From CRISPR-based genome editing, to super resolution microscopy, almost all modern molecular biological investigations at some point require transgenic gene expression. However, despite the decreasing costs of commercial gene synthesis, complex expression vectors are still prohibitively expensive and difficult to synthesize (for example if they feature repetitive DNA elements or contain inserts greater than 3-4 kb in size), creating an obstacle for many laboratories. While some modern recombinant DNA methods, such as Golden Gate cloning ^1^ or Gibson assembly ^2^, have decreased barriers to assembling more complicated constructs and can obviate the need for unique restriction enzyme cut sites, many biologists still employ conventional “cut-and-paste” restriction enzyme/ligation-based techniques due to unfamiliarity or perceived difficulty with the protocols, or the cost of reagents. Furthermore, PCR-based cloning strategies (such as Gibson assembly) require extensive sequencing to confirm that no errant mutations have been introduced during PCR amplification. However, conventional restriction enzyme-based cloning also suffers from significant drawbacks, as it can be incredibly time consuming and laborious. Finally, these disadvantages are amplified when one wants to “mix and match” components from multiple vectors, such as combining a promoter from one plasmid with a cDNA payload from another source, and place them upstream of a unique fluorescent tag or a second cDNA in a bicistronic fashion. Unfortunately, these methods also suffer from a lack of interchangeable parts that are continually curated and expanded by the research community. However, the zebrafish developmental biology community created a blueprint, so to speak, for interchangeable and affordable multi-component cloning when they integrated the Tol2 transposon-based transgenic system with multisite Gateway cloning^3–5^.

The MultiSite Gateway cloning system (ThermoFisher) leverages site-specific recombination to insert anywhere from one to three DNA elements into a vector ^6,7^, obviating the need for restriction enzyme-based cloning. Indeed, this well-established, fast, and efficient system is ideally suited for rapidly combining insert cDNAs into established, expression ready plasmids, as well as assembling multiple components, such as tissue specific promoters with cDNAs and downstream fluorescent reporters ^6,7^. An array of gateway compatible “toolkits” enable modular assembly of various DNA elements^3–5,8^, in addition to numerous standalone gateway compatible transgenic expression plasmids ^9–11^. Furthermore, open reading frame libraries (ORFeomes) encoding protein coding sequences from humans ^12–17^, worms ^14,15,18^, frogs ^19^, and bacteria ^20–23^ have been cloned into Gateway-compatible plasmid backbones, further expanding the utility of this system. However, a significant drawback of many of these plasmid collections and toolkits is that they often do not employ the same arrangement of multisite Gateway recognition sequences. In essence, their assembly is often unique to a particular design logic, limiting their utility across experimental platforms (as the valuable plasmid resources available from one laboratory are not compatible with those generated by another group).

For this reason, we present MAGIC, a collection of novel, predominantly mammalian oriented, Gateway vectors that are compatible with existing Tol2 Entry and Destination vectors employed by the zebrafish community. Our Entry vectors span a variety of applications, ranging from tissue-specific promoters, to tunable/inducible gene expression systems, to fluorescent reporters, site-specific recombinase vectors, bicistronic expression systems, dual fluorescence and microRNA expression plasmids, optogenetics, and beyond. Importantly, the MAGIC system is compatible with ORFeome collections that have cDNA inserts in so-called “middle Entry” plasmids, such as pENTR1a, pDONR221, and pDONR223. This design allows for maximal flexibility and high throughput cloning. For example, by combining middle Entry fluorescent reporter clones and Destination vectors available at Addgene with the p5E pan-astrocytic *GLAST/EEAT-1/SLC1A3* promoter clone we generated, one could generate pLenti-Glast:EGFP, pLenti-Glast:mCherry, pTol2-Glast:EGFP, and pTol2-Glast:mCherry in a single set of parallel Gateway cloning reactions. In addition to these the novel Entry clones, we also created new Destination vectors for inducible gene expression in mammalian cells, as well as Cre/loxp and Dre/rox-dependent gene expression, and Destination plasmids with either *piggyBac* or Tol2 transposon / inverted terminal repeat [ITR] flanking sequences for use with their respective transposases.

We illustrate the utility of the MAGIC toolkit with multiple examples of primary research applications. Using an Entry vector encoding Luciferase and H2A-mScarlet-I (red fluorescent protein, hereafter referred to as mScarlet-I) ^24^ separated by a porcine teschovirus-1 2A (P2A) self-cleaving peptide ^25^, we compare the on/off kinetics of multiple Tet-inducible plasmids in cultured mammalian cells for fine tuning gene expression in vitro. Next, we add in the requirement for site-specific recombinases to show how Cre/lox and Dre/rox systems can be leveraged in cell culture with MAGIC vectors, and we combine the systems for doxycycline-tunable, Cre-dependent expression in vitro. Then, we show how a multisite Gateway system leveraging *piggyBac* transposase, analogous to the Tol2 system used in zebrafish, can be exploited for studies in cell culture, mice, and chickens. We then complement these tools with novel Tol2-based vectors to enable Cre-dependent expression within zebrafish in vivo. Finally, we offer a relational, open source database for managing plasmid records, electronic files, and sequences.

## RESULTS

### Overview

The MAGIC toolkit is only compatible with the use of 1, 2, and 3, Entry fragment Gateway cloning, which utilizes the *att* site-specific recombinase system from lambda phage^7^. By leveraging engineered and unique *att* sites, along with the commercially available enzymes that catalyze these reactions, one can directionally insert between one and three different DNA fragments into a single plasmid backbone. In this design, specific DNA fragments are first cloned between unique flanking *attL* recombination sites within an “Entry” vector (**Figure 1A**). A core feature of Gateway-based cloning is that *att*L sites react with complementary *att*R sites of identical numerical specificity, but they do not recognize other numerical Gateway sites. For example, L1 sequences will recognize and react with R1 sequences, and L3 sites with R3 sites, but L1 will not recombine with R3 (nor will L1 react with L2). In a single-Entry reaction, an Entry plasmid with *attL* sites flanking the insert is then mixed with a single “<u>Dest</u>ination” <u>v</u>ector (e.g. “Dest” or “DV” plasmid) that contains *att*R sites flanking the *ccd*B gene and a chloramphenicol resistance cassette (*ccdB*/Cm^R^) for negative-selection. The addition of a purified “LR” clonase enzyme mixture (commercially available from Invitrogen) composed of bacteriophage λ Integrase (Int) and Excisionase (Xis), as well as Integration Host Factor (IHF), mediates recognition and recombination of *attL* and *attR* sites via an “LR Reaction” (**Figure 1B**). In a two-insert fragment reaction, DNA from two Entry clones are arranged 5’ to 3’ within a single Dest backbone (for instance, to insert an ORF of interest followed by a 3’ IRES-fluorescent reporter into the same mammalian expression vector) (**Figure 1C**). Similarly, in a three-insert fragment reaction, DNA fragments from three different Entry vectors are directionally inserted into a suitable Destination plasmid (**Figure 1D**).

**Figure 1:**
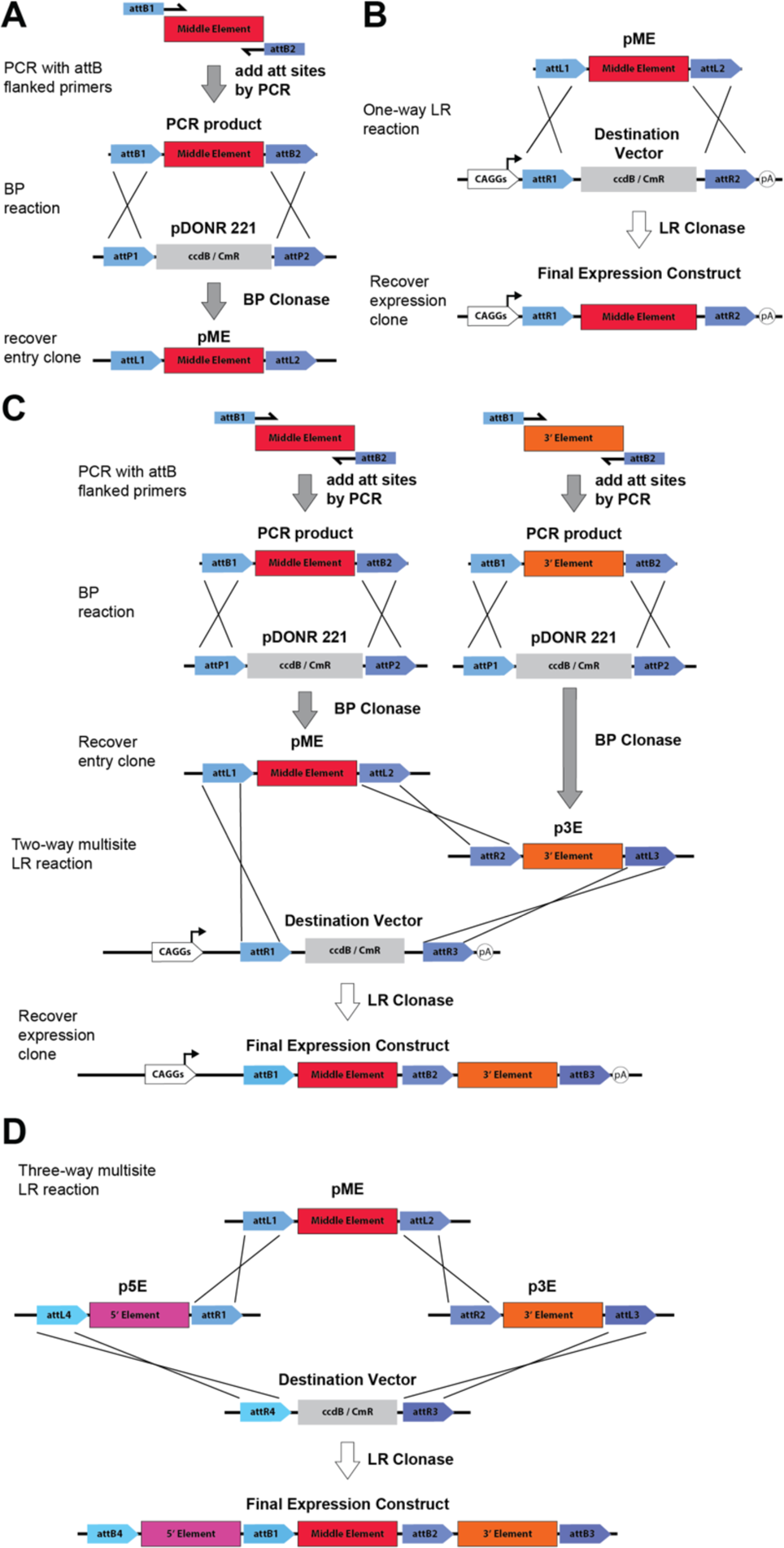
Gateway recombination reactions. **(A)** Generation of an insert by adding flanking attB1/B2 arms via PCR and subsequent BP clonase-mediated recombination into a pDONR backbone to generate an ENTRY clone. (**B**) Diagram of a one-way LR recombination of an entry clone into an DESTination vector to generate an expression clone. (**C**) Examples of simultaneous generation of a middle entry attB1/B2 flanked insert and an attB2/B3 flanked 3’ insert, both via PCR, and subsequent generation of a pME (middle ENTRY) and p3E (3’ ENTRY) clones via BP clonase-mediated recombination, then a two-way LR reaction with a DEST plasmid to generate an expression clone containing both inserts. (**D**) A simplified schematic of a three-way LR reaction between an attL4/attR1 5’ Entry clone (pME), an attL1/attL2 middle entry clone (pME), and an attR2/attL3 3’ entry clone (p3E) with a DEST vector.

The LR reaction, which contains a mixture of (1) unrecombined parental Destination vector harboring the intact *ccd*B/Cm^R^ casette, (2) unrecombined Entry vector^1^, as well as ^2^ the desired recombined Entry plasmid, are then transformed into bacteria. Transforming the reaction into an *E. coli* strain sensitive to CcdB (e.g. DH5α) negatively selects against the parental, non-recombined vector, as expression of CcdB (a DNA gyrase poison) will ultimately cause cell death^26^. Furthermore, as most Entry clones are generated in plasmids encoding Kanamycin resistance, and Dest vectors have cassettes encoding Ampicillin resistance, this positively selects for correctly recombined Destination vectors. Thus, using both negative and positive selection, Gateway cloning favors recombined, ampicillin resistant Destination clones that have replaced the *ccdB*/CmR cassette with the Entry DNA fragments (**Figure 1B-D**).

A key benefit to Gateway cloning is that it eliminates the laborious, and often inefficient steps of standard cut and paste restriction enzyme / DNA ligase cloning. Furthermore, this recombination-based cloning strategy also allows one to simultaneously arrange multiple DNA fragments within the same vector in cis, such as a promoter, ORF, and fluorescent reporter, all within a single reaction (saving precious time). However, as astutely pointed out by Villefranc and colleagues, taking full advantage of the Gateway system requires the establishment of Dest and Entry clones suited for functional studies in specific model organisms ^5^. While extensive *Tol2-* and *Tol1*-based kits are available for use in zebrafish, as well as one-off Destination lentiviral plasmids for mammalian cell culture work, these kits require the constant addition of novel promoters, ORFs, and the inclusion of newer biosensors, fluorescent reporters, and constructs to keep them current and maintain their utility. Herein we update the repertoire of available Entry clones, while also providing useful new mammalian *piggyBac* transposase-compatible Dest plasmids, murine gene targeting Dest vectors for use with Dre recombinase, and novel zebrafish *Tol2*-based Dest plasmids for single-, dual-, and three-way-Entry fragment Gateway cloning with the MAGIC kit.

### Making Entry clones and recombination reactions

There are three varieties of ENTRY clones for use in one-, two-, and three-insert Gateway cloning reactions: 5’ clones (p5E), middle Entry clones (pME), and 3’ Entry clones (p3E). The p5E vectors have flanking *attL4*-*attR1* sites, and typically contain a promoter element to drive expression *in vitro* or *in vivo*. The pME vectors contain inserts, usually a fluorescent reporter or gene of interest, flanked by *attL1*-*attL2* sites. The p3E vectors usually contain a fluorescent reporter (e.g. p3E-IRES-mCherry), an epitope tag (e.g. p3E-HA), or a polyA signal, flanked by *attR3*-*attL3* sites. The p5E, pME, and p3E ENTRY vectors are generated from three distinct “Donor” (pDONR) plasmids. Relying on cis DNA elements and an enzyme mixture containing INT and IHF (BP clonase), one can recombine DNA inserts flanked by homologous attB sites with Donor plasmids that have complementary attP sites (Figure 1A). The Donor constructs used for creating Entry clones in the MAGIC kit are as follows: for making p5E clones we use pDONR P4-P1R (with *attP4/P1R*); for creating pME clones we use pDONR221 (with *attP1/P2* sites); and for generating p3E clones we use pDONR P2R-P3 (with *attP2R/P3* sites). To generate Entry clones, we typically use polymerase chain reaction (PCR) to add the desired *attB* sites to the template of interest in vitro, or synthesize gene fragments with these flanking *attB* sites, followed by a BP reaction and transformation into bacteria (**Table 1**). As in attL/R recombination, the *attB* sites flanking an insert fragment will only recombine with homologous *attP* sequences within the pDONR vectors, and they must be numerically identical (e.g. *attB1* will recognize *attP1*, but *attB1* will not recombine with attP2). BP recombination between attP and B sites the pDONR clone into an ENTRY vector with flanking *attL* sites (Figure 1A).

**Table 1.**
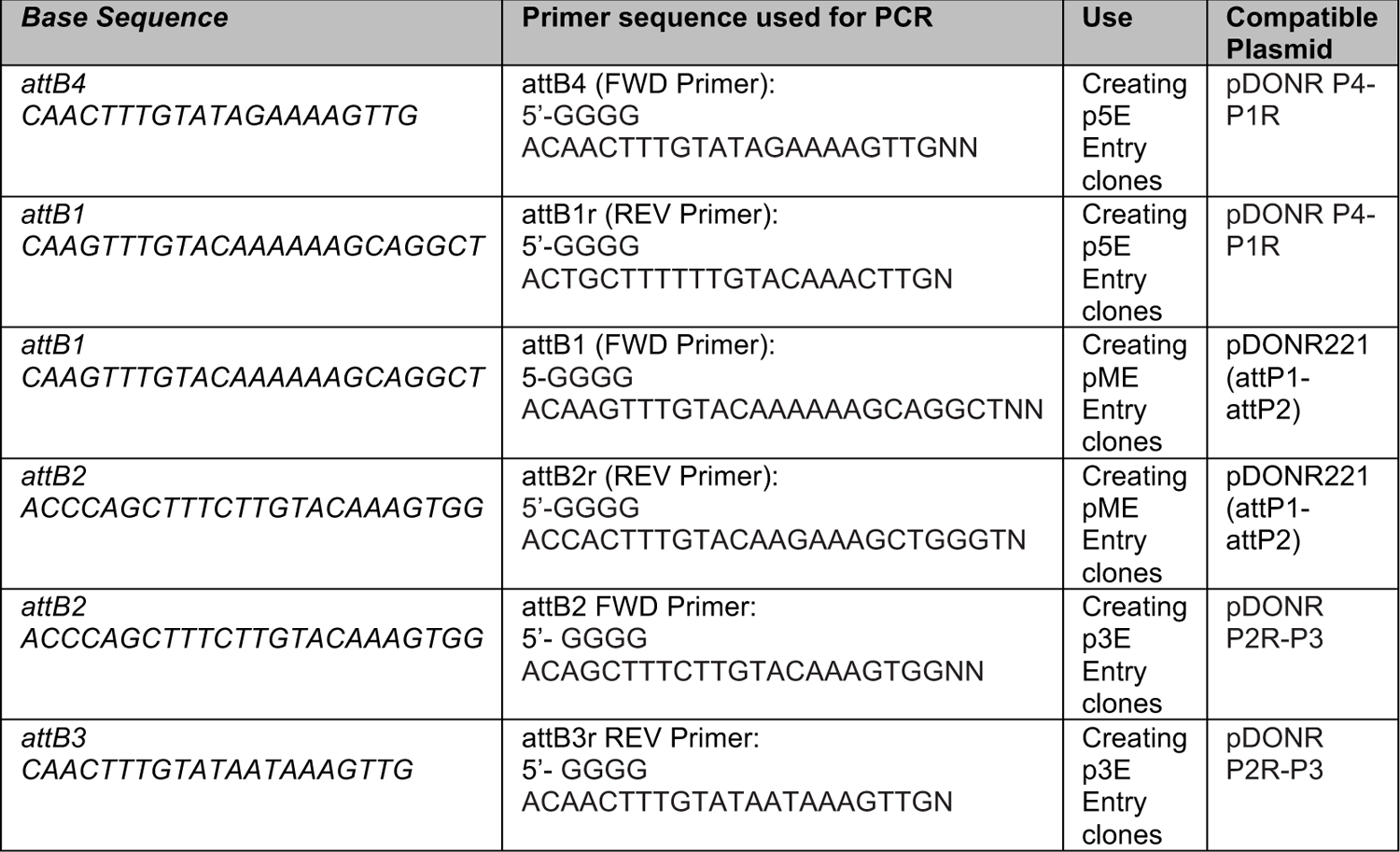
Att sites/sequences.

Importantly, the *attP* sites within pDONR vectors typically flank a *ccd*B cassette. Thus, negative selection for loss of the parental *attP* flanked *ccdB* pDONR insert favors recovering the desired new ENTRY clones, as non-recombined parental plasmids retaining the *ccd*B cassette should not be propagated in normal bacteria strains (non *ccd*B-).

Occasionally Entry plasmids will bear point mutations due to proof reading errors during PCR amplification, or synthesis errors within the primers used for PCR, or errors during manufacturing of a gene block or double-stranded DNA fragment. In instances when PCR or gene synthesis fails, or is not practical, each of the Entry vectors are available with multiple cloning sites instead of the *ccd*B/Cm^R^ cassette (e.g. p5E-MCS, pME-MCS, p3E-MCS from Addgene, or within the MAGIC kit) for conventional restriction enzyme-based cloning.

### Types of Entry clones

The components included in the MAGIC toolkit were chosen for their utility in either transient or stable transgenesis in mammalian cell culture or in vivo settings, as well as for transgenesis in zebrafish. They include several regulatory elements and promoters in 5’ Entry clones (CAGGS, Tetracycline Response Elements/TREs, etc.); reporter constructs (mTagBFP2, AmCyan, mCerulean and mCerulean3, EGFP, mKate2, mCherry, mScarlet-I, mScarlet-3 iRFP and tandem dimer iRFP (tdiRFP), etc.) in middle Entry vectors; as well as 3’ tags or fluorescent reporters (mScarlet, etc). Additionally, we have generated multiple Destination plasmids with 5’ and 3’ flanking *piggyBac* or *Tol2* inverted terminal repeats (ITRs) for stable transgenesis in vitro and in vivo. Below, the following sections will describe the 5’, middle, and 3’ Entry clones, as well as novel Destination vectors, as well as examples of the resultant expression using reagents provided by the MAGIC kit.

### 5’ ENTRY CLONES

The MAGIC system contains “empty” Destination backbones (or DEST vectors) that lack eukaryotic promoter elements, similar to the *Tol2* kits developed by other laboratories^3–5^. In this context, one needs to supply enhancer-promoter elements within 5’ ENTRY clones to drive expression of the middle and 3’ Entry clones (Figure 1). In general, all 5’ Entry clones are either amplified by PCR with flanking attB4/attB1r sites (**Table 1**), then cloned via a BP reaction into pDONR P4-P1R (Invitrogen), or they are inserted via conventional cut and paste restriction enzyme-based cloning into p5E-MCS (originally from Chi-Bin Chien’s lab). Regardless of their method of generation, all 5’ Entry clones described herein (denoted p5E) contain attL4/R1 sites flanking the promoter inserts (all p5E clones are listed in **Table 2**).

**Table 2.**
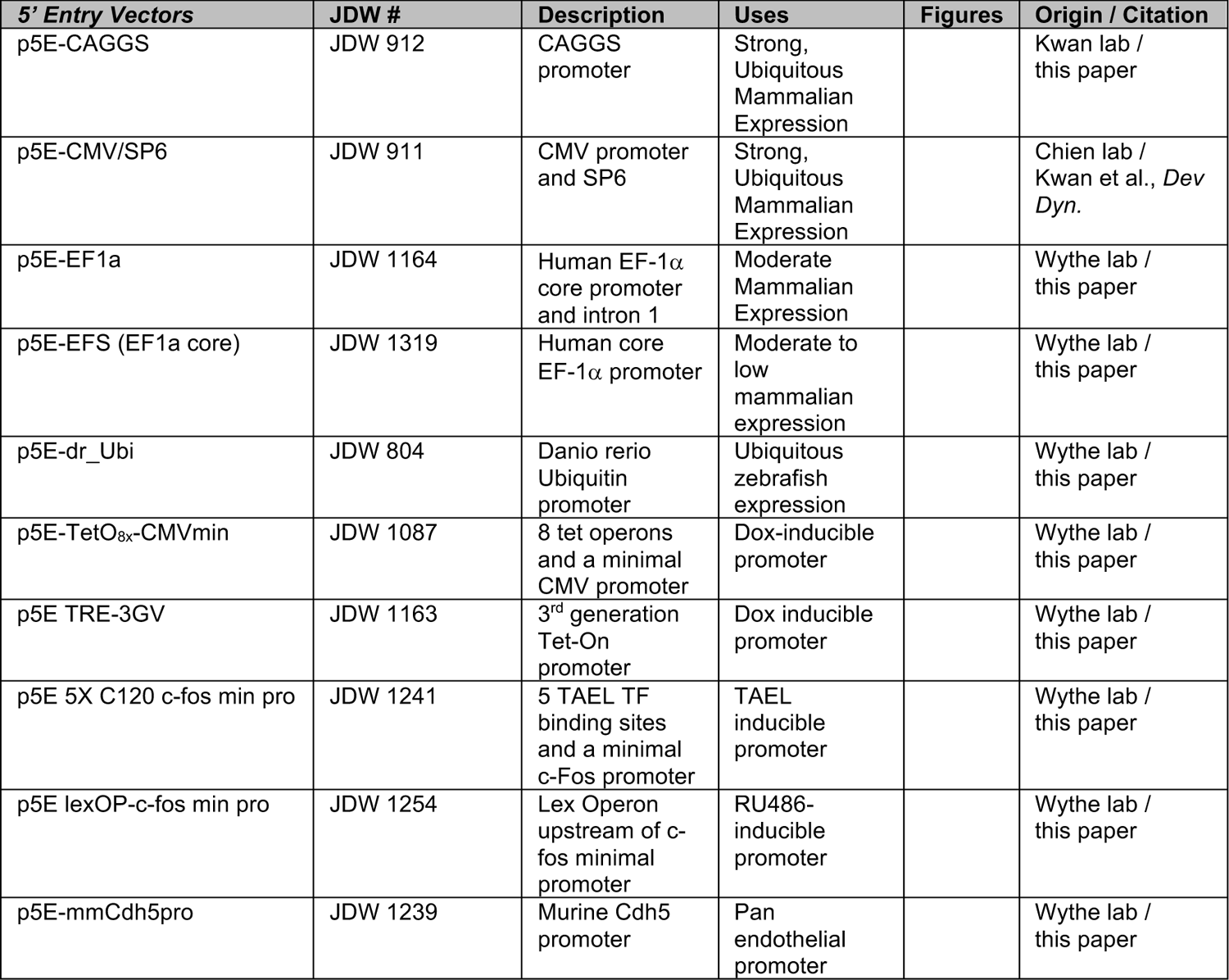
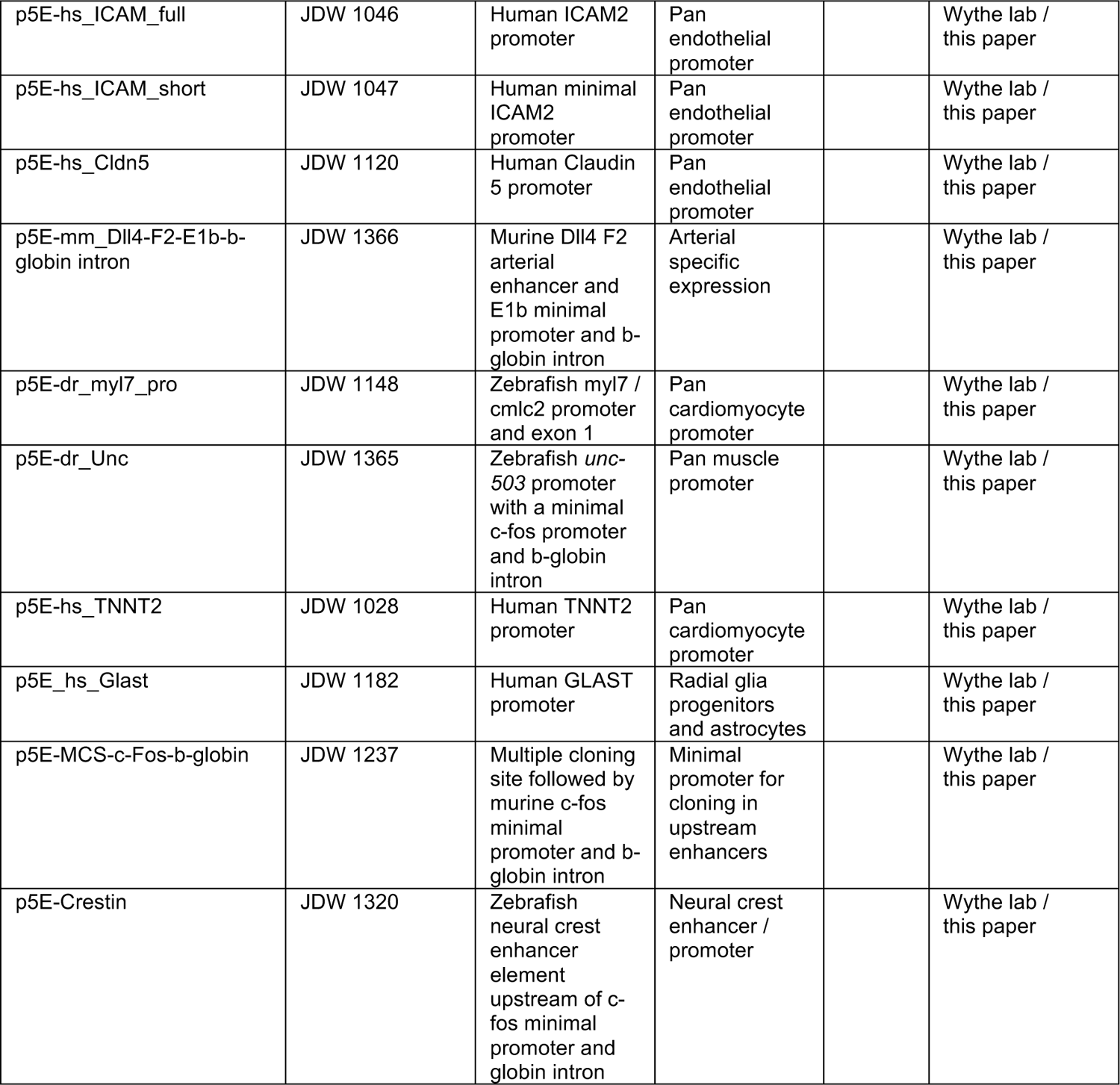
Existing and Novel Entry and p5E Vectors.

### Promoters for broad expression

Four p5E promoter constructs are included for broad, constitutive expression at various levels. We created p5E-CAG (or CAGGS) (JDW 912) for strong, ubiquitous expression in mammalian cells ^27,28^. We also include p5E-EF-1α (JDW 1164), which contains the human *Elongation Factor 1α* (EF-1α) promoter and ∼1 kb distal intron, as well as a shorter variant, p5E-EFS (JDW 1319) that only contains the EF-1 α core promoter (known as <u>EF</u>-1 α <u>s</u>hort, or EFS)^29^. We also provide a more moderately active promoter, p5E-*Ubi* (JDW 804), which contains a ∼3.5 kb insert spanning the promoter, exon1, and intron 1 of the zebrafish *ubiquitin* locus for constitutive expression in *Danio rerio*. Multiple other p5E constructs are available at Addgene, such as p5E-CMV/SP6 for broad expression in mammalian systems and p5E-*bactin2* for broad expression in zebrafish, both of which were described in one of the original zebrafish Tol2 Gateway kit manuscripts ^4^.

### Promoters for inducible expression

In addition to broad, constitutively active promoters for mammalian and zebrafish studies, we also provide p5E inducible promoter clones. The Tet system, based upon the high affinity recognition of the *Escherichia coli* Tet repressor protein (TetR) to the *tet* operator (TetO) DNA sequence, is a well-established, stringently controlled regulatory system for inducible gene expression in mammalian cells. In this system, either tetracycline (Tc), or its derivative doxycycline (Dox), binds to TetR and induces a conformational change that prevents this repressor protein from binding to the *tet* operator sequence. Subsequent fusion of the transcriptional activation domain of VP16 from Herpes simplex virus to TetR created the transcriptional activator tTA. In this constitutively active, “Tet-Off” system, tTA drives expression from *tetO* containing promoters, but the addition of Tc or Dox abolishes this tTA-tetO interaction and silences gene expression^30^. Subsequent manipulations resulted in the creation of the “Tet-On” system, where binding of a reverse tTA (rtTA) to DNA is induced by the presence, but not absence, of Dox^31^.

We created two Gateway compatible tetracycline regulatable promoter variants ^30,31^. The first, p5E-TetO_8X_-CMV_min_ (JDW 1087), contains a 2^nd^ generation Tet response element, also known as a TRE, composed of eight 19-bp *tet* operator sequences located upstream of a 104-bp minimal CMV promoter ^32,33^. The second, p5E-TRE-3GV (JDW 1163), contains a 3^rd^ generation TRE promoter composed of seven 19-bp *tet* operator sites upstream of a 12-bp pTight hybrid sequence^34^ followed by a modified 68-bp minimal CMV promoter that contains a synthetic TFIIB binding site and TATA box^35^, which together ensure low basal activity and maximal responsiveness to doxycycline and high performance with lentivirus and retrovirus ^36^.

We have also adapted an optogenetic-based regulatory system optimized for gene expression in zebrafish that relies on the naturally occurring light-responsive transcription factor, EL222, found in *Erythrobacter litoralis*. EL222 consists of a LOV (light-oxygen-voltage) domain followed by a helix-turn-helix (HTH) DNA-binding domain (DBD)^37^. When illuminated with blue light (450-490 nm), EL222 dimerizes and binds to a regulatory element known as C120^38,39^. N-terminal addition of the transactivation domain of KalTA4 to EL222 generates a robust, photo-activatable gene expression system with rapid on/off kinetics known as TAEL (for TA4-EL222)^40^. In the second iteration of this system, TAEL 2.0, the carboxy-terminal addition of the nucleoplasmin nuclear localization signal (NLS) to TAEL (termed TAEL-N), and the use of five concatenated C120 regulatory elements (5xC120) followed by the murine basal *c-Fos* promoter (which allows for high expression levels with minimal background in zebrafish) ensures robust light-inducible expression^41–43^. Accordingly, here we provide a p5E-*5xC120-c-Fos-minpro* (JDW 1241) construct for generating TAEL responder vectors for use in the TAEL/C120 system.

Finally, we also adapted the hormone inducible binary LexA/LexO system for targeted gene expression in either zebrafish^44,45^ or in mammals. In this system, a ligand-dependent fusion between the DNA-binding domain (DBD) of the bacterial LexA repressor (residues 1-87) ^46,47^ and a truncated ligand-binding domain (LBD) of the human progesterone receptor (residues 640-914) is joined to the activation domain of the human NK-kB/p65 protein (residues 283-551)^48^. In the presence of the progesterone agonist mifepristone (also known as RU-486), this chimeric LexPR transactivator (LexA^DBD^-PR^LBΛ1^-p65^AD^) binds to the synthetic LexA operon (LexOP), which is composed of four *ColE1* operator sequences^49^ placed upstream a minimal 35S promoter sequence from the Cauliflower Mosaic Virus^50^, to drive expression of an effector cassette. Here, we provide a p5E clone where the cauliflower mosaic virus minimal promoter has been exchanged for the murine *c-Fos* promoter (p5E-lexAOP-c-fos, JDW 1254).

### Promoters for tissue-specific expression

The MAGIC toolkit also contains multiple pan-endothelial specific promoters, such as that of murine *VE-Cadherin* (*Cdh5*) and human *ICAM2* (both the full length 397 bp promoter and minimal 139 bp sequence)^51–54^, as well as that of *Claudin 5* (*Cldn5*)^55^, and the arterial-specific enhancer of murine *Delta like 4*, F2 (*Dll4-F2*)^56^. We also provide the zebrafish *myl7* (previously referred to as *cmlc2*) promoter^4^ and the 550 bp human *cardiac troponin T* (*TNNT2*) promoter ^57^ for cardiomyocyte specific expression in zebrafish and mice, respectively, as well as a 503 bp pan-muscle promoter (*-503unc*)^58^, and an 844 bp zebrafish-specific neural crest driver (*crestin*)^59^, both as 5’ entry clones (**Table 2**). For studies in mice, chick, and cell culture models, we generated p5E clones containing the human *GLAST* (*EAAT1*/ *SLC1A3*) promoter for expression in radial glia and astrocytes^60,61^. Finally, we also provide a p5E clone with an extensive multiple cloning site (KpnI, XhoI, HindIII, SpeI, BglII, PacI, BamHI, PmeI) for insertion of any enhancer element upstream of a minimal 93 bp murine *c-Fos* promoter followed by the 652 bp rabbit *β-globin* intron (JDW 1237). Collectively, this suite of ubiquitously active strong and moderate transcriptional drivers, as well as inducible and tissue-specific promoters, will facilitate experiments in various eukaryotic cell culture lines, as well as in vertebrate animal models.

### MIDDLE ENTRY CLONES

The MAGIC system typically uses middle Entry clones to supply fluorescent reporters or genes of interest, similar to the *Tol2* kits developed by other laboratories ^3–5^. Unless stated otherwise, all middle Entry vectors were produced by PCR amplification of the desired template using attB1/B2-flanked primers, or direct synthesis of the fragment with flanking attB1/B2 sites, followed by a BP reaction with pDONR221 (Invitrogen). In some cases, we also used a modified pDONR221 that we created with a novel multiple cloning site (EcoRI-SalI-BamHI-KpnI-SmaI-NotI-XhoI-EcoRI), known as pME-MCS (JDW 455), for cut and paste restriction enzyme-based cloning. The inserts within these pME vectors usually contain a consensus Kozak sequence ^62^ (5’-GCCACC-3’) for optimal ribosome binding and initiation of translation upstream of the first start codon (ATG). Unless otherwise noted, all middle Entry clones contain a stop codon at the 3’ end, and thus cannot be used for N-terminal fusions to a protein of interest by placing them upstream of a p3E clone. In cases where middle Entry clones contain multiple inserts, the coding regions of unique cDNAs are separated by a viral 2A peptide (2A). The use of a 2A-generated ribosomal skip event should ensure more equivalent protein production of independent polypeptide inserts, unlike the use of an internal ribosomal <u>e</u>ntry <u>s</u>ite (IRES), which is less likely to generate equal expression levels of the downstream insert relative to the upstream cassette^4,25,63,64^. A complete list of all pME vectors is provide in **Table 3**.

**Table 3.**
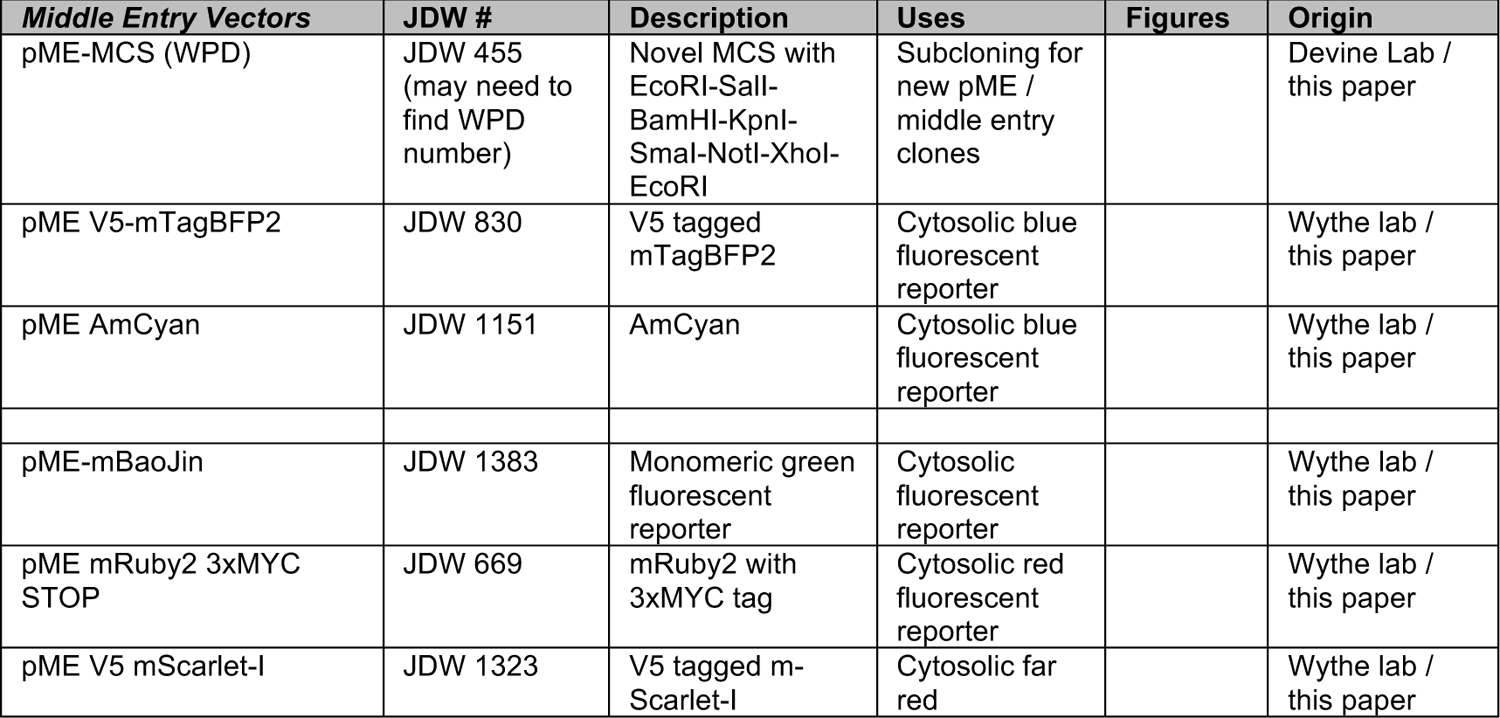
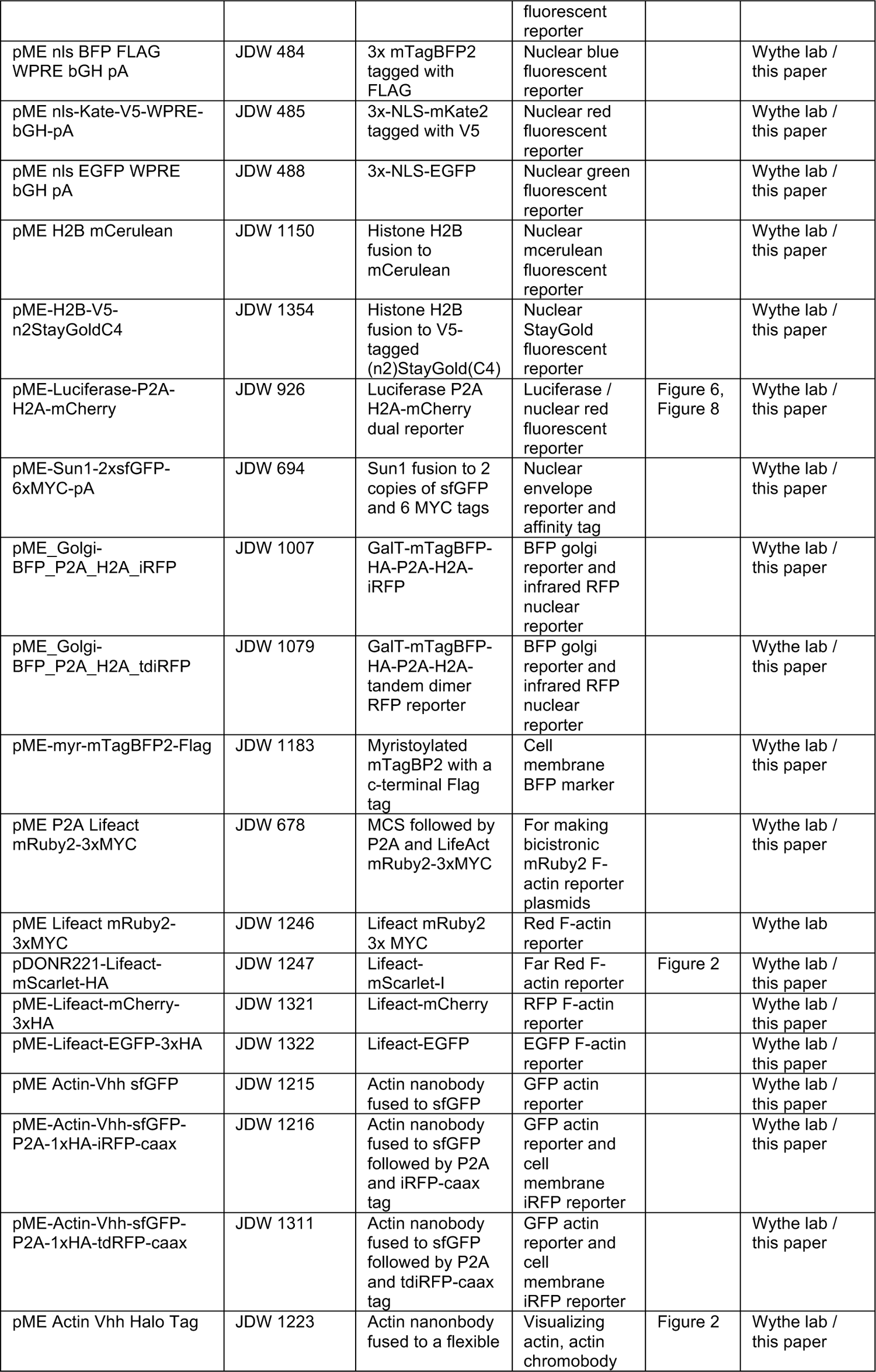
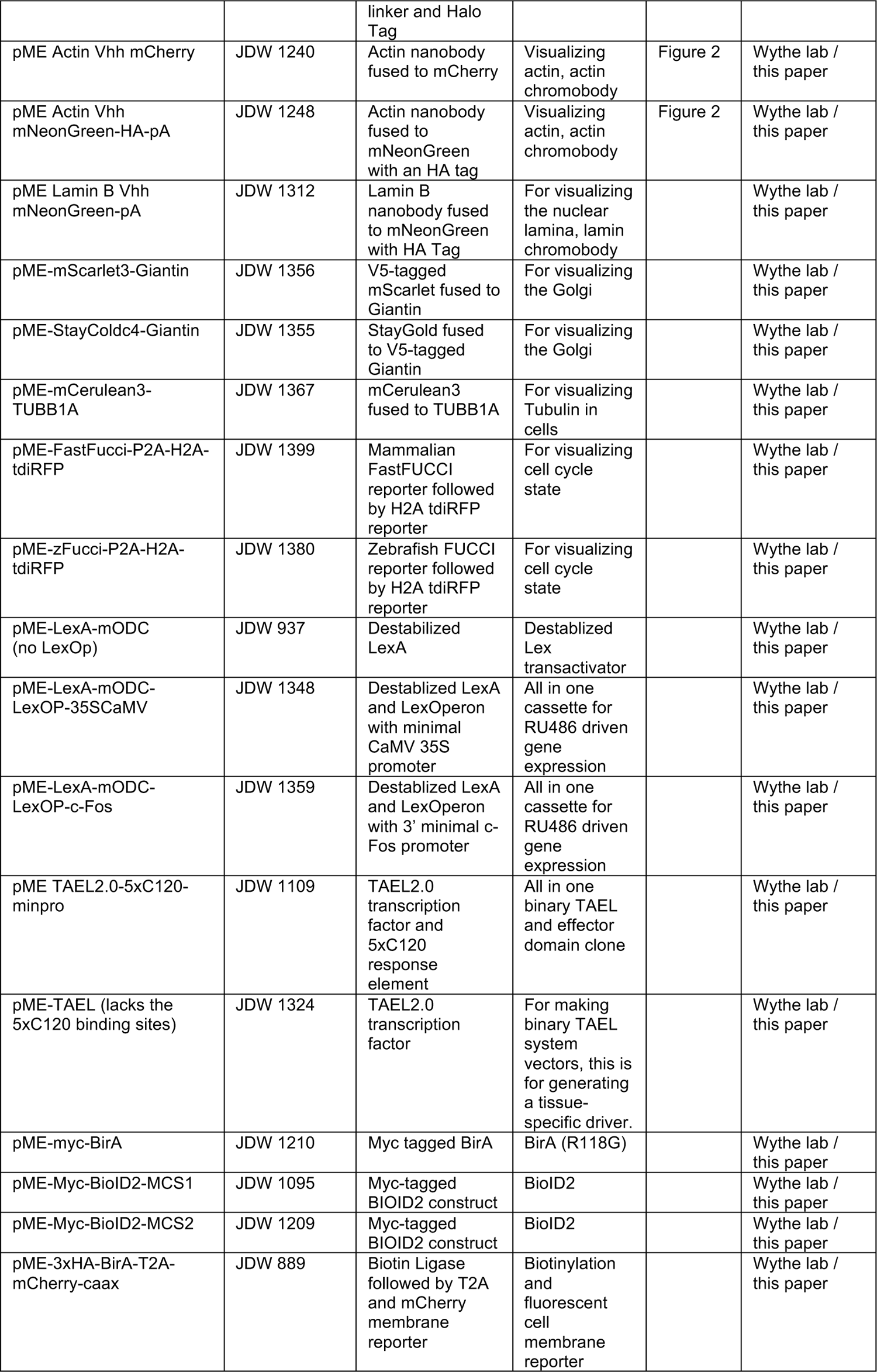
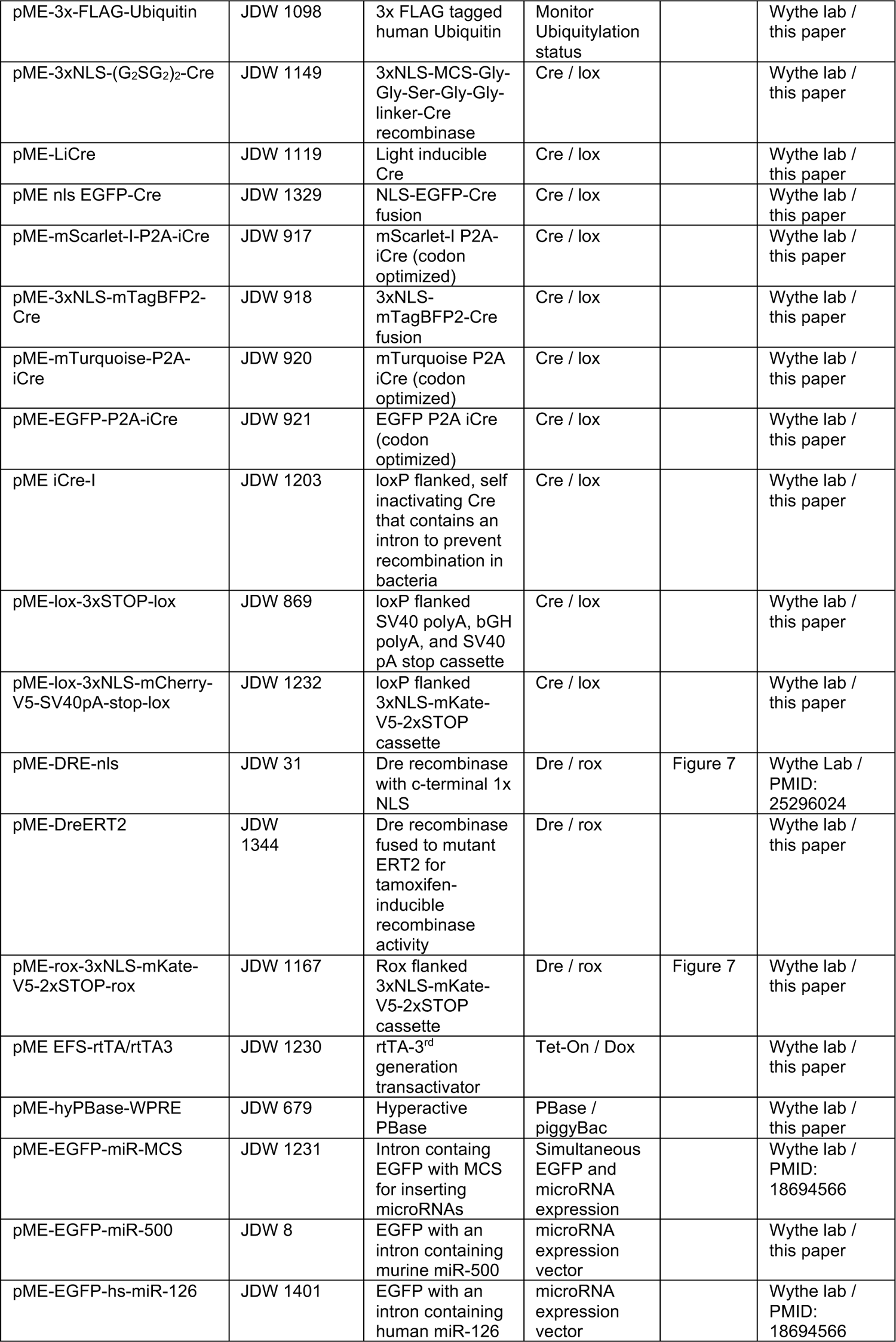
Existing and Novel Entry and pME Vectors.

### Fluorescent reporters

The MAGIC kit includes middle Entry clones encoding 16 distinct fluorescent reporter proteins that span the imaging spectrum, including mTagBFP2^65^, AmCyan1^66,67^, mCerulean^68^, mCerulean3^69^, mTurquoise2^70^, EGFP^71^, StayGold^72^, mCherry^73^, mRuby2^74^, mKate2^75^, mScarlet-I^24^, mScarlet3 and mScarlet-I3^76^, mCardinal^77^, iRFP and tandem iRFP dimer (tdiRFP)^78^. This collection of fluorescent proteins (FPs) was intentionally chosen to maximize fluorescence signal while spanning the spectrum from blue (mTagBFP2) to cyan (mCerulean, mCerulean3, mTurquoise2), green (EGFP, mClover3, mNeonGreen, StayGold), red (mCherry, mKate2, mScarlet), far red (mCardinal) and near infrared (iRFP and tandem dimer iRFP/tdiRFP).

Relatively newer fluorescent reporters, such as the red variants mScarlet-I ^24^ and improved versions mScarlet3 and mScarlet-I3^76^, and the green variant StayGold^72^ were included due to their reported superior stability and brightness for live imaging. Like other bacterial phytochrome-derived near infrared FPs, iRFP (34.6 KDa) and tdiRFP (71.5 kDa) (also known as iRFP670) require the metabolite biliverdin to form a fluorophore^79–82^. These two near-infrared FPs were chosen because their wavelength is outside the absorption spectrum of hemoglobin and melanin (which occur extensively in mammalian tissues), thus the signal should be less attenuated and yield a better signal to background ratio than other fluorescent proteins in intact tissues, facilitating in vivo imaging^83^. While some reports suggest expression of these near-infrared FPs may require exogenous biliverdin in zebrafish, they are sufficient to produce robust signal in vivo^84^. For most of the FPs without readily available antibodies we added an epitope tag to facilitate western blot analysis or immunostaining.

Numerous permutations of these different FPs are supplied in the MAGIC kit to allow for maximal flexibility and labeling of different subcellular compartments. Specifically, for nuclear labelling, we provide pME plasmids containing a 3x nuclear localization signal fused to a fluorescent protein followed by a WPRE and bGH polyA (nls-mTagBFP2-FLAG, nls-mKate-V5, nls-EGFP), as well as histone H2B or H2A chimeric fluorescent proteins (H2B-mCerulean / JDW 1150; mCardinal-H2B), as well as an anti-lamin B nanobody ^85^ fused to mNeonGreen to label the nuclear lamina (pME anti-laminB-mNeonGreen / JDW 1312) and a fusion of the nuclear envelop protein Sun1 to 2 copies of super folder GFP (pME-Sun1-2xsfGFP-6xMyc / JDW 681). Of note, Sun1-2xsfGFP-6xMYC allows for both visualization and immunoprecipitation of nuclei ^86,87^ (Figure 2A).

**Figure 2:**
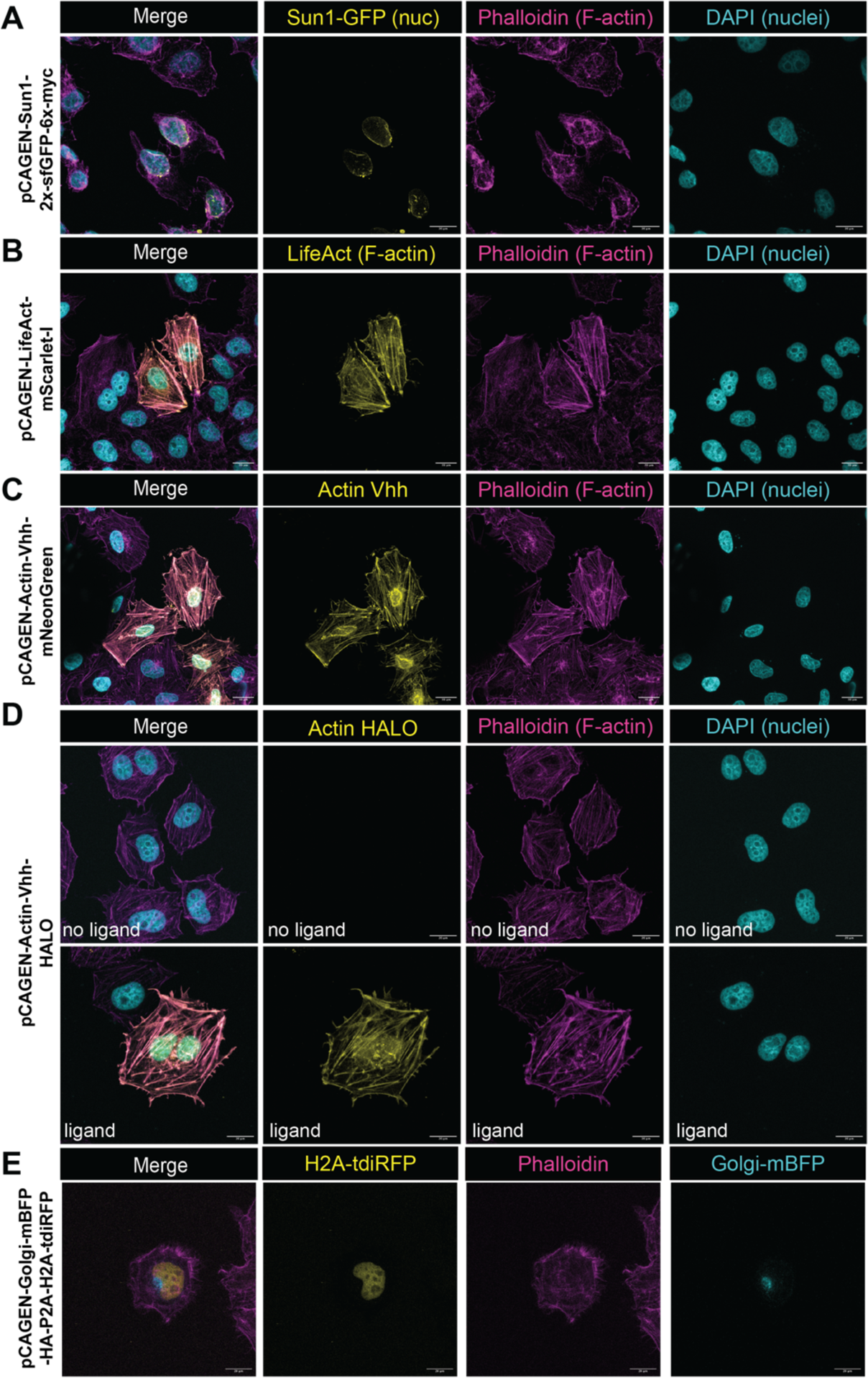
Subcellular Labelling Using M.A.G.I.C. Kit Components. Clones were made using middle entry clones described in **Table 3** and the pCAGEN-DEST vector to facilitate labelling of (**A**) nuclei with a 2x super folder GFP fusion to the nuclear envelope protein Sun1, (**B**) filamentous actin with a fusion of mScarlet to the short F-actin probe Lifeact, (**C**) a pan-actin probe generated from a fusion of a nanobody that recognizes actin (actin Vhh) fused to mNeonGreen for a live reporter, as well as (**D**) a fusion of the same actin nanobody fused to Halo tag for flexible use of antibody or dye labeling (in this case Janelia dye JFX646), with and without dye, and (**E**) dual labeling of the Golgi and nucleus with GalT-mTagBFP and an H2A-tdiRFP probe. All pME constructs were LR recombined into pCAGEN and transiently transfected into HeLa cells. Some cells were fixed, permeabilized, and counter stained with DAPI to label nuclei and phalloidin to label F-actin, and then imaged by confocal microscopy.

For visualizing cytoskeletal dynamics, we focused on two strategies. The first employs a 17 amino acid peptide tag derived from yeast Abp140^88^. This short fragment, known as Lifeact, recognizes filamentous actin (F-actin) structures in eukaryotic cells (Figure 2B). Unlike tools such as fluorescent phalloidin, actin-FP chimeric proteins, or bulky anti-actin antibodies, the small size of Lifeact enables the live study of endogenous intracellular actin. We provide multiple red FPs conjugated to Lifeact in middle entry clones (pME-Lifeact-mRuby2-3xMYC/JDW 1246, pME-Lifeact-mScarlet/JDW 1247, pME-Lifeact-mCherry / JDW 1321, and pME-Lifeact-EGFP / JDW 1322), as well as one plasmid with an MCS located upstream of a P2A cleavage peptide and a fusion of Lifeact to mRuby2 that allows for the easy generation of unique bicistronic expression constructs (pME-MCS-P2A-Lifeact-mRuyb2-3xMYC / JDW 678) (Figure 2B). As recent reports highlighted unanticipated artifacts at the cellular, organ, and whole animal level resulting from overexpression of Lifeact-fluorescent reporters ^89–91^, we also provide a nanobody that recognizes α-actin fused to fluorescent reporters (sometimes referred to as a chromobodies) for visualizing cytoskeletal dynamics in cultured cells. Specifically, we fused Actin Vhh to superfolder GFP (sfGFP) ^92^ to create pME-Actin-Vhh-sfGFP (JDW 1215) (Figure 2C). We fused this same actin nanobody to a versatile c-terminal 33 KDa HaloTag ^93^ (pME-Actin-Vhh-Halo / JDW 1223). The self-labeling HaloTag has been engineered to form an irreversible, covalent bond with its ligands. In the context of imaging, while a HaloTag itself does not fluoresce, intracellular labeling using small, cell-permeable dyes with high brightness and photostability enables robust labeling of subcellular structures in vitro and in vivo^94^ (Figure 2D).

We also include general cytoplasmic localized fluorescent reporters (pME-AmCyan1/ JDW 1151, pME-V5-mScarlet-I/JDW 1323), and FP chimeric fusions to human Giantin to facilitate labeling of the Golgi (pME-mScarlet3-Gigantin/JDW 1356; pME-(n2)StayGold(c4)-Giantin/JDW 1355) as well as tubulin to visualize cytoskeletal dynamics (pME-mCerulean3-TUB/JDW 1367).

Additionally, we provide middle Entry clones encoding more than one ORF to facilitate simultaneous labeling of different subcellular compartments or structures. For example, JDW 1216 encodes a nanobody recognizing actin fused to 2 copies of superfolder GFP followed by a P2A cleavage peptide and HA epitope fused iRFP with a c-terminal CAAX tag from H-Ras to allow for visualization of actin as well as the plasma membrane (pME-Vhh-actin-sfGFP-P2A-iRFP-caax). We modified this construct, by replacing the iRFP insert with tdiRFP to generate pME-Actin-Vhh-sfGFP-P2A-HA-tdiRFP-caax (JDW 1311). Other clones include pME-Golgi-mTagBFP-HA-P2A-H2A-iRFP (JDW 1007) and pME-Golgi-TagBFP-HA-P2A-H2A-tdiRFP (JDW 1079) to simultaneously label the Golgi (via a fusion of human GalT to mTagBFP2) and the nucleus (via H2A.F/Z tagged iRFP or tdiRFP). Recombination of this Entry clone with pCAGEN-Dest (JDW 472) yielded pCAGEN-Golgi-mTagBFP2-HA-P2A-H2A-tdiRFP (JDW 1219). Transient transfection of this construct into HeLa confirms the fidelity of these subcellular labelling inserts in mammalian cells (Figure 2E).

### Cell Cycle Tools

In addition to these reagents, we provide fluorescent probes to indicate the status of the mammalian and zebrafish cell cycle based on <u>F</u>luorescence <u>U</u>biquitin <u>C</u>ell <u>C</u>ycle Indicator (FUCCI) technology^95^. This system exploits the licensing factor Cdt1 and its inhibitor Geminin. The abundance of these two antagonistic factors oscillates during the cell cycle in a mutually exclusive manner dependent upon the E3 ligases SCF^Skp2^ and APC/C^Cdh1^, which control the destruction of Cdt1 and Geminin, respectively. Thus, the FUCCI system allows direct visualization of G1 and S/G2/M by virtue of differential accumulation of the fluorescent probes, as Cdt1 levels peak in G1 prior to the onset of DNA replication, and then decline rapidly after the initiation of S phase. Conversely, Geminin levels are high during S and G2 phases, but decrease during late mitosis and G1. Here we provide entry clones for the mammalian FastFucci reporter^96^, where simultaneous, equimolar expression monomeric Kusabira Orange 2 (mKO2)–human CDT1(amino acids 30–120) [mKO2-hCDT1(30-120)] and monomeric Azami Green (mAG)–human GEM(amino acids 1–110) [mAG-hGem(1-110), separated by a T2A peptide cleavage sequence (pME-FastFucci, JDW 1399). We modified this plasmid to also include a tdiRFP reporter downstream for labelling of all cells, regardless of cell cycle state (pME-FastFucci-P2A-H2A-tdiRFP, JDW 1399) and provide a pCAGEN-FastFucci-P2A-H2A-tdiRFP expression vector (JDW 1402). To complement these tools, we also provide a modification of a previous zebrafish FUCCI reporter^97,98^, where 3xFLAG-mCerulean-zGem(1-100)-P2A-mCherry-zCdt1(1-190) is followed by a P2A and tdiRFP such that cells in late G1 are labelled by mCherry, while those in S/G2/M display cerulean fluorescence, and all cells are labelled with iRFP (pME-zFucci-P2A-H2A-tdiRFP, JDW 1380).

### microRNA Tools

For studying micro-RNAs, we provide an entry vector, pME-EGFP-miR-MCS (JDW 1231), where the ORF of EGFP is interrupted by an internal intron that contains an MCS. Insertion of the region surrounding a microRNA into this MCS enables the simultaneous transcription of EGFP and a primary-microRNA (pri-miRNA) by polymerase II. After intronic splicing, the pri-miRNA containing the stem loop hairpin is then processed by the microprocessor (i.e. DGCR8 and Drosha) to generate a ∼60-70 nt pre-miRNA, which is exported to the cytoplasm and then processed by Dicer and Ago2. Eventually the single stranded ∼22 nt miRNA is loaded into RISC, where it will facilitate post-transcriptional repression via base-pairing to its target mRNAs^99^. Herein, we provide the endothelial-enriched miRNA *miR-126*, which is embedded within the human *Epidermal Growth Factor Like-domain 7* (*EGFL7*) gene (pME-EGFP-hs_*miR-126*, JDW 1401) ^100^. We also provide miR-500, embedded within an intron of the murine *Chloride Voltage-Gated Channel 5* (*CLCN5*) gene on the X chromosome (pME-EGFP-mm_*miR-500*, JDW 8). miR-500 is expressed in the embryonic mouse in the central nervous system and asymmetrically in the limb bud ^101^. LR recombination into a commercial vector containing the human *Eukaryotic Translation Elongation Factor 1 Alpha* (EF-1α/*EEF1A1*) (pEF1-DEST51, Invitrogen) promoter ensures robust expression in vitro as assessed by qRT-PCR (data not shown). Expression vectors for both constructs are provided (**Table 6**).

### Cre and Dre site-specific recombinase tools

The MAGIC kit also contains middle Entry clones designed for site-specific recombinase (SSR) experiments. We, and others, previously showed that the SSR Dre recombinase works well in vitro and in vivo in murine models and demonstrates minimal, if any, cross-reactivity with loxP sites in vivo ^102,103^. We include middle Entry clones encoding DRE recombinase fused to a C-terminal NLS, as well as one containing DRE recombinase fused to mutant estrogen receptor (pME-DRE-ERT2) to allow for tamoxifen-inducible DRE activity ^102,103^. We provide CAG driven expression vectors of both a constitutive Dre (pCAGEN-Dre-nls, JDW 1131) and tamoxifen-inducible Dre (pCAGEN-DreERT2, JDW 486). Additional Dre reporter tools will be discussed in the Destination Vector section below.

A suite of plasmids encoding direct FP fusions to Cre recombinase are included, such as pME-nls-EGFP-Cre (JDW 1329) and pME-nlsTagBFP-Cre (JDW 918), as well as P2A bi-cistronic Cre encoding fluorescent plasmids (pME-mTurquoise-P2A-iCre, pME-EGFP-P2A-iCre, pME-mScarlet-I-P2A-iCre), and a clone with two flexible Gly-Gly-Ser-Gly-Gly (G_2_SG_2_) linker peptides flanking an MCS upstream of the Cre ORF (pME-3xNLS-2x(G_2_SG_2_)-Cre) (JDW 1149) to enable direct amino terminal fusions to Cre by restriction enzyme-based cloning. We also provide a middle entry clone containing a self-excising Cre insert that contains 5’ and 3’ loxP sites flanking the insert, as well as a synthetic chimeric intron separating the 5’ and 3’ halves of the ORF to prevent recombination in *e. coli* (JDW 1203). In terms of *loxP* compatible Cre reporter systems, we provide a pME vector, pME-lox-3xSTOP-lox (JDW 869), that can act as a transcriptional stopper when placed upstream of a p3E fluorescent reporter (see below), as well as a novel 3xNLS-mCherry reporter flanked by loxP sites, pME-lsl-3xNLS-mCherry-V5-SV40pA (JDW 1232).

Herein we also provide a middle Entry vector encoding a single chain, fast-responding, light-activatable Cre (LiCre) for optogenetic control of Cre recombinase activity ^104^. LiCre is a chimeric fusion between the photropin 1 LOV2 domain of *Avena sativa* (AsLOV2) engineered for light-induced dimerization that is fused to the αA-terminal helix of a destabilized Cre recombinase carrying mutations in its N- and C-terminal domains (p.E340A, p.D341A) ^104^. LiCre can be activated solely by illumination with blue light and doesn’t require additional chemicals or other co-factor proteins (such as in a split component system). We have also generated CAG-driven LiCre (JDW 1220) and validated its activity in vitro (Figure 3A**,B**). Notably, lentiviral transduction of LiCre via stereotactic injection and simultaneous optogenetic implantation, followed by optical induction 14 days later, led to robust recombination of a *Rosa26-lsl-TdTomato* (e.g. Ai14) fluorescent reporter in mice within one week (Figure 3C-G).

**Figure 3:**
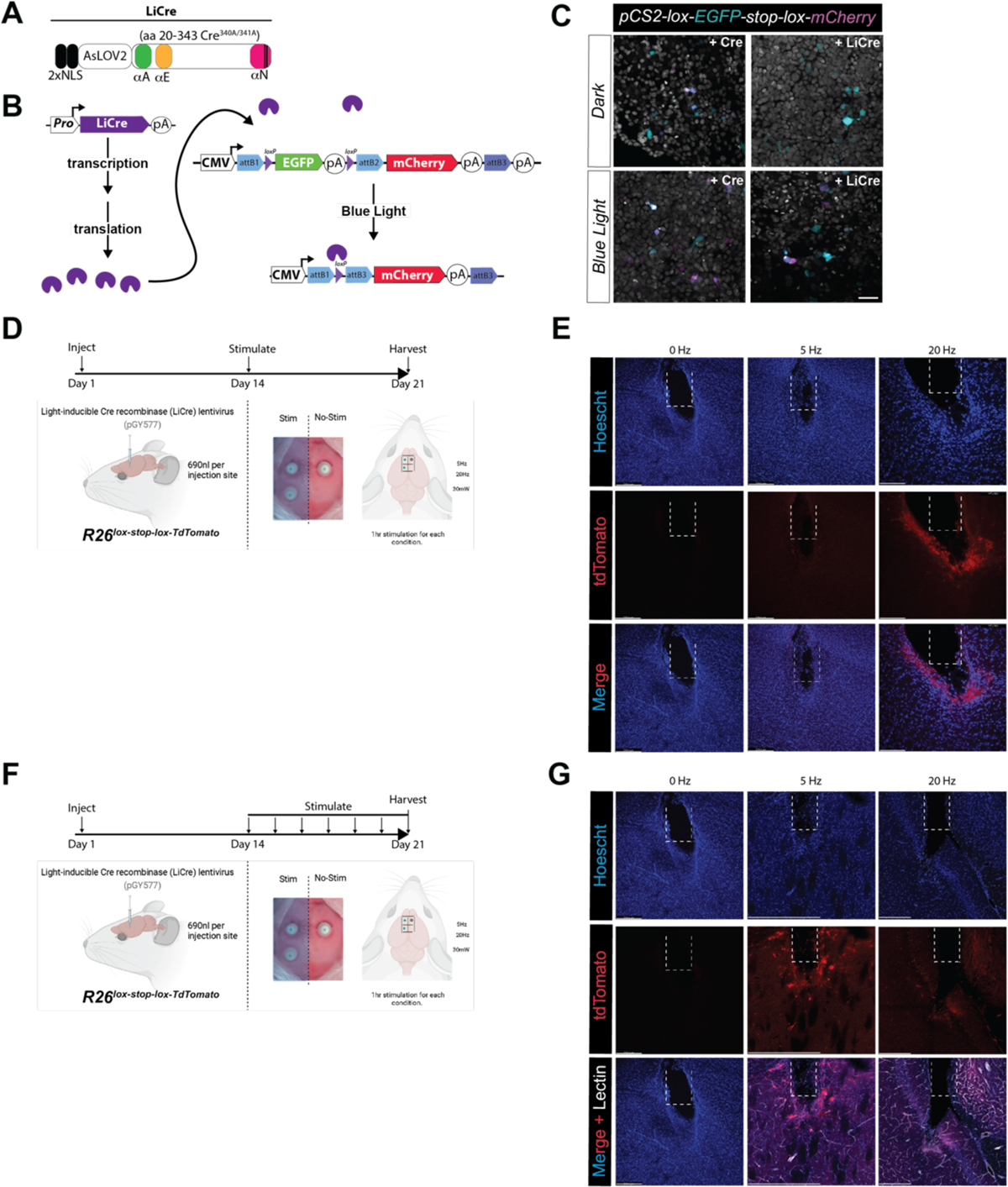
An Optogenetically Controlled Cre Recombinase System. **(A)** Schematic of the LiCre cassette, composed of a 2x NLS tag followed by the photoreceptor domain of *Avena sativa* (AsLOV2) fused to a N-terminally truncated variant of Cre recombinase with destabilizing mutations in its C-terminal domains. (**B**) LiCre activity is induced within minutes of illumination with blue light without the need for additional chemicals or components. (**C**) Transfection of LiCre in mammalian cells, along with a Cre-dependent switch reporter, confirms activation in the presence, but not absence, of blue light. Scale bar = 50µM (**-G**) Experimental design for lentiviral transduction via stereotactic injection to the murine brain, followed 2 weeks later by varying illumination protocols. (**E**) Robust recombination of a *Rosa26^lox-stop-lox-TdTomato^* fluorescent reporter, as detected by TdTomato fluorescence within the murine brain, is evident one week after a single hour of stimulation at 5 Hz and 20 Hz. (**G**) Robust recombination is also evident following serial illumination, 1 h/day at 5 Hz. Variable Cre activity, as evidenced by TdTomato expression, is seen after stimulation at 20 Hz, for 7 days.

### Light-inducible transcriptional activators

In addition to these light-activable Cre plasmids, we included a modified pME-TAEL2.0 “all in one” plasmid that encodes the TAEL-nls chimeric light-inducible transcriptional activator and responder cassette. In the presence of blue light, this chimeric transcriptional activator binds to cognate DNA regulatory elements to drive expression of responder cargo in a light-inducible manner (Figure 4A)^42^. We created a pME vector containing the TAEL2.0 ORF followed by a transcriptional pause site and a 5X concatenated C120 binding domain (pME-TAELn-pause-5xC120, JDW 1109), and combined this with a Tol2 destination vector and a ubiquitous promoter to drive expression of TAEL2.0 followed by the C120 cassette and a p3E donor containing an mCherry reporter. Injection of this construct into a *Tg(kdrl:EGFP)* transgenic line and selective illumination confirms of mCherry in animals exposed to a blue light induction paradigm, but not those animals raised in the dark (Figure 4B-G).

**Figure 4:**
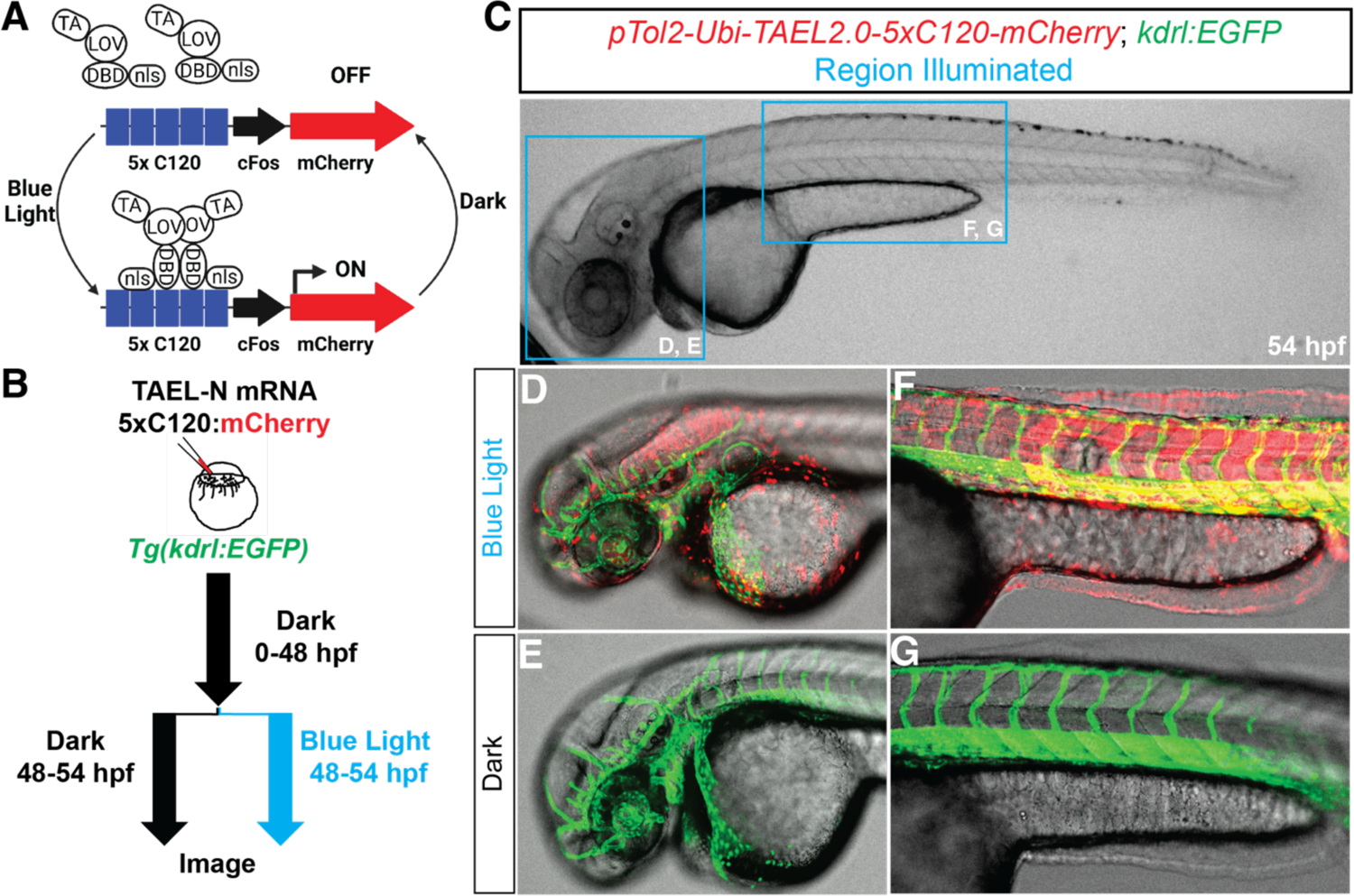
An Optogenetically Inducible in vivo Transcriptional Effector System. **(A)** Design of the TAEL2.0 system, pioneered by Woo and colleagues, where fusion of the KalTA4 transcriptional activator (TA) domain to blue light-oxygen-voltage-sensing ^90^ domain and DNA binding domain (DBD) of the Erythrobacter litoralis HTCC2594 transcription factor EL222, followed by a c-terminal NLS tag, allows for 405 nm light-induced conformational change and dimerization of TAEL (Kal<u>TA</u>4-<u>EL</u>22), allowing it to bind and initiate transcription from the C120 regulatory element upstream of a minimal c-Fos promoter and mCherry reporter cassette. (**B**) Experimental paradigm for injection into a endothelial EGFP reporter, *Tg(kdrl:EGFP)* at the 1-cell stage, followed by illumination with blue light. (**C**) Whole mount, sagittal view, phase image of a 54 hpf embryo containing the Tg(*kdrl:EGFP)* reporter and illuminated in the indicated, blue boxed regions in panel. (**D-G**) Blue light illumination led to robust induction in pTol2-Ubi-TAEL-N-5xC120-mCherry-injected embryos within a few hours of exposure to blue light, while embryos maintained in the dark did not show induction of mCherry.

### LexA/LexO tools

To complement our p5E LexO promoter clone, we generated pME plasmids containing the LexPR transcriptional activator. However, previous reports cited slow off-rate kinetics using this system, with expression of an EGFP reporter persisting for up to 5 days post withdrawal of mifepristone ^44^. To overcome this drawback, we created a pME vector encoding the LexPR fused c-terminally to the PEST domain of murine ornithine decarboxylase (amino acids 422-461, known as mODC), which reduces t_1/2_ of EGFP six-fold in mammalian cells ^105,106^. We supply this pME-LexPR-mODC construct (JDW 937) to facilitate the generation of tissue-specific transgenic lines using the Tol2 system.

The two elements of Lex transactivator and synthetic Lex operon are traditionally combined in trans in zebrafish to regulate gene expression ^44^ (Figure 5A**,B**). Here we sought to first determine if these elements could be combined in an “all in one” vector system (Figure 5C).

**Figure 5:**
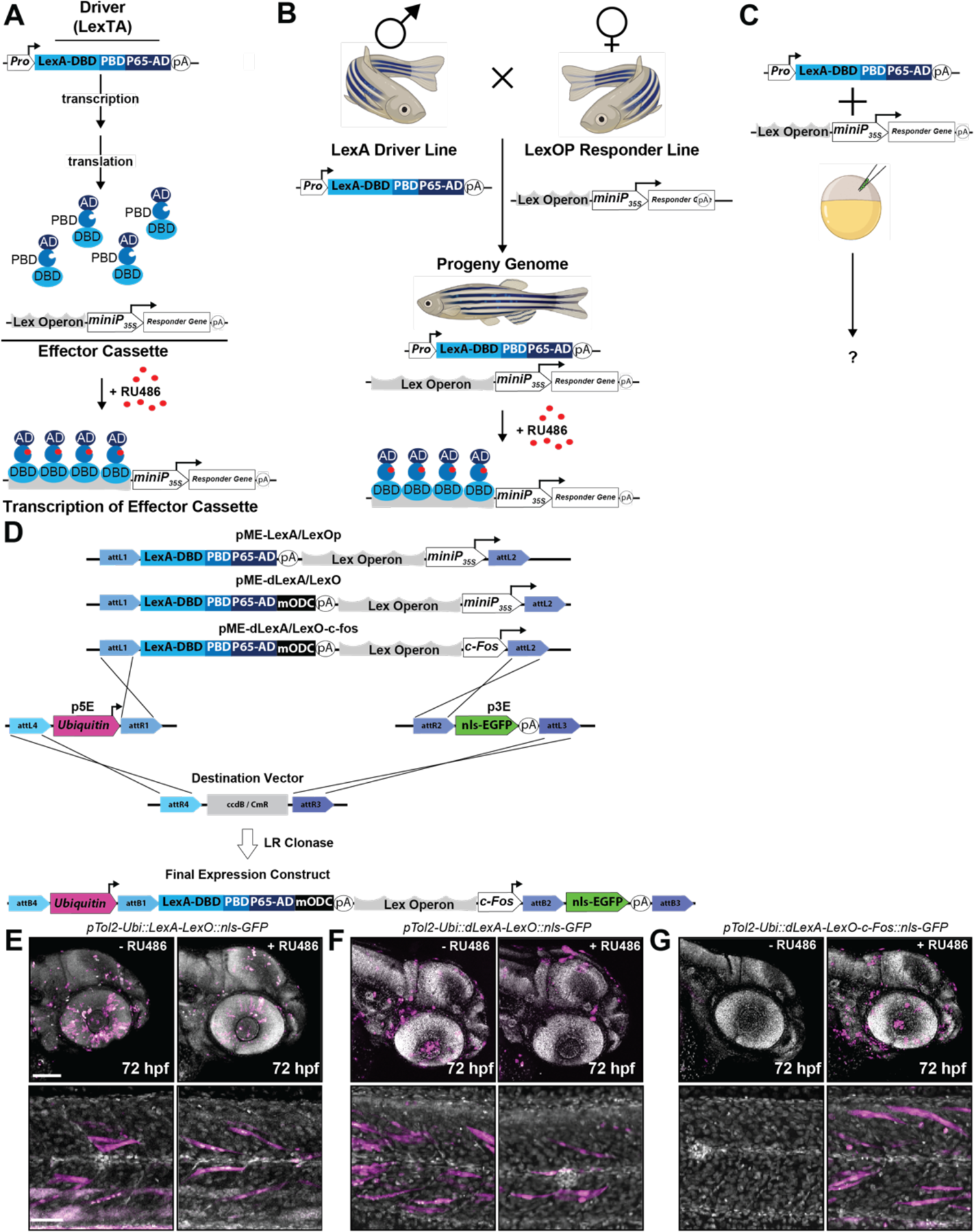
The LexA/LexO Inducible System. **(A)** The mifepristone (RU486)-inducible LexA/LexO gene expression system consists of two components. The first is a “Driver”, in this case the LexA transcriptional activator (LexTA). LexTA is a fusion between the Lex DNA binding domain-the progesterone ligand binding domain-and the transcriptional activator domain of NK-κB p65. This hybrid, or chimeric, transcription factor binds to the synthetic steroid, mifepristone (RU486) and binds the second component, an “Effector Cassette”, composed of a synthetic lexA operator (LexO) upstream of a cauliflower mosaic virus 35S minimal promoter (*miniP35s*) in a ligand-dependent manner to drive transcription of the downstream gene. (**B**) Typically, the driver and effector are maintained in different lines and only combined in *trans*. (**C**) Potential design of combing the LexA and LexO responder cassette within the same embryo for simpler genetics using an “all in one” approach. (**D**) Modification of the LexA system by addition of a destabilization domain from ornithine decarboxylase (mODC) to the LexTA to prevent accumulation of the transcriptional activator and replacement of the cauliflower miniP35s promoter with the minimally active mammalian *c-Fos* promoter. (**E-G**) Confocal images following treatment with either vehicle (DMSO) or RU846 (2 μM) from 24-72 hpf. Note has leaky transgene induction of this “all in one” system unless LexTA is destabilized and the *miniP35s* is replaced with the *c-Fos* minimal promoter (compare the far left and middle panels to the far right panels). Head image scale = 100 µm. Somite image scale = 50 µm.

In this context, a tissue-specific promoter (provided by a p5E plasmid) would drive expression of the LexPR in the tissue of interest and the downstream LexOP promoter (with LexPR and LexOP supplied in the same pME vector) would drive expression of a downstream effector cassette (supplied by the p3E clone). Accordingly, we generated three pME LexA/LexO clones. In the first, we combined the LexPR transactivator with the LexOP and minimal 35S promoter. In the second, we decreased the half-life of LexPR by fusing it to an mODC destabilization domain, but retained the 35S minimal promoter. In the third iteration, we combined the destabilized LexPR with a Lex operon followed by the minimal *c-fos* promoter (Figure 5D). We then placed each of these pME clones downstream of a *ubiquitin* promoter (p5E-Ubi), followed by a nls-EGFP reporter (supplied via a p3E-nls-EGFP-pA). Injection of a single vector containing the traditional LexA/LexO componenets (*pTol2-Ubi-LexA-LexO-CAMV35S-nls-EGP-pA*) resulted in robust nls-GFP expression, even in the absence of RU486 treatment.

Destabilizing the LexPR seemed to have a minimal impact on reporter expression, as nls-EGFP was still evident in the absence of RU486 in *pTol2-Ubi-dLexA-LexO-CMV35S-nls-EGFP-pA* injected embryos. However, combing a destabilized LexPR together with the LexOperon and a minimal c-fos promoter (*pTol2-Ubi-dLexA-LexO-c-fos-nls-EGFP-pA*) mitigated leaky reporter induction, demonstrating that an all in one approach can be used for RU486-inducible gene expression in zebrafish (Figure 5E-G).

### 3’ ENTRY CLONES

Similar to previous Tol2-based Gateway kits, we routinely use 3’ ENTRY clones to supply a polyadenylation (polyA) signal. Unless stated otherwise, all 3’ ENTRY vectors were produced by PCR amplification of the desired template using attB2/B3r-flanked primers, or direct synthesis of the fragment with flanking attB2/B3 sites, followed by a BP reaction with pDONR P2R-P3 (Invitrogen). In some cases, we used a modified p3E ENTRY vector that contains a multiple cloning site flanked by the P2R and P3 sites, known as p3E-MCS (Addgene #75174)^107^, for cut and paste restriction enzyme-based cloning. All 3’ Entry clones are designated as p3E, and the DNA inserts are flanked by *attR2/L3* sites. (all p3E clones are listed in **Table 4**).

**Table 4.**
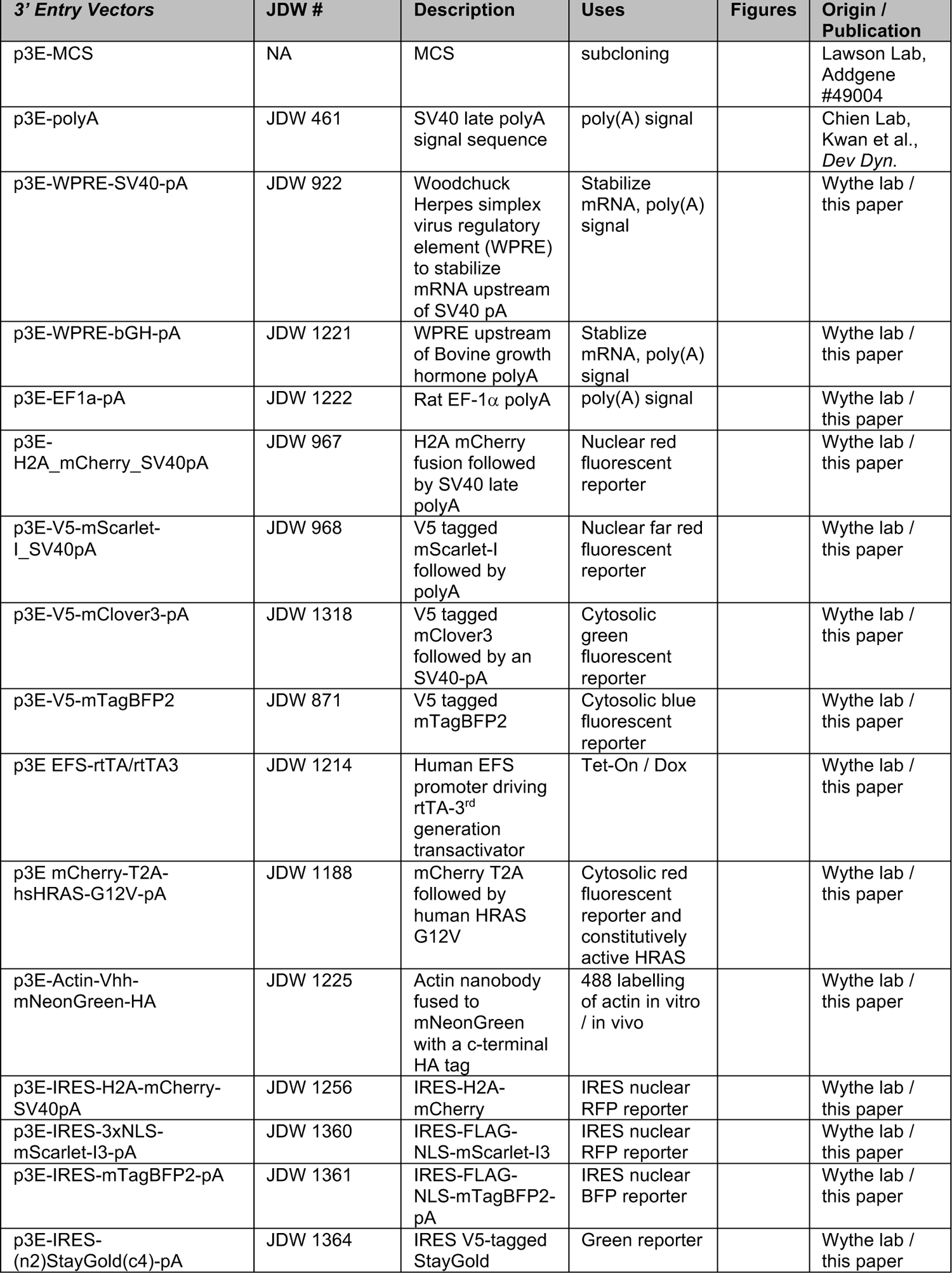
Existing and Novel Entry and p3E Vectors.

### polyA signal

Addgene has multiple p3E vectors containing the SV40 late polyA signal sequence (Addgene # 49004, #75174) ^108^. In zebrafish, as well as other systems, the SV40 late polyA is more effective than the SV40 early sequence in stabilizing mRNA transcripts and promoting translation^4,109,110^. We improved upon this by generating a p3E with a tripartite (α, β and ψ subunits) 588 bp regulatory element that enhances mRNA expression: the <u>w</u>oodchuck hepatitis virus <u>p</u>ost-transcriptional regulatory <u>e</u>lement (WPRE) ^111^, followed by the SV40 late polyA signal sequence.

We also provide alternative p3E polyA signal sequences, including a WPRE and the bovine growth hormone polyA (p3E-WPRE-pbGH-pA / JDW 1221) and a rat EF1a polyA (p3E-rEF1-pA / JDW 1222).

### Fluorescent markers and fusion proteins

Using a middle Entry (pME) clone encoding a gene of interest without a stop codon can allow one to make direct, in frame fusions to p3E inserts, as demonstrated elegantly in previous Tol2 Gateway kits ^3–5,107^. As a variety of these are publicly available at Addgene, we focused our efforts on generating standalone, p3E fluorescent reporter cassettes that could function as reporters for site-specific recombination of pME loxP or rox flanked stop cassettes. We provide several plasmids for this purpose: p3E-mCherry-SV40pA, p3E-H2A-mCherry-SV40-pA, p3E-V5-mScarlet-SV40-pA, p3E-Actin-Vhh-mNeonGreen-HA, and p3E-mTagBFP2. Furthermore, we created multiple p3E vectors encoding amino terminal fluorescent fusions to both wildtype and constitutively active KRAS and HRAS (**Table 4**) (as shown in Figure 9).

### Tet-inducible transactivators

To enable the generation of “all in one” Tet-inducible gene expression vectors, and to complement our p5E TRE tet-responsive promoter vectors, we generated a p3E plasmid that contains an upstream bGH polyA, then a synthetic polyA and a transcriptional pause from human *α2 globin*, followed by a minimal human EFS promoter driving expression of a reverse tetracycline transactivator (rtTA-3G). rtTA-3G (also known as Tet-On 3G, or rtTA-V10) is a virally evolved construct with improved intrinsic rtTA activity and effector-sensitivity^112^. In this 3^rd^ generation Tet-On system, the presence of Dox (but not Tc) induces a conformational change in the Tet-On 3G transactivator protein that allows it to bind the *tet* operator sequences within tetracycline response element (TRE)-containing vectors to drive expression of downstream effector cassettes (in our case, an ORF provided by a pME vector). Using the plasmids supplied within the MAGIC kit, one can assemble a p5E-TRE promoter element, followed by a pME encoding a gene of interest, and this p3E-EFS-rtTA-3G (JDW 1214) to create an “all in one” tetracycline responsive expression vector.

## DESTINATION VECTORS

Many others have created useful Destination vectors that provide Tol2 transposon elements which mediate genomic integration for effective transient and stable transgenesis in zebrafish, and recent work has also leveraged an analogous Tol1 transposase system ^113^. The MAGIC toolkit predominantly contains mammalian compatible Destination vectors. We provide both promoterless Destination plasmids for three insert Gateway cloning, as well as single site, “plug and play” vectors that already contain a promoter and polyadenylation signal sequence. Many of the promoterless, multisite Destination plasmids we provide contain flanking *piggyBac* ITR sequences for use with pBase transposase for robust genomic integration in eukaryotic cells, such as in cultured mammalian cells, or in chick or mouse via electroporation. Originally isolated from the cabbage looper moth genome^114^, *piggyBac* is a DNA transposon that with an enormous cargo limit (>200 kb) ^115,116^, it is active in many cell types, and it mediates long-term expression in mammalian cells in vivo and in vitro ^117,118^. *piggyBac* inserts at target sites with the sequence TTAA, but has little selectivity for particular regions of the genome ^119–121^. Given these advantages, we generated several new Destination vectors for single and multisite Gateway cloning in vectors with flanking piggyBac ITRs. We also provide gene targeting vectors (Cre and Dre compatible) for murine transgenesis, as well as a suite of ubiquitous and tissue specific promoter driven Destination vectors for use in mammalian systems, as well as novel multisite Gateway Tol2 and I-Sce vectors for zebrafish transgenesis.

### CAGGS Destination vectors for single ENTRY cloning

Multiple pME-compatible DESTINATION plasmids (e.g. with an *attR1-ccd*B-CmR-*attR2* cassette) exist for transient expression in mammalian cells, such as those with an EF1a promoter (e.g. pEF-DEST51, Invitrogen) or a CMV promoter (e.g. pCDNA-3.2-V5-DEST, Invitrogen). A particularly versatile group of Destination vectors utilize the pCS2 backbone, which features a CMV immediate early enhancer (CMV-IE) and CMV promoter upstream of an SV40 late polyadenylation signal sequence, ensuring robust expression in mammalian cells ^4,5,122^. DEST vectors utilizing this backbone have the attractive feature of also being able to generate mRNA for injection into *Xenopous laevis* and zebrafish embryos via in vitro transcription due to the presence of an SP6 promoter immediately 5’ to the DEST cassette^4,5,122^.

However, the CMV-IE enhancer/promoter cassette is often less active than synthetic variants in transiently transfected cells. An example of a more robust synthetic promoter is CAG, which is a fusion between the <u>C</u>MV immediate early enhancer and the *<u>AG</u>* promoter (hence, CAG). The *AG* promoter itself is a hybrid between the promoter and first intron of chicken *β-actin* and a fragment spanning intron 2 and exon 3 of the rabbit *β-globin* gene that contains an effective splice acceptor^27,123^. This resultant plasmid, pCAGGS (also referred to as pCAG), contains a downstream rabbit *β-globin* polyA signal sequence, and an SV40 origin of replication in a pUC13 backbone, and is more active than CMV in most transient transfection assays ^28^. We converted pCAGEN, a second-generation version of this vector that contains an expanded multiple cloning site^124^, into a Destination vector by inserting an *attR1-ccdB*-Cm^R^-*attR2* Gateway cassette into the MCS (Figure 6A) pCAGEN-DEST allows for robust expression in cultured mammalian cells, and similar to the parental vector it should effectively drive expression following electroporation into murine retinas^124^.

**Figure 6:**
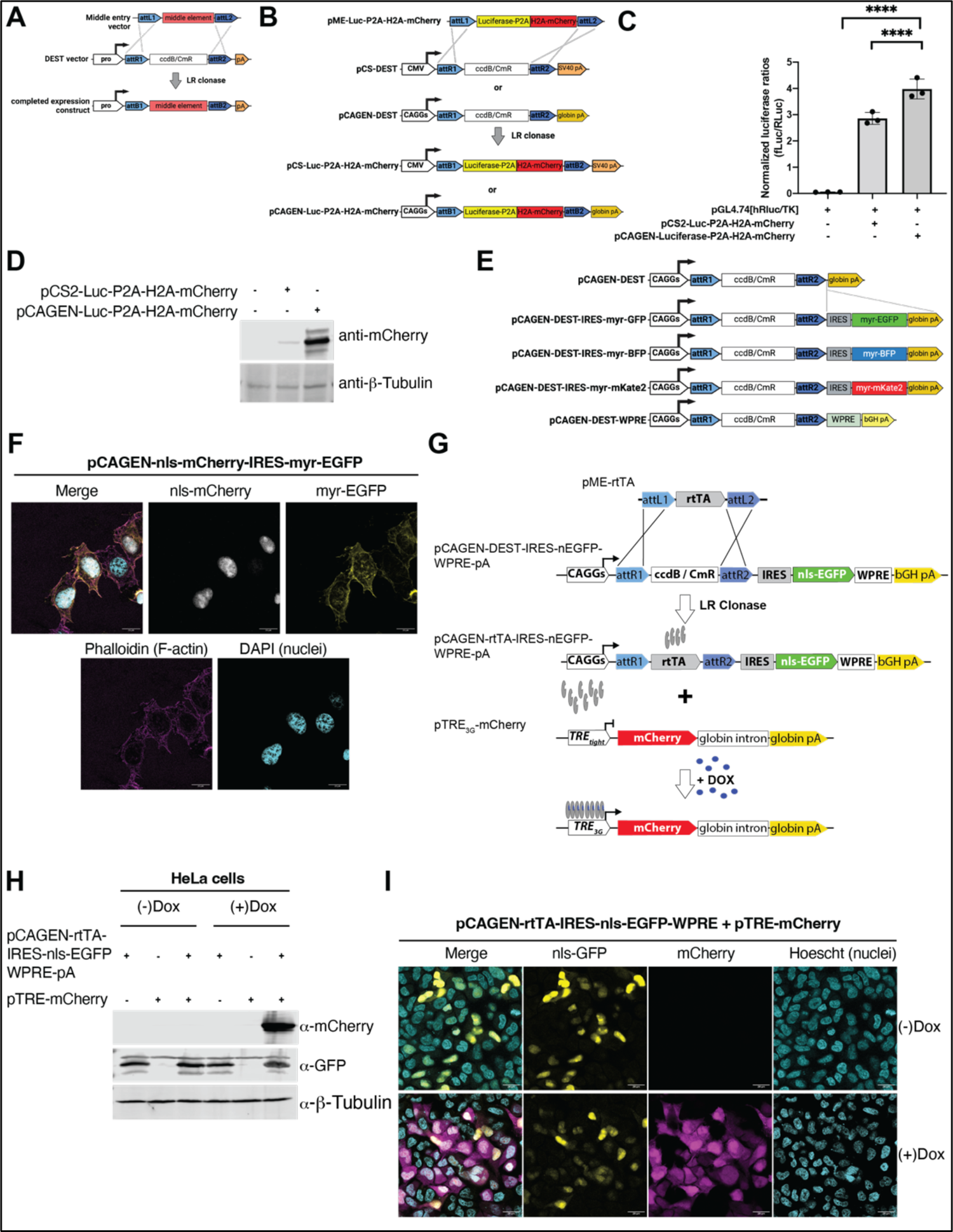
pCAGEN-Based Destination Vectors. **(A)** Schematic for pME-compatible, Gateway Destination mammalian expression vectors. (**B**) Previous pCS2-backbone based vector (pCSDest) developed by the Lawson laboratory (see Villefranc et al., Dev Dyn, 2007) which utilizes a CMV promoter with the CMV immediate early enhancer cassette and the SV40 polyA sequence, compared with the new pCAGEN-Dest vector which utilizes the synthetic CAG promoter and B-globin polyA from pCAGGs (see Matsuda and Cepko, PNAS, 2004). Example of an LR reaction used to generate pCS-Luciferase-P2A-H2A-mCherry and pCAGEN-Luciferase-P2A-H2A-mCherry. (**C**) Luciferase assays confirm greater transcriptional activity using the pCAGEN-Dest backbone than pCSDest. (**D**) Western blots and densitometry for mCherry protein confirm the luciferase analysis. (**E**) A suite of other dual reporter (IRES-fluorescent reporter backbones) pCAGEN-DEST vectors, as well as one with a WPRE element for increased transcript stability. (**F**) Confocal images of pCAGEN-nls-mCherry-IRES-myr-EGFP show dual fluorescence activity. (**G**) Schematic for pME-rtTA recombination with pCAGEN-DEST-IRES-nls-EGFP-WPRE-pA and co-transfection with a pTRE-mCherry reporter. (**H**) Western blot for detection of mCherry, GFP, and a loading control (β-Tubulin) following transfection in HeLa cells of Tet-On pCAGEN-rtTA-IRES-nls-EGFP-WPRE-pA and a pTRE-mCherry reporter confirm induction of mCherry in the presence but not absence of Dox. (**I**) Epifluorescent imaging following transfection of the indicated construct in the presence and absence of Dox confirm selective induction of the TRE-mCherry reporter.

To verify the functionality of this pCAGEN Destination vector, we generated a middle Entry clone where a strong consensus Kozak sequence precedes luciferase, followed by a P2A cleavage peptide, then an H2A-mCherry fusion protein (pME-Luc-P2A-H2A-mCherry, JDW 926). This Luc-P2A-H2A-mCherry insert was shuttled from the middle Entry vector by LR recombination into pCS2-Dest, pCAGEN-Dest, and pCAGEN-WPRE-Dest (Figure 6B). The transfer of the Luc-P2A-H2A-mCherry insert into each Destination vector was greater than 85-90% efficient (data not shown). To confirm the inserts and Destination vectors were functional, we transiently transfected the resulting plasmids into HEK-293 cells and processed them for either luciferase analysis, western blotting, or immunofluorescence. Luciferase activity shows that an increase in activity in the pCAGGS backbone compared to pCS2 (Figure 6C), which is confirmed by western blot (Figure 6D). Because stabilizing mRNA production may be desirable in some experimental settings, we replaced the original rabbit β-globin polyA in pCAGEN-DEST with a WPRE followed by a polyA from bovine growth hormone (bGH) (pCAGEN-WPRE-bGH-DEST, JDW 1309) (Figure 6E).

Given that even something as innocuous as a few amino acids from a P2A peptide may alter a protein’s function, we constructed a parallel set of bicistronic pCAGEN vectors in which a Gateway cassette lies upstream of an IRES followed by a fluorescent reporter. In this context, the CAG promoter drives transcription of both the mRNA transcript of interest (shuttled into pCAGEN via an LR reaction with a pME vector), as well as the downstream fluorescent reporter within the Destination plasmid backbone. However, whereas translation initiates in the first cistron at the 5’ capped mRNA for the protein of interest, translation of the second cistron occurs in a cap-independent manner mediated by the IRES. We have generated pCAGEN-IRES-myristoylated (myr) BFP, myr-mKate, myr-EGFP Destination plasmids (**Table 5**) (Figure 6E). N-myristoylation, a co-translational event in which myristate (a 14-carbon fatty acid) is covalently attached to an N-terminal glycine within this 18 amino acid peptide tag, drives subcellular localization of proteins to the plasma membrane, enabling fluorescent labelling of the cell surface ^125^.

**Table 5.**
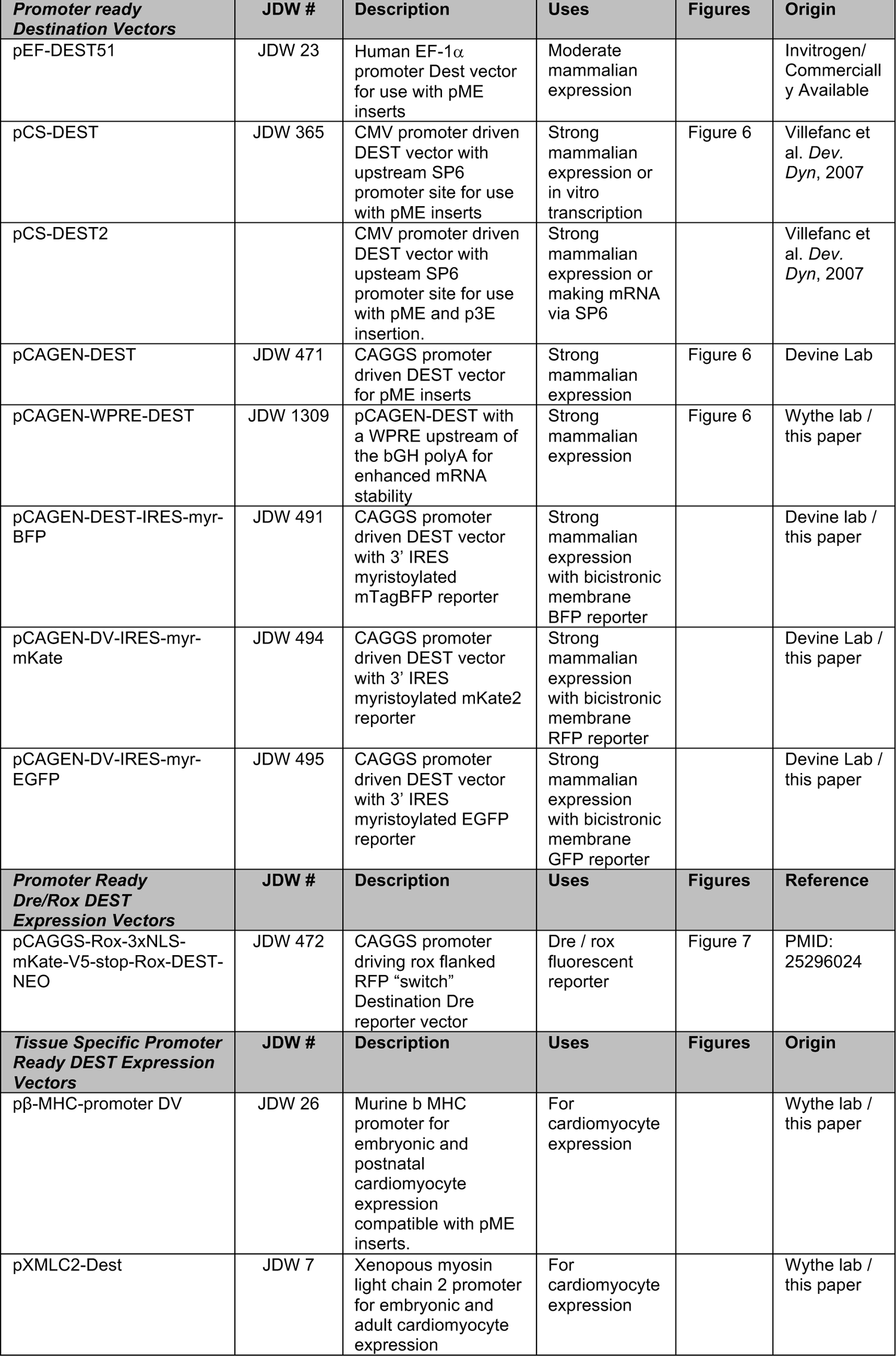
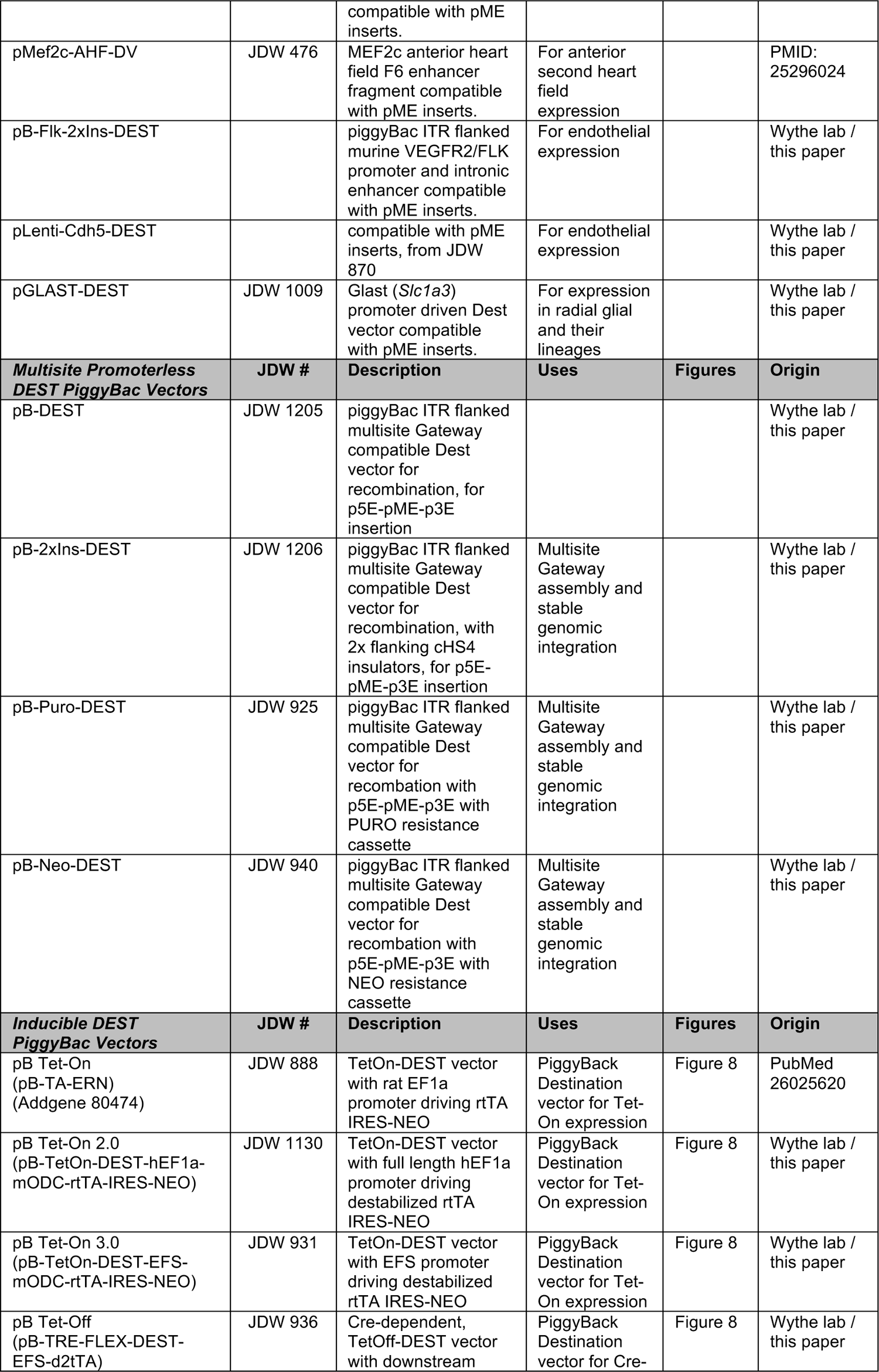
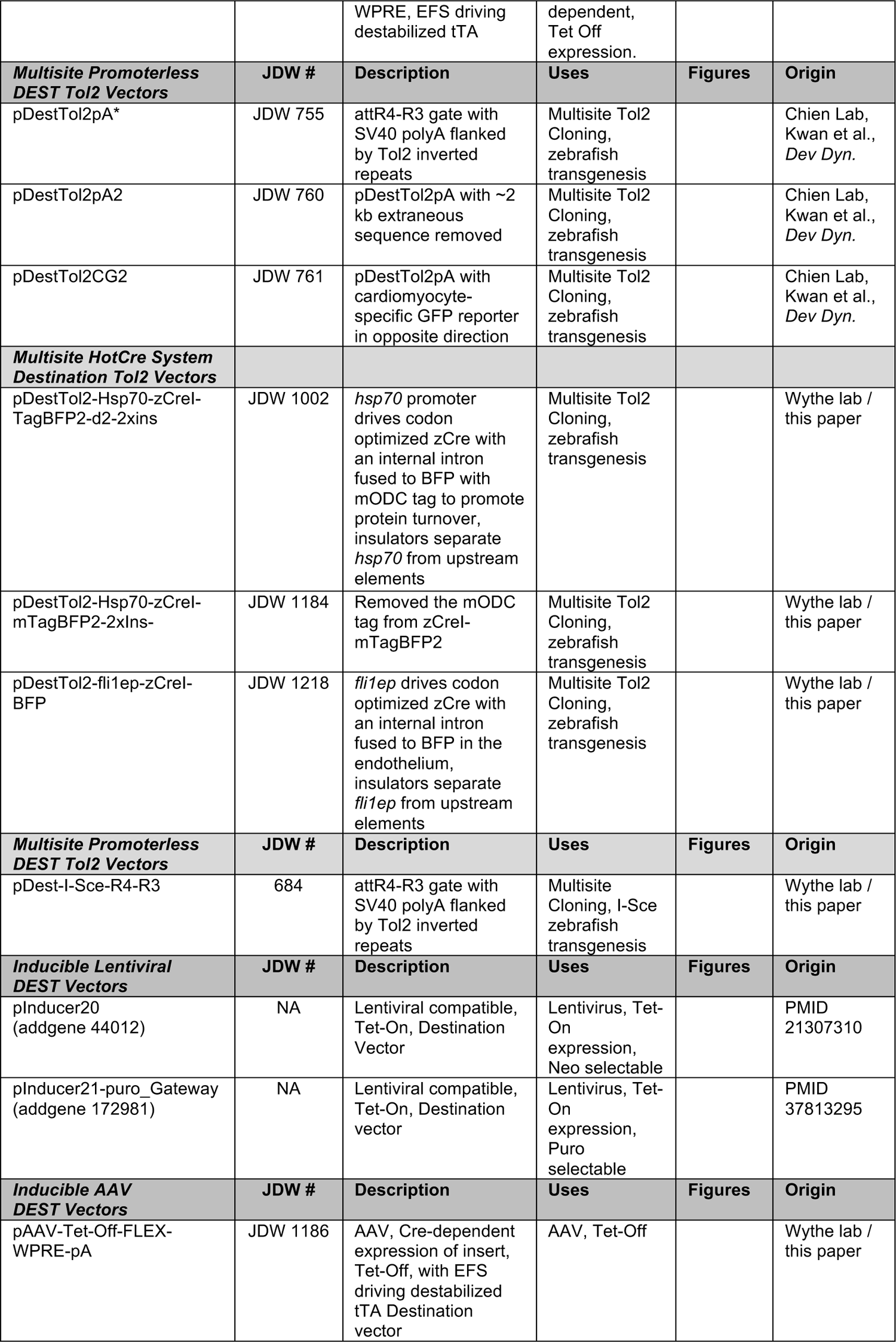
Existing and Novel Destination Vectors.

The utility of the other myristoylated IRES vectors was first confirmed by recombination of a middle Entry clone containing 3x-NLS-mCherry, as co-transfection yield clear mCherry localization to the nucleus and EGFP was confined to the cell membrane, as confirmed by counter-staining with DAPI and phalloidin to label the nucleus and F-actin, respectively (Figure 6E**,F**). We also validated the IRES-myristoylated FP DEST vectors by recombining a 3^rd^ generation rtTA transcriptional transactivator (pME-rtTA3 (Tet-On), JDW 1230) with pCAGEN-IRES-nls-EGFP-WPRE-DEST to generate pCAGEN-rtTA-IRES-nls-EGFP-WPRE-bGH-pA (JDW 410). Subsequent co-transfection with a TRE-mCherry reporter plasmid and immunofluorescence confirmed robust co-expression of the IRES reporter and the tetracycline-inducible mCherry reporter only in the presence of Doxycycline (Figure 6G-I).

**Table 6.**
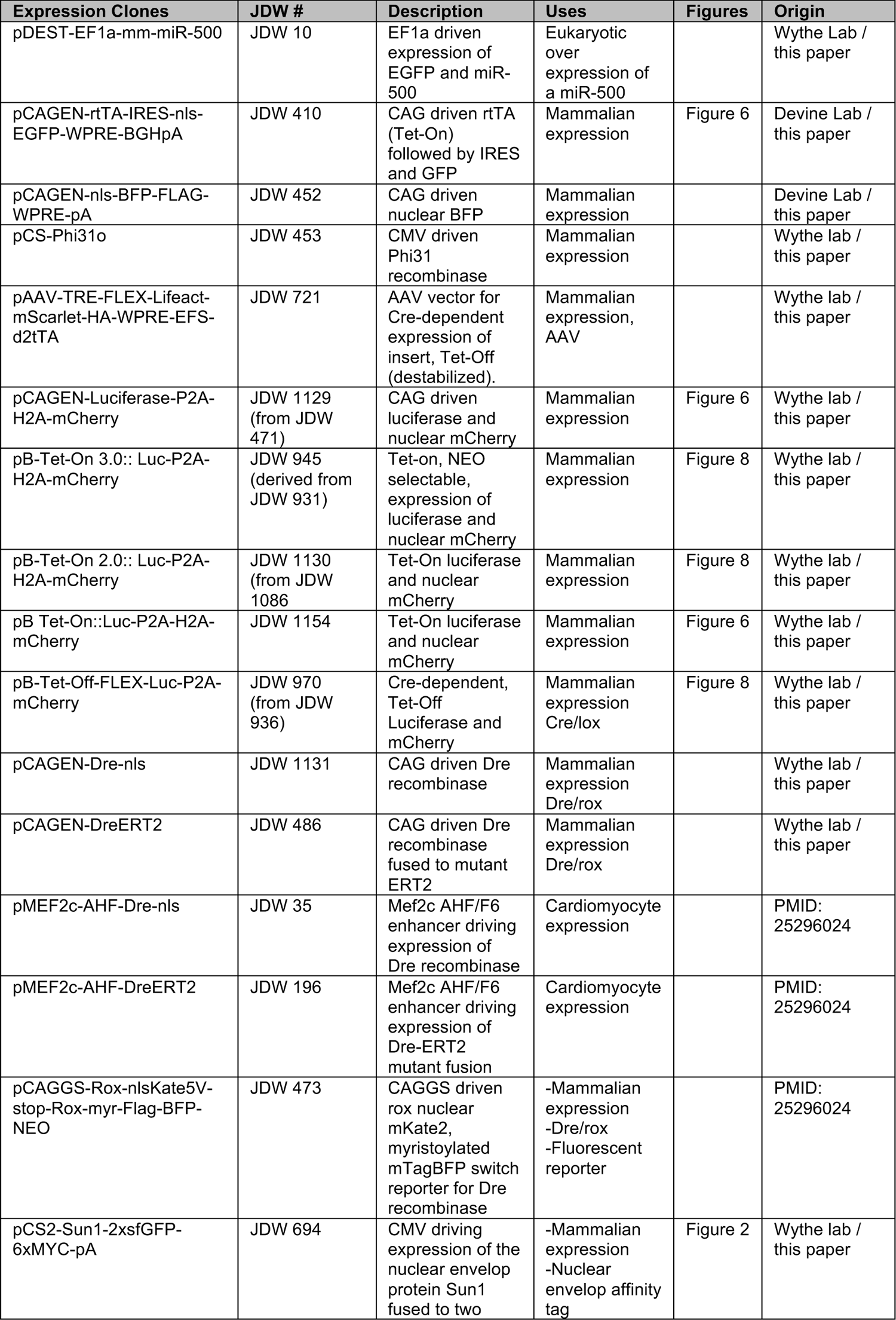
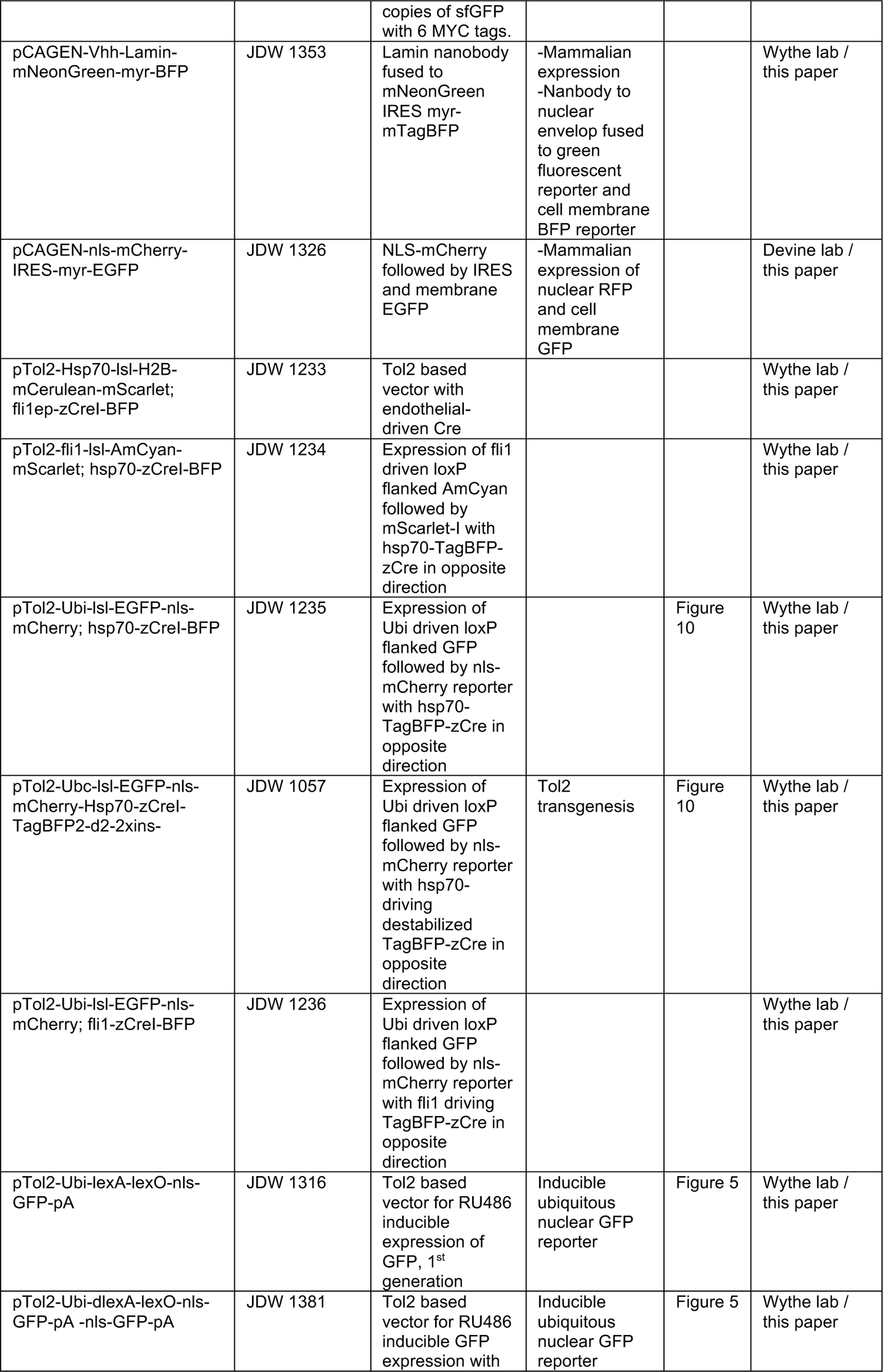
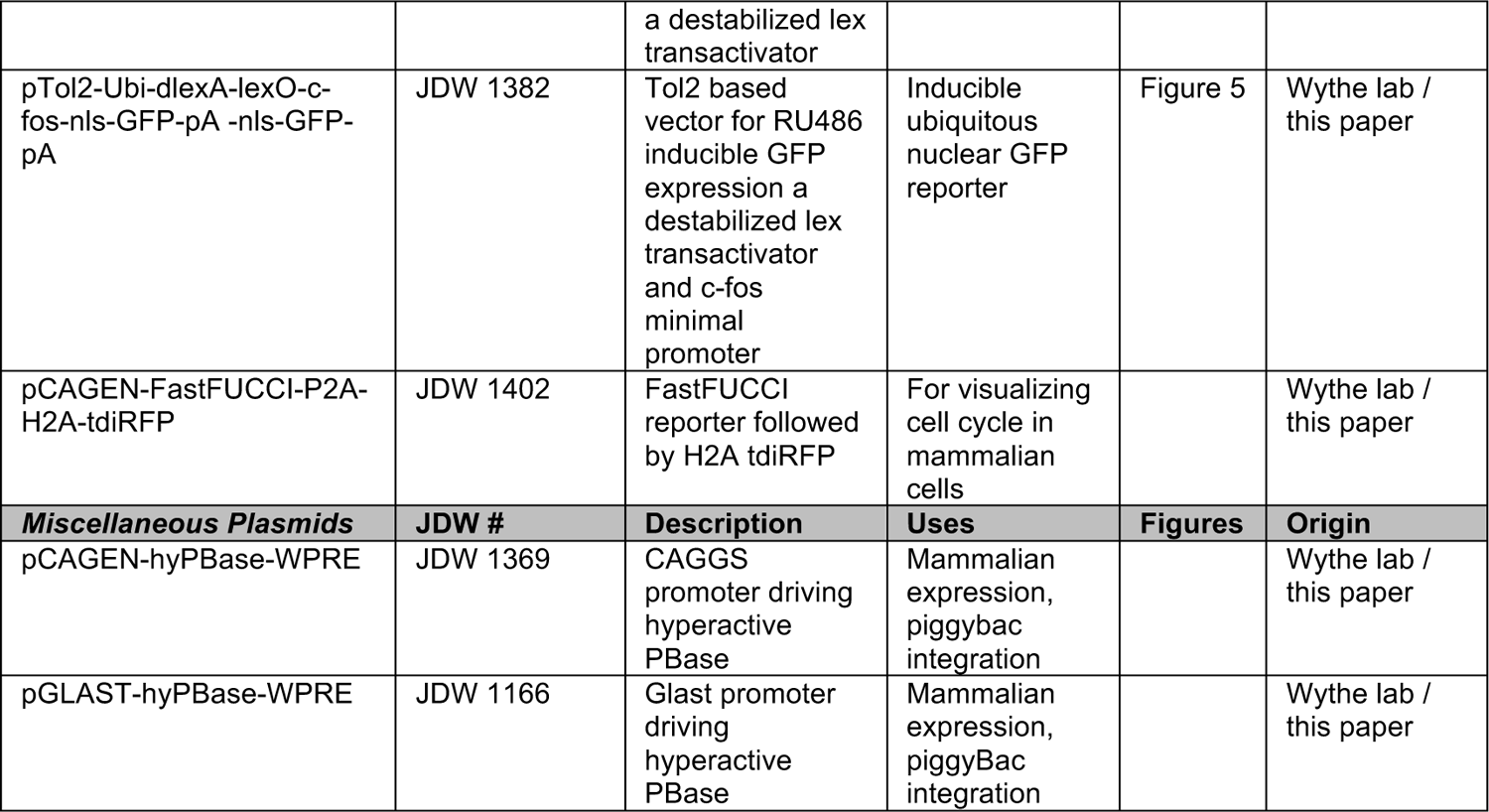
Novel Expression Clones.

To confirm the utility of our previously mentioned Dre recombinase containing pME clones, we also generated a Dre recombinase reporter vector wherein a nuclear localized, V5 epitope tagged RFP variant, mKate (nls-mKate-V5), followed by a rabbit β-globin polyA sequence, was placed between two ROX recombination sites, all downstream of a CAG promoter. A Gateway RFA Destination cassette (including a rabbit globin polyA sequence) was subsequently added downstream to generate pCAGGS-rox-3xnls-mKate-V5-stop-rox-Dest (pCAG-rsr-nK-Dest). A SacI-SalI fragment containing an FRT flanked PGK-NEO cassette was cloned downstream of the Gateway RFA-rabbit globin polyA sequence in the opposite direction to enable selection for stable clones via neomycin resistance (pCAGGS-rsr-nK-DEST-FRT-PGK-NEO-FRT / JDW 472). As a proof of concept, the latter vector was recombined with a pME-myr-TagBFP2-FLAG (JDW 1183) to generate a rox-dependent red to blue “switch” vector: pCAGGS-rox-3xNLS-mKate2-V5-stop-rox-myr-TagBFP2-FLAG-FRT-pgk-NEO-FRT or pCAGG-rsr-nKmB (JDW 1217) (Figure 7A). Transient transfection in HEK-293 cells revealed efficient recombination, as evidenced by switching of nuclear mKate signal for membrane localized BFP upon co-transection with Dre recombinase (Figure 7B). The entire CAG-rsr-nKmB cassette can be removed via AscI to PacI for insertion into murine gene targeting vectors. We previously validate the functionality of this cassette at the *Rosa26* locus in the embryonic mouse^103^. As an alternative to the prevalently used *Rosa26* locus, we also provide a Dest vector targeting the gene desert region in the *Hipp11* locus (JDW 478) on chromosome 11 between *Eif4enif1* and *Drg1* ^126^. Recombination of this *rox* switch reporter cassette into this Dest vector generated *pHipp11*-Rox-nls-Kate-V5-Rox-myr-TagBFP-FRT-NEO-FRT-DTA to allow for gene targeting of this same construct to the *Hipp11* locus^126^. presence and absence of Dre recombinase confirms a clear gain of mNeon-Green signal within the actin cytoskeleton in the presence, but not absence, of Dre recombinase, unlike cells that did not receive pCAGEN-Dre (Figure 7C**,D**). Due to the presumed stability of the 3xNLS-mCherry reporter, there were some cells that had simultaneous mCherry signal in the nucleus and mNeonGreen labelling of actin.

**Figure 7:**
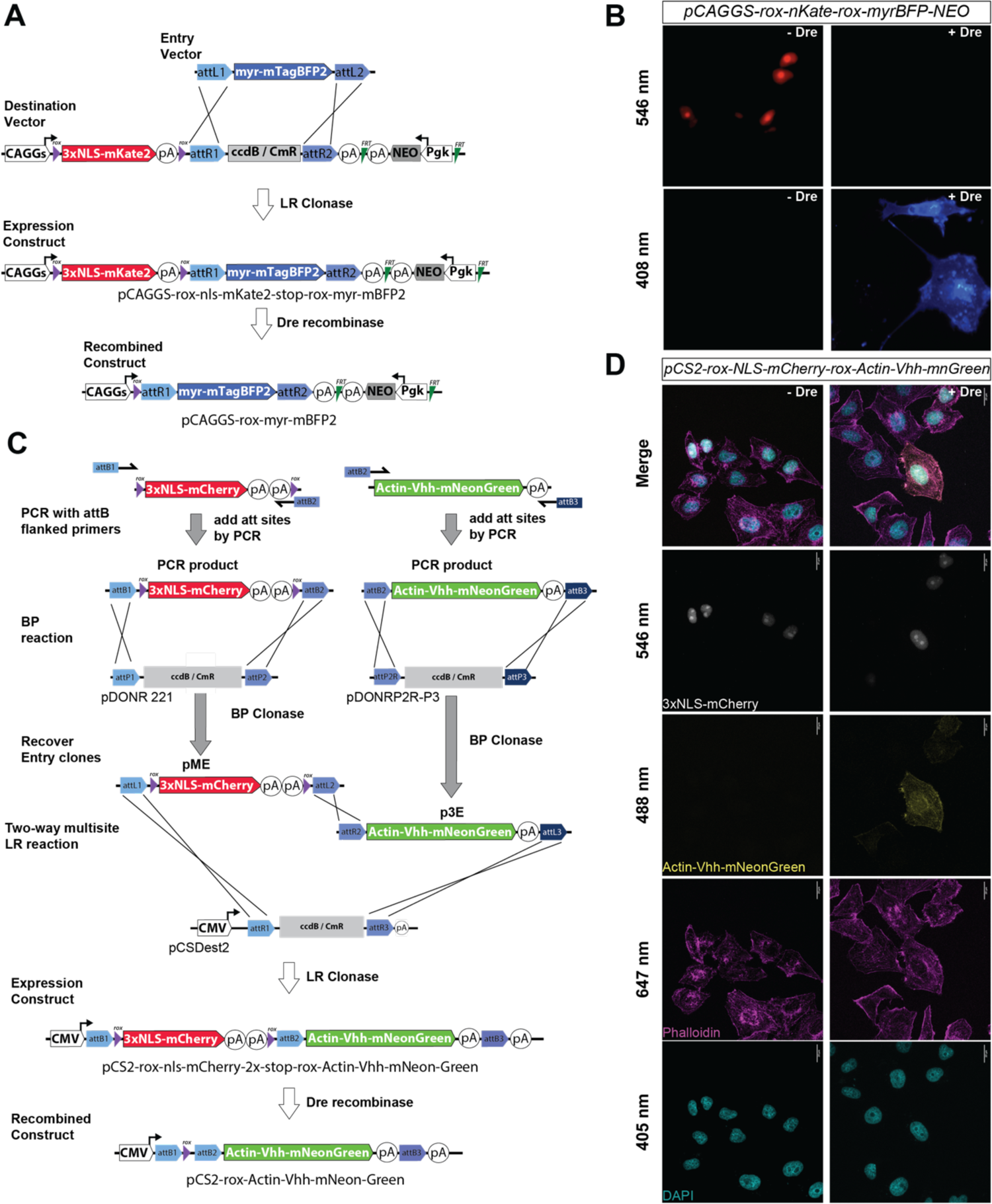
The Dre/Rox Recombinase System. **(A)** Schematic for a pME-compatible Gateway Destination vector where a CAG promoter drives expression of a rox flanked nuclear mKate2 reporter, and a pME cassette, such as one containing myristoylated mTagBFP2, can be inserted downstream. Upon rox mediated recombination, mKate2 and the transcriptional stop cassette are removed, and the downstream myr-mTagBFP2 cassette is expressed. (**B**) Validation of the pCAG-nuclear-Kate-myr-BFP (nKmB) reporter following co-transfection in HEK-293 cells with and without Dre recombinase. (**C**) Generation of a novel dual Dre/rox reporter, where 3xNLS mCherry and a 2x stop cassette are flanked by rox recombination sites in a pME vector, and a novel p3E construct containing an Actin nanobody (Vhh) fused to mNeonGreen are inserted distally into a pCSDDest2 vector. (**D**) Validation of the switch reporter activity in the presence and absence of Dre recombinase.

### Tissue specific Destination vectors for single Entry cloning

In some settings, ubiquitous expression of a gene of interest may lead to unwanted developmental defects due to the lack of spatial and/or temporal control over expression. To accommodate such instances, we provide Destination vectors compatible with pME Entry clones to drive expression of an ORF in a particular cell type and/or at a specific time point in development. We have generated one vector for broad, embryonic cardiomyocyte-restricted expression that leverages the murine *β myosin heavy chain* (*MYH7*) promoter, pβ-MyHC-promoter Dest (JDW 26)^127^, as well as a plasmid where cardiomyocyte expression is driven by the *Xenopous laevis Myosin Light Chain 2* promoter, pxMLC2-pro-Dest (JDW 7). We have also generated and validated a Dest vector that drives expression within the anterior second heart field progenitors of the early mouse embryo via the MEF2c-AHF enhancer (also known as F6/frag3) (pMEF2c-AHF-Dest)^103^.

We have also created two DESTINATION vectors where expression in driven by an endothelial-specific promoter. One of these is in a lentiviral backbone where expression is driven by the murine *Cdh5* (VE-Cadherin) promoter, allowing for viral transduction and expression within endothelial cells (pLenti-Cdh5-DEST). For non-lentiviral experiments, we placed the murine *Vegfr2* (*Kdr*/*Flk*) promoter and downstream intronic enhancer^128^ between two flanking *piggyBac* ITRs and a pair of core insulator elements to prevent positional effects (pB-Vegfr2-pro-WPRE-Dest). Finally, we also provide a Destination plasmid containing the 2.1 kb human *GLAST* (*EAAT-1*) promoter for driving expression within the radial glia lineages: cortical neurons, astrocytes, oligodendrocytes, and olfactory bulb interneurons^60,61^. Collectively, these tools should expand the toolkit available to researchers interested in both in vitro and in vivo neuronal and cardiovascular research.

### Tet-regulated Destination vectors for single Entry cloning

While tissue specific expression can be desirable, in some instances the ability to rapidly induce, or silence, gene expression can be quite desirable either in vitro or in vivo. Many Doxycycline-inducible (e.g. “Tet-On”) expression plasmids compatible with mammalian systems exist, such as the lentiviral pInducer suite ^129^, the adeno-associated viral vector pRAM ^130^, or the *piggyBac* vector pB-TA-ERN ^33^. However, in our experience some of these tools have “leaky” activity in the absence of Doxycycline (data not shown), likely due to excessive accumulation of the reverse tetracycline transactivator and non-specific activation of the TRE promoter element. To mitigate this non-specific activity, we focused on improving the Tet-On, *piggyBac* compatible pB-TA-ERN vector (JDW 888, addgene # 80474), which also contains a PGK-NEO cassette for stable selection after integration into mammalian cells (Figure 8A). We first replaced the potent CMV-IE enhancer and the full-length rat EF1a promoter with a transcriptional pause site and a full length human *EF1a* promoter (*hEF1a*) cassette. Next, we added an amino terminal PEST degradation domain from murine *Ornithine Decarboxylase* (mODC)^105^ to rtTA to reduce the half-life of the transactivator (pB_TetOn_DEST_hEF1a_mODC_rtTA_IRES_ NEO, or pB Tet-On 2.0) (JDW 1086). Using this substrate, we then created another variant where the full length human *EF1a* promoter cassette was replaced with a short, intron-less form of the human *EF1a* promoter (*EFS*, also known as *EF1a short*, or *EF1a core*)^29^ to generate pB-TetOn-DEST-EFS-mODC-rtTA-IRES-NEO, or pB Tet-On 3.0 (JDW 931). To determine whether these alterations in rtTA stability and the promoter affected the ‘on/off’ kinetics of the system, we recombined each of them with an ENTRY vector encoding for Luciferase and nuclear localized mCherry (pME-Luc-P2A-H2A-mCherry) (JDW 926).

**Figure 8:**
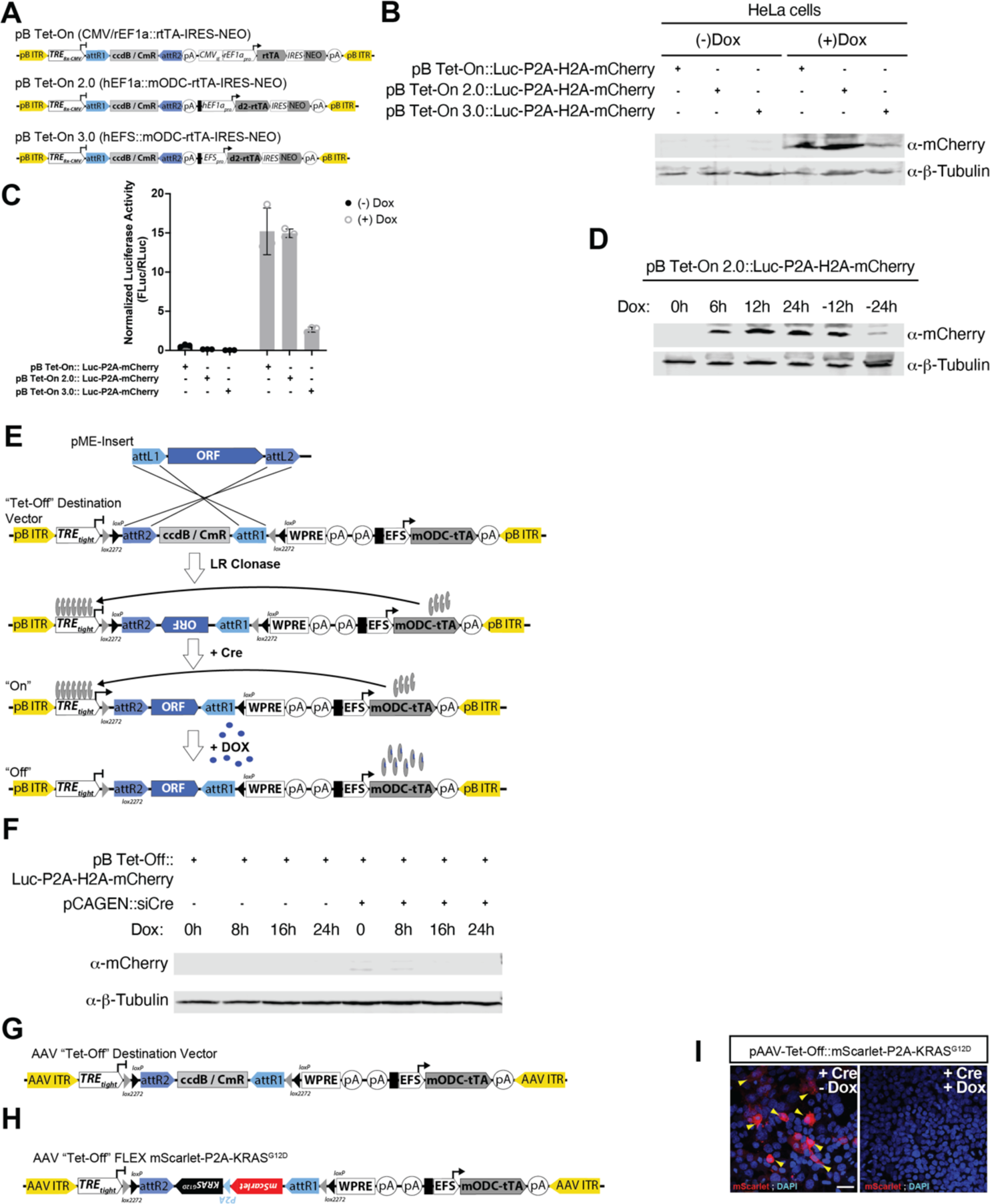
A suite of Tet-On and Off Gateway Vectors for Eukaryotic Studies. (**A**) Three variants of Tet-On vectors, with modifications to the promoters driving rtTA expression (CMVIE and rat EF1a versus human EF1a alone versus an EF1a short or EFS minimal promoter), along with regular rtTA versus a destabilized rtTA (d2rtTA) to prevent accumulation of the transcriptional activator. (**B**) Western blot for mCherry and a loading control, β-tubulin, confirm varying levels of Dox induction for each of the three variants following transient transfection in HeLa cells. (**C**) luciferase reporter activity following transient transfection in HeLa cells. (**D**) Kinetics of Luciferase-P2A-H2A-mCherry expression shown by Western blot following addition and withdrawal of Dox. (**E**) A binary, Cre-dependent, Tet-Off Destination vector design. (**F**) Validation and kinetics in HeLa cells. (**G**) A similar AAV-based FLEX or DIO approach for Cre-dependent, Tet Off expression and an (**H**) example vector, AAV-Tet-Off-FLEX-mScarlet-P2A-KRASG12D and (**I**) validation of Cre-dependent expression and Dox-induced silencing of transcription following transient transfection in HEK-293s.

First, we confirmed varying levels of mCherry induction following transient transfection in HeLa cells and addition of Doxycycline by western blot (Figure 8B). Quantification of luciferase activity post-transient transfection showed that pB-Tet-On and pB-Tet-On 2.0 had similar fold inductions, although pB-Tet-On 2.0 had less variation, while pB-Tet-On 3.0 was less robust, in agreement with the western blot data (Figure 8C). Notably, basal induction of pB-Tet-On 2.0 and 3.0 was substantially less than the original pB-Tet-On. To verify that the mODC tag and removal of the CMV enhancer resulted in more sensitive induction, following transient transfection cell lysates were collected 0, 6, 12, and 24 hours after Dox addition, as well as 12 and 24 hours after withdrawal of Dox. This newer variant showed little to no induction prior to addition of Dox, and substantial reduction of reporter activity within 24 hours of Dox withdrawal, confirming that these newer variants exhibit better on/off kinetics than previous designs (Figure 8D).

To complement these tools, we engineered a *piggyBac* compatible, Tet-Off vector where expression of the insert requires Cre-mediated recombination (pB-Tet-Off-FLEX-EFS-mODC-tTA) (JDW 936). In this variant, the TRE-tight promoter (composed of 7x, Tet operons and a minimal CMV promoter) is upstream of a reverse oriented Destination cassette (*attR2*-*ccd*B-Cm^R^-*attR1*, rather than *attR1-ccd*B-Cm^R^-*attR2*) that is flanked by <u>d</u>ouble inverse <u>o</u>riented *loxP* and *lox2272* recombination sites (DIO, or FLEX), followed by a WPRE and bGH polyA, with a human EFS promoter driving expression of tTA followed by the SV40 late polyA (pB_Tet-Off-FLEX-EFS-d2tTA) (JDW 936) (Figure 8E). This Destination vector was recombined with pME-Luc-P2A-H2A-mCherry (JDW 926) and the resulting plasmid (pB-Tet-Off-FLEX-Luc-P2A-H2A-mCherry, JDW 970) co-transfected with and without Cre. Western blot confirmed H2A-mCherry was only detected in the presence of Cre. However, this expression was undetectable 16 hours post addition of Doxycycline (Figure 8F). We extended this combinatorial approach from the piggyBac system to generate an adenovirus associated vector (AAV) compatible, Tet-Off vector wherein TRE_tight_ is upstream of a Destination cassette flanked by a pair of double inverse-oriented *lox* sites, a WPRE and bGH polyA, followed by a transcriptional pause site and minimal EFS promoter driving expression of a destabilized tTA (pAAV-Tet-Off-FLEX-Dest) (JDW 1186) (Figure 8G). As expected, expression of an mScarlet-P2A-KRAS^G12D^ insert was evident in the presence of Cre recombinase, but undetectable following addition of Doxycycline (Figure 8H**,I**).

### PiggyBac Destination vectors for multisite Gateway cloning

To maximize the flexibility of the MAGIC kit in mammalian systems, we constructed *piggyBac* compatible multisite Gateway plasmids in a manner analogous to the zebrafish *Tol2* multisite Gateway system to facilitate transgenesis with integration of the cargo plasmid into the genome (Figure 9A). In this design, three Entry clones (p5E, pME, and p3E) integrate in the 5’ to 3’ direction into a single *piggyBac* ITR flanked Destination vector (Figure 9B). Two of these vectors contain a downstream SV40 promoter driving expression of a selectable marker (either puromycin or neomycin) to allow for the generation of stable lines (pB-Dest-NEO, JDW 940; pB-Dest-PURO, JDW 925) (Figure 9C). A third compact variant lacks the stable selection cassette (pB-Dest, JDW 1205), while a final variant has 5’ and 3’ flanking pairs of the 250-bp core <u>c</u>hicken <u>H</u>ypersensitive <u>S</u>ite-<u>4</u> (cHS4) insulator from *β-globin* (pB-2xIns-Dest, JDW 1206) to protect integrated transgenes against heterochromatin propagation and silencing by the recruitment of histone acetyltransferases and methyltransferases^131–134^.

**Figure 9.**
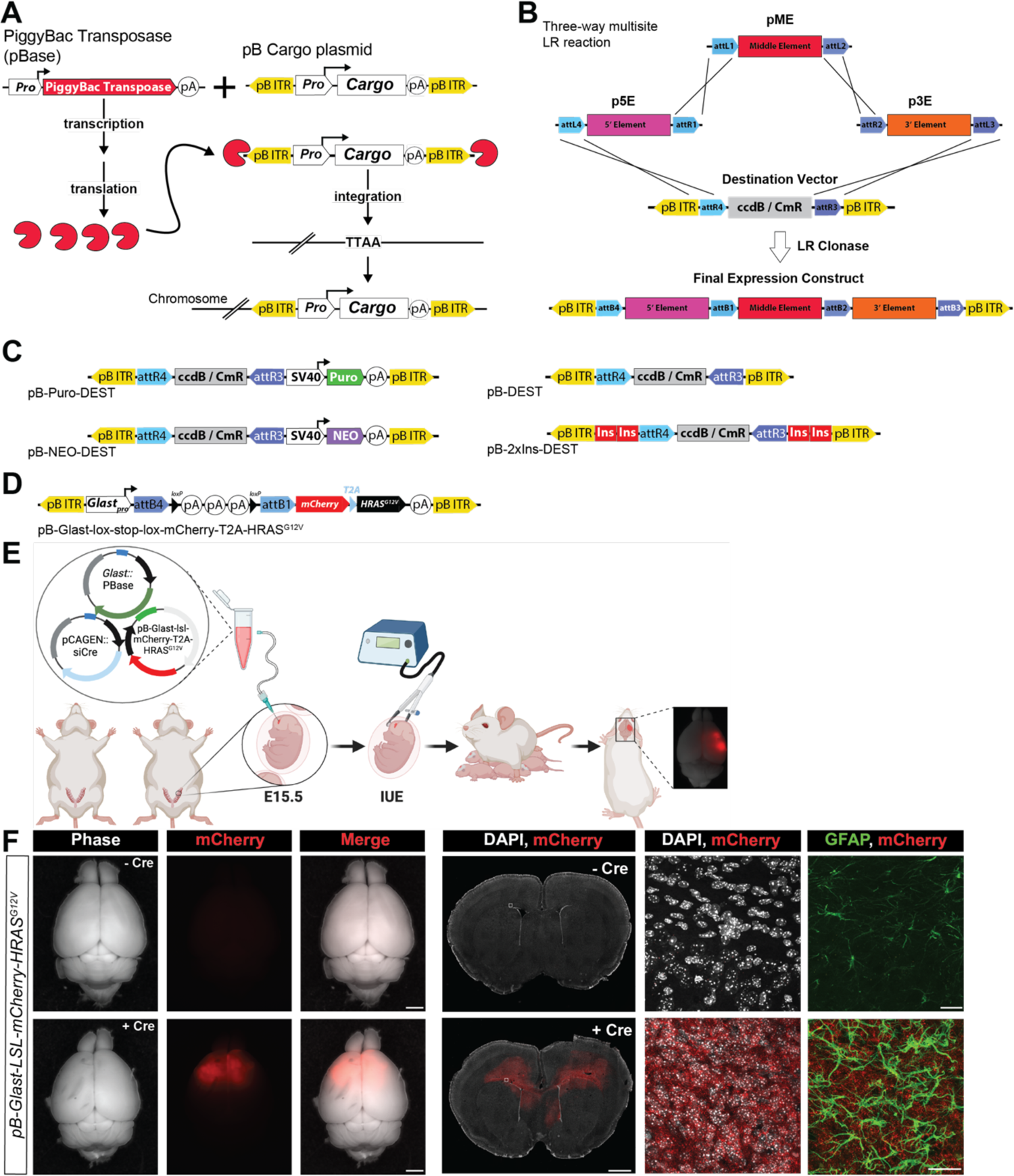
PiggyBac Gateway Compatible Vectors in the M.A.G.I.C. System. **(A)** In the PiggyBac binary system, piggyBac transposase promotes integration of a pB ITR flanked cargo cassette into the genome at AATT sequences. (**B**) A schematic of the three-way LR Destination scheme for Gateway cloning using a piggybac ITR flanked Destination vector. (**C**) A suite of selectable and non-selectable pB ITR flanked Destination plasmids for multisite Gateway cloning. (**D**) An example plasmid, *pGlast-3xSTOP-mCherry-T2A-HRASG12V*. (**E**) Schematic for in utero electroporation in mice followed by visualization of mCherry-positive, HRAS-driven glioblastoma in mice. (**F**) Wholemount phase and epifluorescence images from representative brains (far left), scale bar = 2 mm. Far right images are confocal images of frozen cryosections stained for mCherry (tumor), GFAP (reactive astrocytes), and DAPI (nuclei). Scale bar = 20 µm.

To illustrate the utility of these reagents, we performed a three-way multisite LR reaction using p5E-*Glastpro*, pME-*loxP*-3xStop-*loxP*, and p3E-mCherry-T2A-HRAS^G12V^-SV40-polyA into the pB-2xIns-Dest backbone to generate a cargo plasmid where *GLAST* promoter driven expression of oncogenic HRAS^G12V^ within radial glia and astrocytes is dependent upon Cre-mediated excision of an upstream *loxP* flanked stop cassette (Figure 9D). In utero electroporation (IUE) of this pB-2xIns-*Glast*-*loxP*-3x-Stop-*loxP*-p3E-mCherry-T2A-HRAS^G12V^ *piggyBac* transposon flanked cargo plasmid along with a construct where *GLAST* drives expression of a hyperactive *piggyBac* transposase (hyPBase)^135^, together with a self-excising Cre plasmid (pCAGEN-lox-Cre-stop-lox), resulted in stable transgene expression in radial glia descendants (cortical neurons, astrocytes, oligodendrocytes), and yielded frank glioma, as evidenced by extensive upregulation of reactive astrocytes shown by immunohistochemistry for anti-Glial fibrillary acidic protein (GFAP) (Figure 9E**,F**) and neoplastic figures as evidenced by H&E staining (data not shown).

### Cre compatible, Tol2 Destination vectors for multisite Gateway cloning

The *Tol2* transposon element, originally identified in the medaka fish (*Oryzias latipes*), belongs to the hAT family of transposons (*hobo*, *Ac*, *Tam3*), which are flanked by inverted repeats and encode their own transposase^136,137^. Destination vectors containing flanking *Tol2* transposon elements needed for genomic transposition enable robust and efficient transgenesis in zebrafish, particularly when used in conjunction with *Tol2* transposase mRNA^138^ (**Figure 10A**). By combining a heat-shock promoter for temporal control of Cre recombinase expression, together with a tissue-specific promoter upstream of a floxed stop cassette, we created an “all-in-one” vector that obviates the need for crossing multiple lines, or injecting multiple plasmids, that is analogous to the previously described two component HOTcre system in zebrafish^139^. In our backbone the *Tol2* elements flank the multisite *attR4-ccd*B-*Cm^R^-attR3* Gateway cassette in the 5’-3’ direction, followed by an SV40 late polyA signal sequence and a pair of 250-bp chicken *β-globin* cHS4 insulators to prevent the interaction between the distal enhancer and the upstream promoter^132,140^ (**Figure 10B**). In the opposite direction, the 1.5 kb heat-shock inducible *hsp70* promoter^141^ drives transcription of *Cre.zf1* – a codon optimized variant for expression in zebrafish^142^ – fused to TagBFP and a c-terminal mODC peptide to reduce the half-life of Cre (**Figure 10C**). Cre recombinase and *loxP* sites should not be present in the same construct due to the risk of recombination during propagation in prokaryotes (i.e. bacteria). Thus, *Cre.zf1* was modified to include an artificial intron separating the 5’ and 3’ halves of the open reading frame (*CreI.zf1,* or *zCreI*), ensuring that Cre expression is restricted to eukaryotic cells^143^. Another variant of this vector was created wherein the mODC peptide was removed, *pTol2-Dest; hsp70-zCreI.zf1-BFP* (JDW 1184), to allow for accumulation of zCreI-TagBFP and potentially more robust recombination frequency. Finally, we also created a modified, tissue-specific version of this 2^nd^ generation HOTcre “all in one” plasmid that targets the endothelium (pTol2Destp2a-2xIns; fli1ep-zCreI-BFPn, JDW 1218). In this DEST vector, the zebrafish *fli1a* enhancer/promoter (*fli1ep*) drives endothelial-specific expression of nls-Cre-mTagBFP-nls, allowing one to insert a p5E promoter of interest upstream of a pME lox-stop-lox cassette, followed by a p3E plasmid containing the insert of interest. As a proof of concept, we performed a three-way multisite Gateway reaction between a p5E plasmid containing a *Ubiquitin* promoter driving a *loxP* flanked EGFP reporter (p5E-*Ubi*-*loxP*-EGFP-*loxP*), pME-nls-mCherry, p3E-polyA and this Destination vector yielded p5E-*Ubi*-*loxP*-EGFP-*loxP*-mCherry;*hsp70*-CreI.zf1-BFP-mODC (JDW 1236) (**Figure 10D**). Injection of this construct and transposase mRNA into 1-cell stage zebrafish embryos and heat shock 24 hours later switched expression from a cytosolic EGFP to nuclear mCherry by 48 hours later (72 hpf), thus establishing the utility of this “all-in-one” vector for spatially and temporally controlled gene expression (**Figure 10E**).

**Figure 10:**
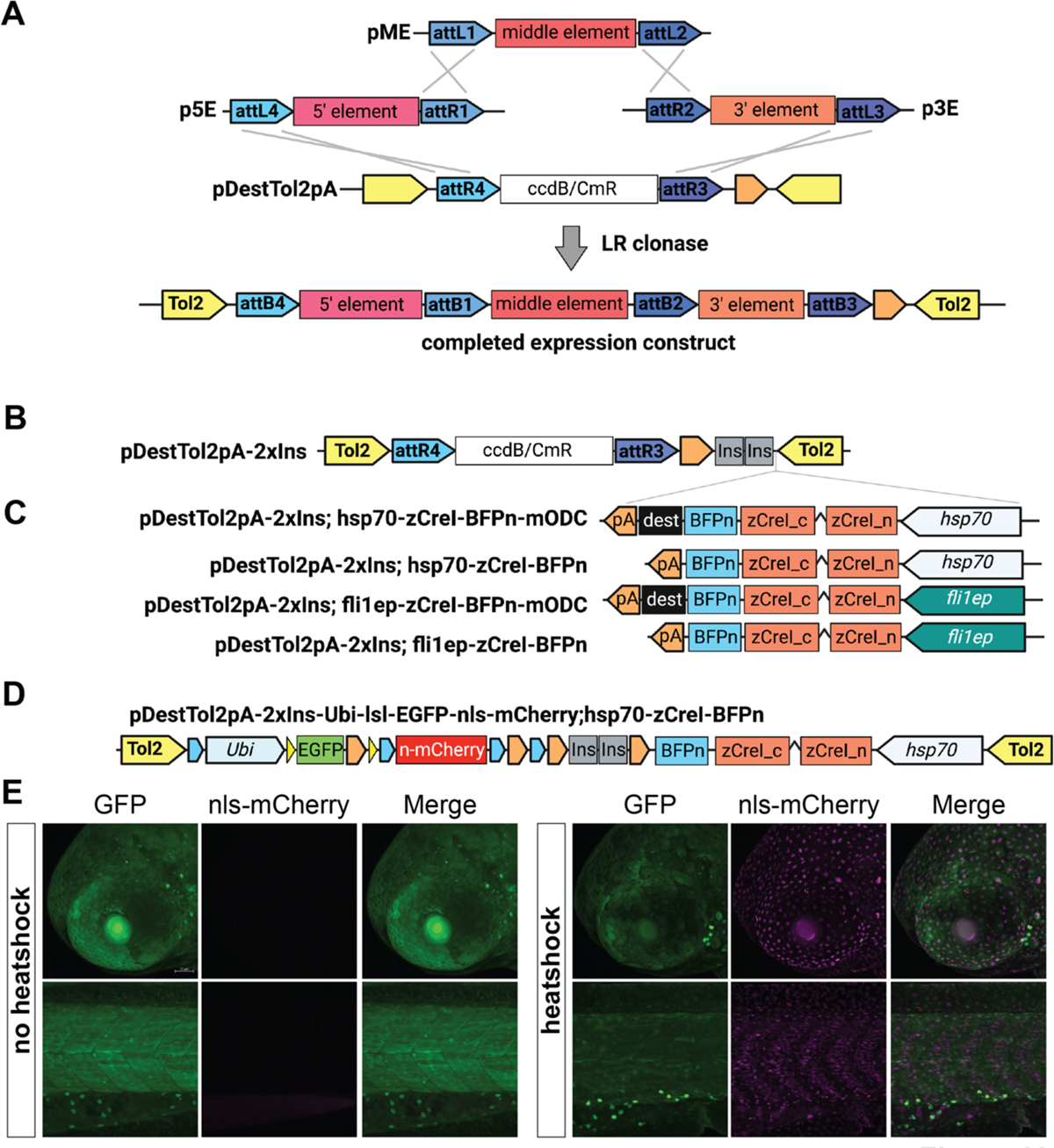
New All-in-One Cre Gateway Tools for Zebrafish. **(A)** The typical three-way LR recombination reaction. (**B**) A novel, insulator flanked Destination vector and (**C**) new derivative backbones where heatshock promoters (Hsp70) or endothelial-specific promoters (*fli1ep*) drive expression of codon optimized zCre separated by an internal intron to prevent spurious expression and resultant loxP recombination in e. coli during routine cloning steps. (**D**) An example vector, *pTol2-2xIns-Ubi-lox-EGFP-stop-lox-NLS-mCherry; Hsp70-zCreI-BFP-NLS*. (**E**) Example from a stable line, *Tg(Ubi-lsl-EGFP-NLS-mCherry;Hsp70-zCreI-BFP)* following either room temperature incubation, or heatshock, for 1 hour and then imaged at 72 hpf.

### I-Sce Destination vectors for multisite Gateway cloning

While *Tol2*-mediated transposition has been a mainstay for more than 25 years for generating transient and stable transgenic lines, the *I-SceI* meganuclease system is equally effective for generating transient and stable transgenic animals by circumventing the limited persistence of episomal plasmid DNA following multiple rounds of cell division and overcomes low germline transmission rates due to unequal integration of episomal DNA fragments into the host genome ^144–149^. *I-SceI*, an intron-mediated homing endonuclease isolated from yeast^150,151^, has an 18-bp recognition sequence expected to occur only once in 7×10^10^ bases of random sequence and is unlikely to cut genomic DNA. Accordingly, co-injection of the *I-SceI* meganuclease with a construct flanked at the 5’ and 3’ ends by the corresponding recognition sites leads to enhanced transgenesis compared to circular, supercoiled plasmid at rates around 30% for stable inheritance of the transgene at low copy number within a single genomic locus^152^. Thus, herein we supply I-SceI-Dest-pA (JDW 684), a plasmid where I-Sce sites flank an attR4-ccdB-CmR-attR3 cassette, followed by a poly A signal to allow for assembly of p5E, pME, and p3E inserts within the same backbone in a manner analogous to the Tol2 and piggyBac systems presented herein.

### A Relational Database for Plasmid Management

Due to the sheer volume of plasmids generated for this toolkit, and with the mandate for central lab organization and more robust data management from the National Institutes of Health, we developed a relational database using FileMaker software for maintaining electronic records of all plasmids and their salient features, including Pubmed identifiers, Addgene information, promoters, open reading frames, various uses of the plasmid, and other notes (Supplemental Figure 1). Critically, electronic files (such as those generated by APE or Snapgene) can be posted directly into database records, as can illustrations for maps of the plasmid (e.g. as .jpeg or .png files). All fields and container fields within the database are modifiable, as the package is presented as an open-source tool that can be easily modified by anyone familiar with FileMaker software. This database can be shared by an entire laboratory either on one centralized computer, or through multiple computers provided they each have access to FileMaker Pro software or that the database is hosted on a local computer or server (we host the database using a dedicated Mac Mini with its own IP address). Importantly, many of the data entry fields in this database correspond to those directly used by Addgene. Dropdown lists for container fields can be modified by any user, or entirely new lists or fields created, to provide a fixed vocabulary for categories such as plasmid use (for instance, zebrafish transgenesis, subcloning, mammalian expression, Gateway subcloning, etc.), and promoter type (e.g. CAGGS, EF1a, CMV, etc.). Other standard categories, such as whether there are bacterial or DNA stocks available, can be created and have yes/no dropdown tabs (Supplemental Figure 2).

Data from these various entry fields can be exported as a tab delimited file to excel to facilitate sharing or depositing plasmids at Addgene or other plasmid repositories, or to other laboratories, and individual PDFs of any record can also be generated directly in FileMaker and exported for sharing information about the plasmid of interest. The various entry fields we have created, such as plasmid use or promoter, can be modified by individual users (or removed entirely), as can the layout of the form, the size of data entry fields/containers, and indeed the entire underlying architecture of the system is open to user manipulation (Supplemental Figure 3, 4). A typical data entry workflow is included for reference (**Supplemental Movie 1**), as is a video showing navigation and search functionality (**Supplemental Movie 2**). Of note, the more detailed the description within the entry fields and containers, the better the search functionality will operate for finding plasmid records. An empty database, with one complete plasmid entry included for a reference, is provided (**Supplemental Data**) and is available on our laboratory website at https://www.wythelab.com/wythe-lab-databases.

Finally, all plasmids are being deposited at Addgene to enable rapid, easy distribution to the biological community. For updates, please see our lab webpage: https://www.addgene.org/Joshua_Wythe/

## DISCUSSION

The interchangeable and affordable genetic engineering capabilities enabled by multisite Gateway cloning, combined with the discovery of Tol2 transposon-mediated transgenesis, revolutionized and democratized the zebrafish field, enabling the rapid and easy generation of novel transgenic lines^3–5^. We hope these MAGIC toolkit reagents, which are compatible with existing Tol2 Gateway kits, will usher in a similar renaissance for transgenesis in other eukaryotic fields, particularly those that can utilize piggybac-mediated transgenesis, as well as lentiviral and AAV-based tools. Indeed, the utility of our toolset spans cultured mammalian cells, to in utero electroporation of murine and chicken embryos, to adult mice via AAV and lentiviral transduction, and, of course, zebrafish and other teleost species. By making each of these resources publicly available at Addgene, we hope to further disseminate these and other recent technologies, such as optogenetics and chemogenetics. Finally, by providing an open-source relational database for tracking and indexing all plasmids, we also hope that these tools and resources will be embraced by the bioscience community and this database will enable more accurate cataloguing of reagents, and the transfer of well characterized genetic tools between laboratories.

Herein we provide new vectors for robust, constitutive CAG promoter driven expression in cultured cells with our pCAGEN suite of Destination plasmids that are also compatible with middle entry clones from previous Tol2 toolkits. We go on to generate dual reporter systems using the IRES system for monitoring transfection efficiency, while also providing a catalogue of novel fluorescent reporters, including mCerulean, StayGold, mCardinal, mScarlet-I and mScarlet-I3, iRFP and tdRFP, mNeonGreen, among others. Furthermore, we provide tools for visualizing actin dynamics, organelles and specific subcellular compartments, as well as cell cycle status with FUCCI reporters. By providing a collection of various strength promoters, as well as Destination vectors, combined with a host of novel middle entry fluorescent reporters, this toolset will facilitate new in vitro and in vivo studies of subcellular dynamics.

Simultaneously, we also generated Gateway compatible tools for optogenetic-(light) and chemogenetic-regulated (Doxycycline, RU486) gene expression and validated their use not only in vitro in cultured mammalian cells, but also in vivo in mice and zebrafish. We extended these chemically-inducible gene expression systems by making them Cre-dependent, expanding the utility of these Gateway compatible tools with piggyBac and AAV-compatible vectors. At the same time, we generated novel Cre-dependent tools, as well as reagents for the robust site-specific recombinase Dre, for in vitro and in vivo studies. These advances, as well as our validation of novel Tol2-dependent “all-in-one” Cre/loxP and LexA/LexO compatible vectors for use in zebrafish open new possibilities for manipulating gene expression in both time and space in the developing and adult zebrafish, mouse, and chick, as well as in numerous in vitro systems.

## MATERIALS AND METHODS

### BP Cloning

Entry vectors were created following the recommendations described in the Multisite Gateway manual (Invitrogen), with minor modifications. In most cases, att sites were added onto the ends of DNA fragments by PCR using either Q5 High-Fidelity DNA Polymerase (NEB Cat # M0491) or Platinum Taq DNA Polymerase High Fidelity (Life Technologies, Cat #11304011). For 5’ ENTRY clones (using pDONR P4-P1R), the forward PCR primer contained an *attB3* site preceding the template specific sequence, while the reverse primer contained a reverse *attB1* site following the template specific sequence. For generation of middle ENTRY clones (using pDONR221 or pENTR1a), the forward primer contained an attB1 site, while the reverse primer contained an attB2 site. For generation of 3’ ENTRY clones (using pDONR P2R-P3), the forward primer contained an attB2 site, while the reverse primer contained a reverse *attB3* site. **Table 1** lists all *att* sequences used in these PCR reactions.

Following PCR, the amplified products were either PCR purified (Qiagen PCR Cleanup kit), or gel extracted using the Qiaquick gel extraction kit (Qiagen, Cat #). As noted by others (REF), storage of purified PCR products, even at −20, significantly decreased recombination efficiency. Thus, 150 ng of PCR product was immediately combined with 150 ng of pDONR221 (pME), then mixed with 2 μL of BP Clonase II enzyme mix (Invitrogen, Cat # 11789020) and TE (pH 8.0) in a total volume of 10 μL and incubated overnight at room temperature. The following day, the reaction was terminated by the addition of 1 μL of Proteinase K (2 μg/μL) and incubated at 37°C for 10 minutes, then transformed into chemically competent STBL3 bacteria (Invitrogen) and plated on LB agar plates containing the appropriate antibiotic (in most cases kanamycin, unless otherwise noted). Recombinant clones were screened by colony PCR using primers that flanked the insert, and all positive clones were then confirmed by Sanger sequencing across the entire Entry cassette and through the flanking *att* arms.

### p5E Construct Generation

The p5E-*ICAM2*-*c-fos* clone was generated by performing a BP reaction between a gene block (Twist Bioscience) containing the 140-bp minimal human *ICAM2* promoter upstream of the 93-bp murine c-fos minimal promoter, flanked by *attB4* and *attB1* sites, and pDONR P4-P1r. A similar strategy was used to amplify the minimal 140-bp promoter without c-fos, and this was cloned into p5E via XhoI and SacII after amplification with the following oligos: oJDW 1774 (FWD <u>XhoI</u>): 5’-aaaa<u>CTCGAG</u>GTAGAACGAGCTGGTGCACGTGGC, oJDW 1775 (REV <u>SacII</u>): 5’-aaaa<u>CCGCGG</u>CCAAGGGCTGCCTGGAGGGAG.

The full-length 397-bp human *ICAM2* promoter was amplified by PCR from a geneblock, using Addgene plasmid #99736 ^54^ as a template for the design, and adding XhoI and SacII sites to the forward and reverse primers containing *attB4* and *attB1* sites, respectively, and the purified PCR product was digested and inserted into p5E-MCS (Chien lab plasmid #228)^4^ using the same sites.

The human *CLDN5* promoter and chimeric intron was generated by PCR using Addgene plasmid #29299 (pEMS1503) ^55^ and following oligos: oJDW 1948 (CLDN5 FWD + Sal I) 5’-aaaa<u>GTCGAC</u>AACCCTTAATGAATTCGAGCTCCTAGGC and oJDW (1949 CLDn5 REV + <u>HindIII</u>) 5’-aaaa<u>AAGCTT</u>TTTGAAGAATAGGAACTTCGGAATAGGAA CTTCC. The resulting amplicon was digested with SalI and HindIII, then cloned into p5E-MCS using the same sites to generate p5E *CLDN5* (JDW 1120).

To construct p5E-5xC120-c-Fos (JDW 1241), a cassette containing a SV40 polyA followed by a transcriptional pause site, then a NOS terminator, then 5X C120 sites followed by the murine c-Fos minimal basal promoter was amplified from JDW 1109 (pME-TAEL-c-Fos) using the following primers: oJDW 2185 (p5E FWD) (66-mer):5’-ggggACAACTTTGTATAGAAAAGTTGGAATTCTAGCAGACATGATAAGATACATTGATGAGTT TGG and oJDW 2186 (p5E REV) (58-mer): 5’-ccggAGCCTGCTTTTTTGTACAAACTTGATCTAGAACTAGTGCGGCCGCGTTAGTTGC, which contain 5’ attB4 and 3’ attB1 sites, and the PCR product was cloned into pDONR P4-P1r by BP clonase.

p5E *CDH5* (JDW 1239) was generated by PCR amplification of a 2,486 bp fragment and 15 bp of an exon from murine genomic DNA using the following primers containing a 5’ attB4 and 3’ attB1 site: FWD, 5’-ggggACAACTTTGT ATAGAAAAGTTGCTCGAGGTCGACt CTAGTAGCAGAAACAAGG, REV, 5’-ccggAGCCTGCTT TTTTGTACAAACTTGAGAA TTCAGGGCCGAGCTTTGTGGAGAGCAC.

p5E-human EF1a (JDW 1164) was generated by digesting p5E-MCS (JDW 459) (a kind gift of the late Dr. Chi-Bin Chien, University of Utah) with HindIII and BamHI and human EF1a was subcloned using the same sites from pAAV-oChIEF-P2A-TdTomato-WPRE-bGH-A (Addgene # 51094). p5E-EFS (the EF1a core promoter, also known as EF-1α short) was cloned by PCR amplification of only the core promoter and not the downstream intron of human EF-1α.

The human GLAST promoter was generated by PCR with the following primers that contained a 5’ attB4 and a 3’ attB1 site: FWD, 5’-ggccCAACTTTGTATAGAAAAGTTGATCGATAGGTACCATGTCTACACAAACtg, REV, 5’-AGCCTGCTTTTTTGTACAAACTTGACTAGTccggtGGATCTCGAGCCCatcaagc. The resulting amplicon was purified by PCR Cleanup (Qiagen) then recombined with pDONR P4-P1r in a BP reaction to generate p5E Glast (JDW 1182).

p5E CAG was created by inserting the CAGGS promoter into p5E-MCS via KpnI to BamHI. p5E TetO(8x) CMV minpro was created by subcloning the TRE and downstream minimal CMV promoter from pB-TA-ERN into p5E MCS via PspXI and SacII using standard T4 DNA ligation. p5E-TRE3GV (JDW 1163) was subcloned by digesting JDW 447 (pBT346.6-TRE3G-FRT) with XhoI and HindIII and shuttled into p5E-MCS (JDW 459).

p5E Ubi-pro, which contains the zebrafish *Ubiquitin* promoter, exon1, and intron 1, was made by digesting p5E-*Ubi*-loxP-EGFP-loxP (Addgene #27322)^153^ with BamHI, and ligating the digested product together to remove the loxP flanked EGFP cassette and reconstitute the BamHI site.

The ∼900 bp promoter through exon 1 of zebrafish *myl7* (also known as *cmlc2*) was designed as a gene block using pTol2DestCG2 ^4^ as a template for the PCR reaction with flanking attB1 and attB2 sites to generate a pan cardiomyocyte 5’ entry clone p5E-*myl7-pro* (JDW 1148) active in zebrafish.

The 544 bp human *Troponin T* (*TNNT2*) promoter was amplified from an existing template using primers containing a 5’ flanking attB4 and 3’ attB1 site: FWD 5’-GGGGACAACTTTGTATAGAAAAGTTGaaagcttCTCAGTCCATTAGGAGC and REV 5’-ccggAGCCTGCTTTTTTGTACAAACTTGAaccggtGAATTCCTGCCGACAGATCCTGG and then recombined with pDONR P4-P1r (p5E Donor) to make a mammalian pan-cardiomyocyte promoter clone p5E-*TNNT2* (JDW 1208).

The muscle specific *unc503* promoter construct from the endogenous zebrafish *unc-45* locus, described in a previous publication ^58^, was ordered as a gene block with flanking *attB4* and *attB1* sites, respectively, and inserted into pDONR P4-P1r by BP recombination. JDW 1320 (*p5E-Crestin*) contains an 844 bp element from a previously described ∼1 kb neural crest-specific enhancer from zebrafish *crestin* ^59^ followed by a minimal murine *c-Fos* promoter, then a 652 bp rabbit beta globin intron was synthesized as a gene block with flanking *attB4* and *attB1* sites, respectively, and inserted into pDONR P4-P1r by BP recombination. All p5E clones were confirmed by Sanger sequencing.

### pME Construct Generation

pME-MCS-WPD (JDW 455) was construct by annealing two oligos into a double stranded duplex then recombining the dsDNA into pDONR221 via BP reaction to generate a novel pME-MCS vector with a multiple cloning site consisting of EcoRI-SalI-BamHI-KpnI-SmaI-NotI-XhoI-EcoRI flanked by *attL1* and *attL2* sites.

pME-AmCyan was created by amplifying AmCyan from the vector pME lox-AmCyan-stop-lox (a kind gift of Drs. Caroline and Geoff Burns at Boston Children’s Hospital). The insert is flanked 5’ by <u>AgeI</u> before the kozak, then <u>XhoI</u> at the 3’ end after the stop codon. The AmCyan insert with flanking *attB1/B2* sites was amplified by PCR using the following oligos: oJDW 2012 (pME AmCyan FWD) 5’-gggg*ACAAGTTTGTACAAAAAAGCAGGC<u>a</u>*<u>ccggt</u>CGCCACCATGGCCCTG TCC, oJDW 2013 (pME AmCyan REV) 5’-cccc*ACCACTTTGTACAAGAAAGCTGGG<u>c</u>*<u>tcgag</u>CTTCAGAAGGGCACC. The purified PCR product was cloned into pDONR21 by BP reaction to generate pME-AmyCyan1 (JDW 1151).

pME-H2B-mCerulean (JDW 1150), was created by amplifying H2B-mCerulean from pME-*loxP*-H2B-mCerulean-stop-*loxP* ^154^ (a kind gift of Drs. Michael Harrison and Ching-Ling Lien at Children’s Hospital Los Angeles) using the following primers: oJDW XXXX (FWD) 5’-gggga*CAAGTTTGTACAAAAAAGCAGGC<u>T</u>*<u>CTAGAgaattcggtacc</u>GCCACCATG CCAGAGCC and oJDW XXX ^88^: 5’-ccccacc*ACTTTGTACAAGAAAGCTGGG*<u>ctcgaggcggccgc</u>ttaTTACTTGTACAGCTCGTCCATGCC. The purified PCR product was cloned into pDONR221 by BP reaction to generate pME-H2B-mCerulean. The attL1/L2 flanked insert contains 5’ <u>XbaI</u> and <u>KpnI</u> sites upstream of a Kozak sequence followed by human Histone 1 H2Bj, then BamHI and AgeI restriction sites preceding mammalian codon optimized mCerulean^68^, followed by two stop codons, then NotI and XhoI at the 3’ end.

pME-mRuby2-3xMYC (JDW 669) was created using the following oligos: oJDW 836 (mRuby Kozak attb1 FWD) 5’-GGGGACAAGTTTGTACAAAAAAGCAGGCTGCCACCATGGTGTCTAaGGGCGAAGAGc and oJDW 837 (mRuby attb2 REV) 5’-GGG GAC CAC TTT GTA CAAGAA AGC TGG GTCTCAtcaggatctcaggtcctcc, followed by a BP reaction and transformation into *Stbl3* bacteria. A related clone, pME-P2A-Lifeact-mRuby2-3xMYC (JDW 678) was generated for cloning 5’ to the P2A peptide via an EcoRI site.

pME-myr-mTagBFP (JDW 1183) was generated by PCR addition of a 5’ attB1 site upstream of a Kozak sequence and the eight amino acid myristoylation sequence from MARCKS (MGCCFSKT), which is sufficient to target proteins to the plasma membrane ^103,155^ using the following primers: oJDW 2068 (attB1 FWD) 5’-TATCACAAGTTTGTACAAAAAAGCAGGCT and oJDW 2069 (attB2 REV) 5’-ATATCACCACTTTGTACAAGAAAGCTGGGT.

pME-V5-mTagBFP2 contains an attL1/L2 flanked insert with BamHI, SpeI, and NcoI upstream of a Kozak, followed by a V5 epitope tag then a Gly-Gly-Ala-Gly-Gly (G_2_AG_2_) flexible linker, where the a Met-Val-Ser-Lys-Gly-Glu insertion replaces the first Met-Ser-Glu residues of mTagBFP2 (p.I174A) ^65^, followed by HindIII, PacI, a stop codon, AscI, XhoI, and BglII and was shuttled from an AAV2 CAG-V5-mTagBFP2-WPRE-SV40-pA (to be described elsewhere) into an existing pME clone via SpeI to AscI. pME-3xNLS-mTagBFP-WPRE-bGH-pA (JDW 484) was constructed by PCR sewing.

Briefly, mammalian codon optimized 3xNLS-mTagBFP-FLAG and a WPRE-bGH-polyA from Ai9 (Addgene #22799) ^156^ were amplified, and then the entire cassette was cloned into pDONR221 via a BP reaction. A similar strategy was used to generate pME-3xNLS-mKate-V5-WPRE-bGH-pA (JDW 485) and pME-3xNLS-EGFP-WPRE-bGH-pA (JDW 488).

The open reading from, including a 5’ kozak, of mRuby2 followed 3x MYC tag and stop codon was amplified from an existing clone containing P2A-mRuby-3xmyc-STOP (JDW 498, pUC57-KAN-FLEX-rtTA TetOn-deGradFP P2A mRuby2-3xMYC, to be described elsewhere) using the following primers: FWD-GGGGACAAGTTTGTACAAAAAAGCAGGCTGCCACCATGGTGTCTAaGGGCGAAGAGc and REV-GGG GAC CAC TTT GTA CAA GAA AGC TGG GTCTCAtcaggatctcaggtcctcc and recombined with pDONR221 to generate pME-mRuby2-3xMYC-stop (JDW 669). pME-EGFP-miR was based on pg4a (a kind gift of Dr. Michael McManus at UCSF).

Briefly, using the pEGFP-N3 construct ^34^ as template, an artificial MCS and intron were inserted upstream of sequence from human *HPRT* locus, with multiple cloning sites for subsequent insertion of genomic loci containing microRNAs. Primers including approximately 500-700 bp of sequence surrounding either the *miR-500* and *miR-126* locus were designed for cloning into the BamHI and PstI sites of pg4a. The micro-RNA containing GFP inserts, or the empty intron GFP alone, were then amplified with flanking attB1 and attB2 sites to create pME-EGFP-miR (JDW 1231), pME-EGFP-miR-500 (JDW 10), and pME-miR-126 (JDW 1401), respectively.

pME Vhh-sfGFP-P2A-HA-iRFP-caax (JDW 1215) was synthesized by GeneUniversal and contains an actin nanobody, whose sequence is derived from Addgene #159595 (an actin nanobody with a LOV domain insertion) ^92^. In this clone, the Vhh nanobody cDNA is followed by a flexible Arg-Ser-Leu-(Gly_4_Ser)_4_ linker and sfGFP to create a fluorescent chromobody that recognizes actin, followed by an Ala-Ser-Gly-Ser-Gly linker, a Gly-Ser-Gly-P2A (GSG-P2A) cleavage peptide, a single HA epitope and then iRFP, followed by a linker with a NotI and XhoI site for insertion of another copy of iRFP to subsequently generate tdRFP, then a GHGTGSTGSGSSGRSG amino acid linker and the CAAX domain from HRAS for cell membrane localization. In comparison to other nanobodies targeting actin, we note that this clone (like the parental version at Addgene) lacks three amino acids residues (Phe Val Lys) in the CD2 framework region (near amino acid 63). This did not appear to affect recognition of F-actin. However, this difference was corrected in subsequent actin nanobodies that were created as part of the MAGIC toolkit. We further modified this vector to insert another copy of iRFP, creating pME-Vhh-sfGFP-P2A-HA-tdiRFP-caax (JDW 1311).

pME 3xHA-BirA-T2A-mCherry-caax was generated by PCR using the following primers oJDW1306 (attB1_*kozak*_HA_BirA FWD): 5’-aaggggACAAGTTTGTACAAAAAAGCAGGCT*gcc acc*ATGGCCACCTATG, oJDW1307 (attB2r_*stop*_CAAX REV): 5’-aaggggACCACTTTGTACAAGAAAGCTGGGT*tca*GGAGAGCACACACTTGCA and Addgene #80056 ^157^ as a DNA template.

To generate pME-Luc-P2A-H2A-mCherry (JDW 926), luciferase was amplified from JDW 842 (pB-CAG-EGFP-T2A-Luciferase) (a kind gift of Dr. Benjamin Deneen at Baylor College of Medicine) along with a BamHI and SpeI site and upstream Kozak consensus protein translation initiation sequence (GCC ACC ATG G) with primers oJDW 1368 (Luc FWD) 5’-GGGGggatccactagtgccaccATGGaagacgccaaaaacataaagaaagg and oJDW 1369 (Luc REV) 5’-CCCCCGGTACCcacggcgatctttccgcccttcttgG with a 3’ in frame KpnI site (Gly-Thr-Gly). The resultant PCR amplicon and JDW 714 (pME-V5-KRAS-G12V-P2A-H2AZ/F-mCherry) were then digested with BamHI and KpnI, followed by a standard T4 DNA Ligation.

pME-Golgi-mTagBFP2-HA-P2A-H2A-iRFP (JDW 1007) was synthesized by Gene Universal. The construct has a c-terminal linker with NotI and XhoI sites for the subsequent digestion and insertion of a second copy of iRFP to generate pME-Golgi-mTagBFP2-P2A-H2A-tdRFP (JDW 1079).

pME-Dre-nls (JDW 31) was amplified from pCAGGS-Dre-nls (a gift from A. Francis Stewart, Biotechnology Center TU, Dresden, Germany) using the following oligos: FWD (Topo-Kozak-Dre F) 5’-CACCGTCGCCACCATGGGT GCTAGCGAGCTGATCATC and REV (DRE-NLS-Rev-double stop) 5’-TCATCACACTTTCCTC TTCTTCTTAGGACC. The resultant PCR amplicon was purified and directionally TOPO cloned into pENTR/D-TOPO. pME-Dre-ERT2 (JDW 1344) was made by PCR stitching Dre, minus the nls, to ERT2 ^158^, and inserting a CVRGS linker (5’-TGCGTACGCGGATCC-3’) ^159^ between Dre and ERT2.

pME-mScarlet-P2A-iCre (JDW 917) was created by PCR of an existing template within the lab (pCS2-mScarlet-P2A-iCre) with the following primers: oJDW 1342 (attB1-*kozak*-mScarlet-Fwd): aaggggACAAGTTTGTACAAAAAAGCAGGCT*gccaccATGG*TGAGCAAGG, oJDW 1343 (attB2-*stop*-iCre-Rev): 5’-aaggggACCACTTTGTACAAGAAAGCTGGGT*tta*GTCCCCATCCTCGAGCAG. The resultant amplicon was PCR purified, then recombined in a BP reaction with pDONR221.

pME-3xNLS-G_2_AG_2_-mTagBFP2-G_3_-2xHA-GR_2_RGP-G_2_AG_2_-Cre (JDW 918) was PCR amplified from pCS2-3xNLS-mTagBFP2-2x-HA-Cre (to be described elsewhere) using the following oligos: oJDW 1346 (attB1-kozak-nlsTagBFP-FWD) 5’-aaaaggggACAAGTTTGTACAA AAAAGCAGGCT*cgccaccATG*CCAAAAAAGAAG and oJDW 1347 (attB2-*stop*-Cre-Rev) 5’-aag gggACCACTTTGTACAAGAAAGCTGGGT*cta*ATCGCCA TCTTCCAGCAGG. The resultant PCR amplicon was purified, then recombined in a BP reaction with pDONR221.

mTurquoise was amplified from pmTurquoise-H2A (Addgene #36207)^70^ then cloned into pAAV-EF1a-EGFP-P2A-iCre to replace EGFP via BamHI to BsrGI digestion. The resulting pAAV-EF1a-mTurquoise-P2A-iCre vector was used as a template for PCR with the following oligos: oJDW 1344 (attB1_mTurquoise-Fwd): 5’-aaggggACAAGTTTGTACAAAAAAGCAGGCT *gccaccATGG*TGAGCAAGG and oJDW 1345 (attB2-stop-iCre-Rev) 5’-aaggggACCACTT TGTACAAGAAAGCTGGGT*tta*GTCCCCATCCTCGAGCAG. The resultant PCR amplicon was purified, then recombined in a BP reaction with pDONR221 to generate pME-mTurquoise-GSG-P2A-iCre (JDW 920).

To generate pME-EGFP-GSG-P2A-iCre (JDW 921), pCS2-EGFP-GSG-P2A-iCre (to be described elsewhere) was used as a template for PCR with the following oligos: oJDW 1340 (attB1-*kozak*-EGFP-Fwd): 5’-aaggggACAAGTTTGTACAAAAAAGCAGGCT*gccaccATGG*TGAG CAAGGGCG AG and oJDW 1341 (attB2-stop-iCre-Rev): 5’-aaggggACCACTTTGTACAAGAAA GCTGGGT*tta*GTCCCCATCCTCGAGCAGC. The resultant PCR amplicon was purified, then recombined in a BP reaction with pDONR221.

pME-*loxP*-3xSTOP-*loxP* (JDW 869) was created by removing H2B-mCerulean from pME-*loxP*-H2B-mCerulean-STOP-*loxP* (a gift of Drs. Ching-Ling Lien and Michael Harrison) by digestion with HindIII and NotI, then cloning in a gene block with a bGH and SV40 polyA sequence upstream of the already present SV40 polyA to generate pME-HindIII-BamHI-SV40pA-bgHpA-SV40pA-KpnI (JDW 869). To generate a rox-flanked red fluorescent reporter for use with Dre recombinase, a PacI-AgeI-*rox*-HindIII-KpnI-SacII-Kozak-NcoI-3xNLS-BamHI-mCherry-V5-SnaBI-NotI-ClaI-2xSV40 late polyA-HindIII-PstI-*rox*-XhoI-EcoRV-MluI insert was cloned between attL1 and attL2 in pUC57 to generate pME-*rox*-mCherry 2x-stop-*rox* (JDW 1167).

Hyperactive PBase ^135^ (a gift of Dr. Benjamin Arenkiel at Baylor College of Medicine) and a downstream WPRE (without a polyA) were subcloned from Cdh5-hyPBase-WPRE-pA (to be described later) into pME between EcoRI and SalI to generate a pME-hyPBase-WPRE (JDW 679).

To create a pME vector encoding a dominant active human β-Catenin (β-Catenin^S33Y^), the cDNA insert was amplified from pCDNA-β-Catenin^S33Y^ (Addgene #19286) ^160^ using the following primers: FWD (Attb1 KOZAK S33Y FWD) 5’-GGG GAC AAGTTT GTA CAA AAA AGC AGG CT G GAT CCG CCA CCATGG CTA CTC AAG CTG ATT TGATGG and REV (Attb2 FLAG REV) 5’-CCC CAC CACTTT GTA CAA GAA AGCTGG GGC GGC CGCTTA CTT GTCATC GTC. The resultant PCR product was recombined by BP reaction into pDONR221 to generate pME-β-Catenin^S33Y^ (JDW 419).

pME-V5-mClover3-KRAS4A-WT (JDW 812) was created by cutting pAAV-sCAG-mClover3-KRAS4A-WT with SpeI and AscI to shuttle mClover-KRAS into an existing pME clone using the same sites. A similar strategy was used to generate pME-V5-mClover3-KRAS4A-G12D (JDW 813), as well as pME-V5-mScarlet-I-KRAS4A-WT (JDW 822), pME-V5-mScarlet-I-KRAS4A-G12V (JDW 823), pME-V5-mTagBFP2 (JDW 830), pME-V5-mTagBFP2-KRAS4A-WT (JDW 831), and pME-V5-mTagBFP2-KRAS4A-G12D (JDW 832).

3xFLAG-human Ubiquitin was amplified by PCR using pCDNA3-3xFlag-Ubiqutin (a gift of Dr. Andre Catic at Baylor College of Medicine) and the primers FWD 5’-GGGGACAAGTTTGTACAAAAAAGCAGGCTACTAGTGCTAGCGCCACCACCaccatggactacaaagaccatgacgg and REV 5’-GGGGACCACTTTGTACAAGAAAGCTGGGTggatcctctagagtcgac tggtacc. The purified PCR product was used in a BP reaction with pDONR221 to generate pME-3xFlag-hUbiquitin (JDW 1098).

pME-LexA-mODC (JDW 937) contains the lexA transactivator fused to amino acids 422-461 of the PEST degradation domain of murine ornithine decarboxylase (mODC) to promote rapid protein turnover ^105^. This insert, flanked by *attL1* and *attL2* sites in a minimal backbone, was synthesized by Twist Bioscience.

pME TAEL2.0 (JDW 1109) contains an attL1/L2 flanked TAEL2.0 cDNA followed by SV40 late polyA, a pause site, then NOS terminator, followed by five, 20-bp C120 sites and a murine *c-Fos* minimal promoter cassette (JDW 1109).

pME LiCre (JDW 1119) was created by PCR amplification using pGY577 (Addgene #166663) (a gift of Gaël Yvert) ^104^ as a template and the following primers: oJDW 1950 (SalI-Kozak-LiCre-FWD): 5’-aaaaGTCGACgccaccatgggtccaaaaa agaagagaaaggtagatcc and oJDW 1951 (Li Cre Rev HindIII): 5’-atccagaggttgattggatccaagc and the resulting PCR product was used in a BP reaction with pDONR221 to generate the new Entry clone.

pME rtTA3G (JDW 1230) was created by performing PCR using an existing template (p3E-C-EFS-rtTA3G, JDW 1214) and the following primers: oJDW 2124 (attB1 rtTA3G FWD) 5’-GgggACAAGTTTGTACAAAAAAGCAGGCTATGTCTAGGCTGGACAAGAGCAAAG and oJDW 2125 (rtTA3 3’ attB2 REV) 5’-ggggCCACTTTGTACAAGAAAGCTGAATTCCTATCATTACCCGGGGAGC. The resulting PCR product was PCR purified, then BP recombined with pDONR221 and create this middle entry plasmid.

### p3E Construct Generation

The WPRE and bovine growth hormone poly(A) signal cassette was amplified from pAAV-EF1a-FLEX-lifeact-mScarlet-HA-WPRE-bGHpA using the following primers: oJDW 2139 (Fwd WPREBgH) 5’-ggggACCCAGCTTTCTTGTACAAAGTGGActatcgataatcaacctctggattacaa and oJDW 2140 (rev bghwpre) 5’-ggggCAACTTTGTATAATAAAGTTGGGTACCgatgcaatttcctcattttattagg. The PCR product was then recombined into pDONR P2R P3 to generate p3E-WPRE-bGH-pA (JDW 1221).

The rat EF1-poly(A) was amplified using the following primers: oJDW 2137 (EF1a fwd) 5’-ggggACCCAGCTTTCTTGTACAAAGTGGAACGCGTATTATCCCTAATACCTGCCACC and oJDW 2138 (3’ EF1Pa rev) 5’-ggggCAACTTTGTATAATAAAGTTGGGTACCAGCTTTCTATGCAACCCAAG and pWhere-H2B-EGFP-Dest as a template, then the product BP recombined with pDONR-P2R-P3 to generate p3E-EF1a-pA (JDW 1222).

p3E-H2A-mCherry_SV40pA (JDW 967) was created by amplifying H2A-mCherry from JDW 714 (pME_V5_KRAS_G12V_p2a_H2AZ/F_mCherry) with the following primers: oJDW 1529 (5’-aaaaGGATCCTAGGATGGCAGGTGGAAAAGCAGG) oJDW 1018 (CTTTGTACAAGAAAGCTGGGAGATCTCTCGAGCTATCATTACTTGTACAGCTCGTCCATGC) to add BamHI and XhoI restriction enzyme sites. The PCR product and vector JDW 817 (p3E-V5-mScarletI-KRAS4A-G12V-SV40pA) were digested with BamHI and XhoI to remove the insert in the backbone, and then ligated together and then transformed into Stbl3 competent cells.

A 3’ entry clone for three-way multisite gateway LR reactions, containing a V5-mScarlet-I fluorescent reporter with a stop codon followed by a poly(A) signal, p3E-V5-mScarlet-I-SV40pA (JDW 968), was created by amplifying the V5-mScarlet insert from JDW 807 (attB-AC-zsY-V5-mScarlet-I-KRAS4AG12V) with oJDW 1037 (5’-ATATCTCTCGAGGGCGCGCCTTACTTGTACAGCTCGTCCATGCC) and oJDW 1230 (5’-agaattGGATCCACTAGTGCTAGCgccaccatgGGTAAGCCTATCCC) to add BamHI and XhoI cut sites. The insert and JDW 817 (p3E-V5-mScarletI-KRAS4A-G12V-SV40pA) were digested with BamHI and XhoI to remove the insert in the backbone, and then ligated together and then transformed into Stbl3 competent cells. 434-924-0000 (walk in lab tomorrow) p3E-V5-mClover3 (JDW 1318) was constructed by performing a BP reaction with a gene block sequence containing attB2-V5-mClover3-SV40-pA-attB3 and pDONR P2R P3.

To generate p3E-V5-mTagBFP2-pA (JDW 871), the existing plasmid JDW 824 (pAAV-sCAG-V5-mTagBFP2-WPRE-SV40pA) was cut with SpeI and AscI and the resulting V5-mTagBFP2-SV40-poly(A) fragment was then inserted into JDW 814 (p3E-V5-mClover::KRAS4A(WT)-SV40pA) that was cut and gutted with the same enzymes. V5 is separated from the mTagBFP2 insert by a flexible Gly-Gly-Ala-Gly-Gly (G_2_AG_2_) linker.

p3E-EFS-rtTA/rtTA3 (JDW 1214) was created by ordering a gene block (Twist) with the human EFS/EF1a-core promoter upstream of an rtTA-3G (Tet-On, 3^rd^ generation) transcriptional transactivator followed by an SV40 poly(A) signal, followed by recombination with pDONR P2R-P3.

### Destination Vector Generation

The backbone for pCAGEN is pCAGGS (Niwa et al. Gene 108, 193-199 (1991)), from Dr. J. Miyazaki (Osaka University), and the multiple cloning site (MCS) of pCAGGS was modified by adding an EcoRI, XhoI, EcoRV and NotI site to generate pCAGEN. pCAGEN, a kind gift from Connie Cepko (Addgene Plasmid #11160)^124^, was further modified by inserting a Gateway ccdB RFA cassette into the EcoRV site of MCS to generate pCAGEN-DEST. pCAGGS-DEST-WPRE variant was generated by insertion of WPRE bGH polyA fragment by Cold Fusion (SBI Systems) cloning between the XhoI and BglII sites. For the pDEST-CAGEN-IRES vectors, pCAGEN-Dest-IRES-myr-BFP (JDW 491), pCAGEN-Dest-IRES-myr-mKate2 (JDW 494), and pCAGEN-Dest-IRES-myr-EGFP (JDW 495), PCR amplicons of each respective fluorophore, including an additional 18 amino terminal amino acids from GAP43 (MLCCMRRTKQVEKNDEDQKI) ^161,162^ that anchors proteins to the inner plasma membrane, were inserted by restriction digest and ligation into pCAGEN-Dest. Constructs were transformed into chemically competent *ccd*B-bacteria (strain, Invitrogen) and then plated on LB agar plates with ampicillin and chloramphenicol. Transformants were screened by colony PCR for the presence of the insert, then confirmed by Sanger sequencing.

The Mef2c-F6/frag3 plasmid ^163^, a kind gift of Brian Black (UCSF), was digested with XhoI and SalI to isolate the 3,970 bp SHF/AHF enhancer fragment, while pCAGEN was cut with SpeI and XbaI to remove the CAGGS promoter. The resulting fragments were blunted and ligated together to make the Destination vector pMef2c-AHF-Dest (JDW 476).

The β-MHC-promoter (murine *MyH7*) clone #32 in a pBSKII backbone, a gift from Dr. Jeffrey Robbins’ lab (Addgene #53963)^127^, was modified by digestion with HpaI and insertion of an Destination Reading Frame “A” (RFA) cassette to create the embryonic, pan-cardiomyocyte transgenic expression vector pβ-MHC-promoter Dest (JDW 26).

The African clawed frog *Xenopus laevis Myosin Light Chain 2* promoter ^164^, *XMLC2-*pro, is sufficient to drive expression of Cre recombinase in cardiomyocytes from early embryogenesis through adulthood in mice, with labelling evident in all four chambers and the cardiac outflow tract ^165^. This 3.0 kb region was amplified by PCR using 5’ and 3’ oligos that both contained a HindIII site, and pEF-DEST51 (Invitrogen) was digested with the same enzyme to remove the EF1a promoter in exchange for *XMLC2-pro* to generate pxtMLC2-Dest (JDW 7).

pGlast-DEST (JDW 1009) was generated by removing PBase from Glast-PBase (a kind gift of Dr. Benjamin Deneen at Baylor College of Medicine) via PspXI and NcoI, then inserting an *attR1*-Cm^R^-*ccd*B-*attR2* cassette downstream of the human *GLAST* promoter via Cold Fusion cloning. The Gateway cassette was amplified with the following primers: FWD 5’-cctcgttactgcttgatGGGCTCGAGATCc tgttttgacctccatagaagacaccg and REV 5’-CGGCCGGCCGCCCCGACTCTAGATGCATGCattcgatgggggatcccttcg using JDW 931 (pB_TetOn_DEST_EFS_mODC_rtTA_IRES_NEO) as a template.

pCAGGS-*rox*-nKateV5-*rox*-Dest-FRTNeo (JDW 472) was generated by cloning 3x-NLS-mKate2-V5 followed by a rabbit globin polyA sequence between *rox* recombination sites downstream of a CAGGS promoter. Then, a Gateway RFA Destination cassette and a rabbit globin polyA sequence were cloned downstream to generate pCAGGS-rox-nKateV5-rox-Dest. Subsequently, a SacI-SalI fragment containing an FRT-PGK-FRT cassette from pK11 (Courtesy of Dr. Gail Martin at UCSF) was inserted 3 to the Dest cassette in the opposite orientation to generate pCAGGS-rox-nKateV5-rox-FRT-murine PGK promoter-NEO-FRT. This substrate was used in an LR reaction with pME-myr-mTagBFP2-Flag (JDW 1183) to generate pCAGGS-rox-nKateV5-rox-myrBFPflag-frt-PGK-NEO-frt (JDW 1217).

pB-TA-ERN (Addgene # 80474), a gift from Knut Woltjen, with AvrII and a fragment containing a pause site followed by the human EF1a minimal promoter (also known as EFS) and a destabilized rtTA-3G (synthesized by Twist Biosciences) was inserted to create a *piggyBac* transposon flanked Tet-On Destination vector compatible with pME clones: pB-Tet-On-DEST-EFS-mODC-rtTA-IRES-NEO (JDW 931). Conversely, a *piggyBac* ITR flanked Tet-Off vector was created by amplifying an *attR2*-ccdB-Cm^R^-*attR2* with the following primers oJDW 1398 (FWD) 5’-AAAAGCTAGCGATTCGAATTCAAGGATCAACAAGTTTGTAC and oJDW 1399 ^88^ 5-AAAAGGCGCGCCGCTCGAGAGGATCAACCACTTTG, using pCS2-DEST as a template for the PCR reaction. The Destination cassette was inserted downstream of a TRE-tight promoter via NheI and AscI restriction enzyme sites between two inverted pairs of flanking *lox2272* and *loxP* sites, followed by a WPRE and bGH polyA sequence, another polyA signal, then an RNA Pol II transcriptional pause site from the human *α2 globin* gene, followed by a human EFS promoter and a destabilized tTA (mODC-tTA, also known as d2tTA) followed by an SV40 polyA signal, all flanked by two *piggyBac* ITRs to generate pB-Tet-Off-FLEX-Dest-EFS-mODC-tTA (JDW 936).

pDestTol2-2xIns; *hsp70*-zCreI-BFP-mODC (JDW 1002) was digested with SalI and Cold Fusion (SBI Bioscience) was done to insert a Gateway recombination cassette upstream of the polyA using the primers and JDW 760 (pTol2Dest-pA2) as a template with the following primers: oJDW 1630 (FWD): 5’-AACACAGGCCAGATCCTAGGGGGCCCGTTTAAACGCCATG ATTACGCCAA GCTATCAACTTTGT and oJDW 1631 ^88^: 5’-ATTTGTAACCATTATAAGCT GCAATAAACAAGTTGATCATCATCGATGGTACCGTAAAACGAC. pDestTol2-2xIns; *hsp70*-zCreI-BFP-mODC (JDW 1002) was digested with MluI and AsiSI to remove the entire zCre-intron-mTagBFP2-mODC cassette, and a shorter cassette without the mODC tag was amplified using the primers oJDW 2064 (BFP-G2AG2-NLS-AsiS1) 5’-AATTACGCGTGCCGCA

TATGGCCACCatggtgcccaagaagaagagg and oJDW 2065 (zCreI-Mlu1) 5’-AATTACGCGTGCC GCATATGGCCACCatggtgcccaagaagaagagg, and inserted using those same restriction sites to generate pDestTol2-2xIns; *hsp70*-zCreI-BFP (JDW 1184). This same pDestTol2 plasmid, JDW 1184, was then digested with MluI and SpeI to remove the *hsp70* promoter, while the endothelial-enhancer/promoter from zebrafish fli1ep was then amplified from an existing plasmid (JDW 897, pTol2-fli1a-mScarlet-mClover3-Dll4) using the following primers: oJDW 2090 (Spe1 5’ homology JDW 897) 5’-ggccACTAGTATCTCATCTTGACCCATAAACATACACTAAAACC and oJDW 2091 (Mlu1 3’ homology JDW 896) 5’-aaggccACGCGTCGGATGGTTTTTTTTCCTCTAAATTTGGGAA and inserted using the same MluI and SpeI sites to generate an Gateway multisite plasmid with fli1-driving Cre and BFP in the backbone (pDestTol2-2xIns; *fli1ep*-zCreI-BFP, JDW 1218).

### LR Cloning

Gateway two vector, three vector, and four vector recombination was done by mixing equal amounts (150 ng) of each Entry plasmid and a single Destination vector with 2 μL of LR Clonase II Plus enzyme mix (Invitrogen, Cat # 12538120) and TE in a total reaction volume of 10 μL and incubated overnight at room temperature. The following day, the reaction was terminated by the addition of 1 μL of Proteinase K (2 μg/μL) and incubated at 37°C for 10 minutes, followed by transformation into chemically competent STBL3 bacteria (Invitrogen, Cat #C737303) and plating on LB agar plates containing the appropriate antibiotic (in most cases ampicillin, unless otherwise noted). Recombinant clones were screened by colony PCR using primers that flanked the insert and all positive clones were then confirmed by Sanger sequencing across each *att* recombination site.

An LR reaction between pME-Luciferase-P2A-H2A-mCherry (JDW 926) and pB-Tet-On-DEST-EFS-mODC-rtTA-IRES-NEO (JDW 931) was used to generate pB-Tet-On-Luciferase-P2A-H2A-mCherry-IRES-NEO (JDW 945). An LR reaction between pME-Luciferase-P2A-H2A-mCherry (JDW 926) and pB-Tet-Off-FLEX-Luciferase-P2A-H2A-mCherry (JDW 970) to generate pB-Tet-Off-FLEX-Luciferase-P2A-H2A-mCherry (JDW 970). An LR reaction between pME-Luciferase-P2A-H2A-mCherry (JDW 926) and pCAGEN-Dest (JDW 471) was used to generate pCAGEN-Luciferase-P2A-H2A-mCherry (JDW 1129). pB-Tet-On-DEST-hEF1a-mODC-rtTA-IRES-NEO (JDW 1086) was used in an LR reaction with pME-Luciferase-P2A-H2A-mCherry (JDW 926) to generate pB-Tet-On-Luciferase-P2A-H2A-mCherry-hEF1a-mODC-rtTA-IRES-NEO (JDW 1130), while JDW 926 was recombined with pB-TA-ERN (Addgene # 80474) to generate pB-Tet-On-Luciferase-P2A-H2A-mCherry-CMVie-rEF1a-rtTA-ires-NEO (JDW 1154).

pCAGEN-Dre-nls (JDW 1131) and pCAGEN-DreERT2 (JDW 486) were generated by an LR reaction between pME-Dre-nls (JDW 31) or pME-DreERT2 (JDW 1344) and pCAGEN-Dest (JDW 471). pCAGGS-rox-nKateV5-DEST-FRT-NEO (JDW 472) was recombined with pME-myr-BFP-Flag (JDW 1183) in an LR reaction to generate pCAGGS-nK-mB (aka pCAGGS-*rox*-nKateV5-*rox*-myrBFPFlag-FRT-PGK-NEO-FRT (JDW 473). The entire cassette can be excised via AscI and PacI digestion for subsequent cloning to a targeting vector, as done for the *Hipp11* locus (*pHipp11*-*rox*-nK-*rox*-mB-FRT-NEO, JDW 479) and the *Rosa26* locus ^103^.

### Luciferase Analysis

HeLa cells (3×10^4^ cells/well) were seeded in a 96-well white polystyrene plate (Thermo Scientific, #15492) coated with fibronectin from bovine plasma (1:400 dilution of a 1 mg/mL stock) (Sigma-Aldrich, #F1141) the day before transfection. The following day, the cells were co-transfected with 100 ng of a firefly reporter vector and 20 ng of a normalizing renilla reporter (pRL-TK) using PolyJet transfection reagent (SignaGen Laboratories, SL100688) according to the manufacturer’s instructions. For tet-inducible firefly reporters, cells were treated with Doxycycline (1 ug/mL), or vehicle control, for 24 hours prior to measuring luciferase activity using the Dual-Glo Luciferase Assay System (Promega, #E2920) on a BioTek Cytation-1 Cell Imaging MultiMode Reader (Agilent) 48 hours after transfection. Each assay was performed with technical replicates, over three experimental days (biological triplicates). Reporter activity was calculated (using the average of each technical replicate) to determine the ratio of firefly to renilla activity across all three experiments. The data were then plotted using the GraphPad software (PRISM).

### Mammalian Cell Culture and Confocal Imaging

For imaging, the day before transfection, HeLa (1×10^4^ cells/well) (ATCC, CCL-2) or HEK-293T (ATCC CRL-3216) cells were seeded in 4-well Nunc Lab-Tek II chamber slide (Thermo Scientific, 154453) coated with Poly-L-Lysine (Sigma, P6282) (0.1 mg/mL). The following day, the cells were transfected with various fluorescent reporter plasmids (0.36 μg DNA/well) using the PolyJet transfection reagent PolyJet transfection reagent (SignaGen Laboratories, SL100688) according to the manufacturer’s instructions. Forty-eight hours later, the cells were fixed with 4% paraformaldehyde/ 1X PBS at room temperature for 10 minutes, then permeabilized with 0.2% Triton X-100/1X PBS for 10 minutes and then blocked in 3% Bovine serum albumin / 1X PBS for 1 hour at room temperature. Then, the cells were co-stained with DAPI (ThermoFisher, D1306) (a 5 mg/mL stock was diluted to a working concentration of 300 nM), and in some cases Alexa Fluor 647 nm Phalloidin (ThermoFisher, A22287) (1:200) or Rhodamine (TRITC, 565 nm) Phalloidin (ThermoFisher, R415) (1:400 dilution) in 3% BSA/1X PBS for 1 hour at room temperature, and then mounted with Prolong Diamond Mounting Media (ThermoFisher, P36965). For nuclear staining of live cells, cells were incubated with Hoechst (1 μg/mL) (Invitrogen) for 30 min in a 37°C, 5% CO_2_ incubator. For HaloTag labelling, cells were incubated with 200 nM of JFX646-HaloTag Ligand (a generous gift of Janelia Fluor Dyes) for 15 min in 37°C incubator prior to fixation with 4% paraformaldehyde / 1x PBS. Images were captured using a Leica SP8 confocal microscope with the laser power set between 1-5%, using a 63x oil lens (NA=1.4). Captured z-stack images (1024 x 1024 pixels) were processed using LAS software and then exported to Image J ^166^.

### Western Blotting

HeLa cells (1×10^6^ cells/well) were seeded in a 6-well plate the day before transfection. The following day, cells were transfected with 1 ug of plasmid DNA using PolyJet transfection reagent (SignaGen Laboratories). Twenty-four hours after transfection, cells were treated with 1 μg/mL of Doxycycline (or vehicle control) for 24 hours (unless otherwise noted in the figure legend text). Then, cell lysates were extracted 48 hours post transfection in 300 mL of radioimmunoprecipitation assay (RIPA) buffer (25 mM Tris-HCl pH 7.6, 150 mM NaCl, 1% NP-40, 1% sodium deoxycholate, 0.1% SDS) plus Halt™ Protease and Phosphatase Inhibitor Cocktail (ThermoFisher, 78430), and diluted to 1x in 4x Laemmli buffer (Bio-Rad, 1610747) with 2-mercaptoethanol. Samples were then boiled for 5-10 minutes at 100°C, 20 μL of each sample was loaded onto a 12% SDS-PAGE gel for electrophoresis and transferred to Immobilon-FL PVDF membrane (Millipore, IPFL00010) using a Power-Blotter Semi-dry transfer system (ThermoFisher Scientific). Membranes were then incubated in blocking buffer (50% Intercept PBS blocking buffer (LI-COR, #927-70001) diluted in PBST) for 1 h at room temperature, then incubated in blocking buffer with rabbit anti-mCherry (Invitrogen, PAS-34974) (1:1,000) or anti-GFP (Rockland, Cat #600-401-215) (1:1,000), and mouse monoclonal anti-β-Tubulin (Invitrogen, Cat #32-2600) (1:3,000) overnight at 4°C. The next day, the blots were washed in PBST and then incubated with secondary antibodies diluted in blocking buffer for 1 hour at room temperature. The secondary antibodies used were goat anti-rabbit IgG (H+L) Secondary Antibody DyLight™ 800 4X PEG (Invitrogen, SA5-35571) (1:10,000) and goat anti-mouse IgG (H+L) Cross-Adsorbed Secondary Antibody DyLight™ 680 (Invitrogen, Cat # 35519) (1:10,000). Antibody binding was detected using the Odyssey Imaging System (LI-COR Biosciences).

### Zebrafish Experiments

Zebrafish protocols were approved by the Animal Care Committee at Baylor College of Medicine and the University of Virginia. Embryos collected from timed matings and raised in 1X E3 (5 mM NaCl, 0.17 mM KCl, 0.33 mM CaCl_2_, 0.33 mM MgSO_4_) at 28.5°C and staged according to time postfertilization and morphology ^167^. Beginning at approximately 8 hpf, embryos were incubated in 1X E3 with 0.003% (w:v) (200 μm) 1-phenyl 2-thiourea (PTU) to prevent pigment formation. The following transgenic lines were utilized: *Tg(kdrl:GFP)*^s843^ (Jin et al., 2005) and *Tg(kdrl:mCherry)^ci5^* (Proulx et al., 2010). The following lines were created: Tg(*Ubi*-lsl-EGFP-nls-mCherry; *hsp70*-zCreI-BFP)*^va^*^100^.

### Tol2-mediated Transgenesis

Tol2-mediated transgenesis (Kawakami et al., 2004) was performed as previously published^56^. Briefly, NotI linearized, gel purified, pCS2FA-transposase^4^ plasmid was used as a template for in vitro transcription using the mMessage mMachine SP6 kit (Ambion, AM1340) to generate capped, Transposase mRNA. The amplified mRNA was purified and subsequently ethanol precipitated, resuspended in TE diluted 1:10 with ddH_2_O, then quantified. A mixture of 100 ng of purified DNA (phenol:chloroform purified and resuspended in resuspended in TE diluted 1:10 with ddH_2_O), 125 ng of Tol2 transposase mRNA, 1 µL 0.8% Phenol Red/0.1 M KCl, pH 7.0, and ddH_2_O in 10 µL total volume was loaded into a microcapillary and 1 nL injected directly into the cell of one-cell stage zebrafish embryos. Embryos were then maintained at 28.5°C and processed at the indicated age for imaging.

### RU486/LexA-LexO Induction and Imaging

pTol2-based LexA/LexO nls-EGFP reporter plasmids were co-injected along with transposase mRNA into 1-cell stage AB zebrafish embryos as described above. At 24 hpf embryos with cardiac-specific expression of the *cmlc2::GFP* reporter cassette were collected and evenly distributed into 2 wells of a 6-well plates for each tested construct. A 10 mM stock of mifepristone / RU486 (Biotechne/Tocris, cat #1479) in pure DMSO was thawed from −20°C and then diluted 1:5,000 (or an equivalent amount of DMSO for the vehicle control treatment) to make a 2 μM working solution in 1X E3 with PTU (to inhibit pigmentation). Embryos treated with RU486, or vehicle control, were then incubated in the dark at 28.5°C. At ∼72 hpf, embryos were rinsed in ice-cold 1x PBS, then fixed in 4% PFA / 1x PBS in the dark overnight at 16°C with gentle agitation. The following day, embryos were washed three times in 1x PBS at room temperature, then permeabilized in three, 20 minute washes of 0.5% Triton-X / 1x PBS at room temperature on an orbital shaker, before being incubated in 300 nM DAPI (ThermoFisher, D1306) / 1x PBS at room temperature for 40 minutes in the dark with gentle agitation. Embryos were then washed in 3 times in 1x PBS, 5 minutes per wash, briefly post fixed in 4% PFA / 1x PBS, rinsed in 1x PBS, and then mounted in 1% low melt agarose in 1x PBS on a 35 mm glass-bottom tissue culture dish (Mattek, P35G-1.5-14-C) and covered in 1x PBS prior to being imaged on a Leica SP8 confocal microscope using an HC PL APO 10x/0.4 objective. *pTol2-Ubi-LexA-LexO-CAMV35S-nls-EGP-pA* injected embryos were imaged with the 405 nm channel (DAPI) at 8.54% power and a digital gain of 800. The 488 nm channel (GFP) was imaged at 10.8% power and a digital gain of 800. *pTol2-Ubi-dLexA-LexO-CMV35S-nls-EGFP-pA* and *pTol2-Ubi-dLexA-LexO-c-fos-nls-EGFP-pA* injected embryos were imaged with the 405 nm channel at 2.95% power and a digital gain of 692. The 488 nm channel was imaged at 2.4% power and a digital gain of 800. All images were captured with a pinhole size of 1 AU (53 μm). All Z-stacks were collected using a 5 μm step size and exported image sequences were processed using ImageJ and figures made using Adobe Photoshop and Illustrator.

### Heat Shock

To test the pDestTol2-*Ubi*-lsl-EGFP-nls-mCherry; *hsp70*-zCreI-BFP construct (**Figs. 8, 9**), Tg(*Ubi*-lsl-EGFP-nls-mCherry; *hsp70*-zCreI-BFP)*^va^*^100^ freshly embryos collected and incubated overnight at 28.5°C in 1X E3 / 0.003% PTU. The following day, at 24 hpf, embryos were transferred to 1X E3 / 0.003% PTU pre-warmed to 37°C and maintained at that temperature for 30 minutes. Subsequently, embryos were then rinsed once in 28.5°C 1X E3 with PTU, and then incubated at that temperature until the following day (48 hpf) when another 30-minute heatshock at 37°C was performed. Embryos were then returned to 28.5°C and then fixed at 72 hpf overnight in 4% PFA / 1X PBS at 4°C with gentle agitation and protected from light. Fixed embryos were embedded in 1.5-2.0% low-melt agarose dissolved in either 1x PBS or 1x E3 and embedded sagittally on Mat-Tek coverglass.

Embryos were imaged on a Zeiss LSM 880 using a Plan-Apochromat 20X/0.8 M27 objective. The red channel (561 nm) was imaged at 60% laser power, a digital gain setting of 632, and with a pinhole of 57 μm for both heat shock and non-heat shock conditions. The green channel (488 nm) was imaged using 12% laser power, with a gain of 607 and a pinhole of 57 μm. Widefield images were captured on a Zeiss V16.Zoom macroscope.

### TAEL Photoinduction and Imaging

Post-injection, embryos were raised in the dark, with blue light blocked by covering the incubator door, and transmitted light on the microscope for sorting of embryos or aging, with red gel filters (Pangda-Gel Filter Colored Overlays, purchased through Amazon). For photoinduction of TAEL activity, embryos were anesthetized and immobilized in 1X Tricaine / 1X E3 / 0.003% PTU / 0.5% low-melt agarose and then illuminated on a Nikon ECLIPSE Ti2 equipped with a Yokogawa W1 spinning disk unit with a Plan apo lambda 20x lens, using 488 laser at 50% power, and 1 second exposures at 1 minute intervals for approximately 10 minutes with the aperture reduced to target illumination to the region of interest. Images were acquired using a PFS4 camera.

### Murine Experiments

All mouse experiments were approved by the Institutional Animal Care and Use Committee at Baylor College of Medicine and the University of Virginia. Timed pregnancies were set up at the end of the workday. A vaginal plug in the morning was considered embryonic day 0.5 (E0.5). Appropriately timed-mated pregnant dams were used as described below.

### In utero Electroporation

In utero electroporations (IUEs) were performed as previously described ^168^. Briefly, the uterine horns were surgically exposed in a pregnant dam (CD-1 IGS strain) at E15.5 and the embryos were injected with a DNA cocktail containing the following plasmids: [1] pB-*Glast*-*loxP*-3xSTOP-*loxP*-p3E-mCherry-T2A-HRAS^G12V^, [2] a *piggyBac* (pB) helper plasmid with the glial- and astrocyte-specific promoter, *Glast*, driving expression of PB transposase (pGlast-PBase), and [3] a self-inactivating Cre recombinase, pCAGEN-self-inactivating iCre-I (JDW 1226).

The PBase helper plasmid promotes stable integration of the cargo fluorescent reporter vector, which indelibly labels all descendant cells, allowing one to visualize tumors over time via fluorescence. Following injection of the glioma-inducing cocktail (2.0 µg/µL pGLAST-PBase, 1.0 µg/µL all other plasmids) into the lateral ventricle of each embryo, embryos were electroporated six times at 100-ms intervals using BTW Tweezertrodes connected to a pulse generator (BTX 8300) set at 33 V and 55 ms per pulse. Voltage was applied across the entire brain to allow uptake of the constructs. The uterine horns were placed back in the cavity, and these dams developed normally, but their electroporated offspring featured malignancies postnatally, as the tumor suppressor deficient cells expanded. IUE tumors were harvested at postnatal day 21 (P21) from both male and female mice and processed as described below for immunohistochemistry.

### Immunohistochemistry

Brains from both male and female CD-1 mice were harvested and fixed in 4% PFA / 1x PBS overnight at 4°C in the dark with gentle agitation on an orbital shaker. The following day, brains were washed in 1x PBS and mounted in 1% agarose. 35 µm thick coronal sections were collected using a compresstome (World Precisionary Instruments, VF-300-Z) and subsequently mounted on Superfrost Plus microscope slides (Fisher Scientific, #22-037-246). Sections were dried at 50°C for 2 hours, washed once with 1x PBS, and then dried at 60°C for 1 additional hour. A barrier was then drawn around each section using an ImmEdge Hydrophobic Barrier Pen (Advanced Cell Diagnostics, Cat No #310018). Samples were rehydrated with ddH_2_O for 5 minutes before incubating them for 1 hour in blocking buffer (1x PBS / 10% Donkey Serum / 0.5% Triton-X). Mouse monoclonal anti-GFAP 488 (clone GA5) (Thermo Fisher, Cat. No. 53-9892-82) (1:50) and rabbit polyclonal anti-mCherry (Invitrogen, Cat. No. PA5-34974) (1:100) were then diluted in blocking buffer and sections were incubated with primary antibody in blocking buffer solution overnight at room temperature. The following morning, samples were washed 3x in blocking buffer. Sections were then incubated for 2 hours at room temperature in goat anti-rabbit Alexa Fluor 568 (Invitrogen, Cat. No. A-11011) diluted in blocking buffer (1:200). Sections were then washed 5x in blocking buffer, and then mounted with ProLong Gold Antifade Mountant with DAPI (Invitrogen, Cat. No. P36931).

Wholemount images post-dissection were taken with an AxioZoom V16 using a 2.5x objective with 20% overlap for tiled reconstruction of the brain. Tiled images were stitched together using Zen (Blue edition) software. Sections were imaged using a Leica SP8 confocal microscope with laser power set between 1-5% (GFP, mCherry) and ∼20% (DAPI), using a 63x lens (NA = 1.4). Z-stack images (1024 x 1024 pixels) were processed using LAS software and then exported to ImageJ.

### LiCre Induction

Mice were anesthetized by injection of a ketamine/dormitore mixture and were maintained under anesthesia using vaporized isoflurane with O_2_. Mice were then affixed to a stereotaxis apparatus synced to Angle Two software for coordinate guidance. *Ai14* Cre reporter mice ^156^ (12 weeks old) were bilaterally injected into the anterior mouse motor cortex (M1) and a unilateral injection into the posterior motor cortex (M1, AP =+0.7, DV =−1.8, ML = ±2; posterior M1, from bregma: AP =-1.0, DV =−1.3, ML = ±1.1) with 690 nL per hemisphere of a pLenti-CMV-LiCre.

Concurrently, mice were bilaterally implanted with 230 μm silica fibre optic implants (200 µm core with NA = 0.22, RWD *R*-FOC-L200C-22NA) and situated 0.1 mm above the viral injection site. Fibre optic implants were held in place by a cap made from adhesive cement (C&B Metabond Quick! Cement System (Parkell)) for initial base, and crosslinked flash acrylic (Yates-Motloid, 44115 and 44119) for headcap. Mice were allowed to recover, and viral transduction to occur, for two weeks before photoactivation. For stimulation, each animal was tethered to a dual fibre optic cord (Doric Lenses) attached to a 473nm laser source (CrystaLaser CL-2005) and then mice were chronically stimulated with trains of blue light (30 mW, 10 ms pulses, 5 or 20 Hz, 5 s trains, 30 s intervals, for 1 hour total). As a control group, the contralateral side, which was also injected and implanted, was not stimulated. Mice were euthanized one week later, and brains collected for free-floating cryosectioning and counter-staining with DAPI to label all nuclei. In some cases, mice we perfused with fluorescent lectin prior to euthanasia to label the cerebrovascular endothelium. Sections were then mounted and imaged using a confocal microscope (Leica SP8).

24 hours before transfection, HEK-293 cells were seeded into 12 well plates (Fisher, 07-200-82) at a density of 3 x 10^5^ cells/well. The following day, cells were co-transfected with 500 ng of a Cre mediated switch reporter (pCMV-lox-zsGreen-stop-lox-mCherry, JDW 514) and either a self-inactivating cre (pCAGEN-si-CreI, JDW 1226) or a light activated cre (pCAGEN-LiCre, JDW 1220). Transfection complexes were made using PEI (1:3 ratio of DNA:PEI) (Polysciences Inc, 23966). PEI:DNA complexes were allowed to form for 15 minutes and then 50 μL of OptiMEM (Thermo, 31985062) was gently added to the mixture and then the PEI/DNA/Opti-MEM mixture was added, dropwise, to the cells. Cells were then incubated in the transfection mixture for 6 hours the media was aspirated and replaced with DMEM / 10% serum / 1x non-essential amino acids / 1x pen/strep. At 24 hours post transfection, some wells were illuminated with blue light (460 nm, 1.1 watts) for a duration of 30 minutes on / 30 minutes off, for a total of 24 hours. Another group of cells was maintained in the dark. At 48 hours post transfection, both groups were fixed in 2% PFA / 1x PBS for 15 minutes and mounted with ProLong Gold with DAPI (ThermoFisher, P36935). Images were captured using a Leica SP8 confocal with laser power set between 5-10%, using a 20x lens (NA = 0.4). Z-stack images (512 x 512 pixels) were processed using LAS software and then exported to ImageJ.

## ACKNOWLEDGEMENTS

The authors thank Dr. Margot LK Williams (Baylor College of Medicine) for her assistance with the TAEL experiments, and Dr. Benjamin Deneen (Baylor College of Medicine) for the gift of piggyBac ITR flanked cargo plasmid templates and the pGlast promoter. This work was supported by grants from the National Institutes of Health (5T32GM088129-10 to W.D.T., T32HL007284 to G.L., 1R01HL159159 to W.P.D. and J.D.W), Cancer Prevention Research Institute of Texas (RP200402 to J.D.W.), and funding from the University of Virginia School of Medicine (J.D.W.).

## AUTHORSHIP STATEMENT

J.D.W. and W.P.D. conceptualized the study; J.D.W., W.P.D., W.G., Y.Z., O.E.R., J.C.III., J.O-G., and G.L., designed experiments; W.G., Y.Z., O.E.R., J.C.III, J.O-G., G.L., M.S., L.E.M., W.D.T., E.F., C.B.C., K.M.K., W.P.D., and J.D.W performed experiments; W.G., Y.Z., J.O-G., O.E.R., G.L., W.P.D., and J.D.W. analyzed data; W.G., J.O-G., and J.D.W. made the figures; G.L., W.D.T., B.R.A., W.P.D., and J.D.W. secured funding. B.R.A. and J.D.W. oversaw the studies. J.D.W. wrote the manuscript; all authors edited and approved the manuscript.

---

**Supplemental Figure 1.**
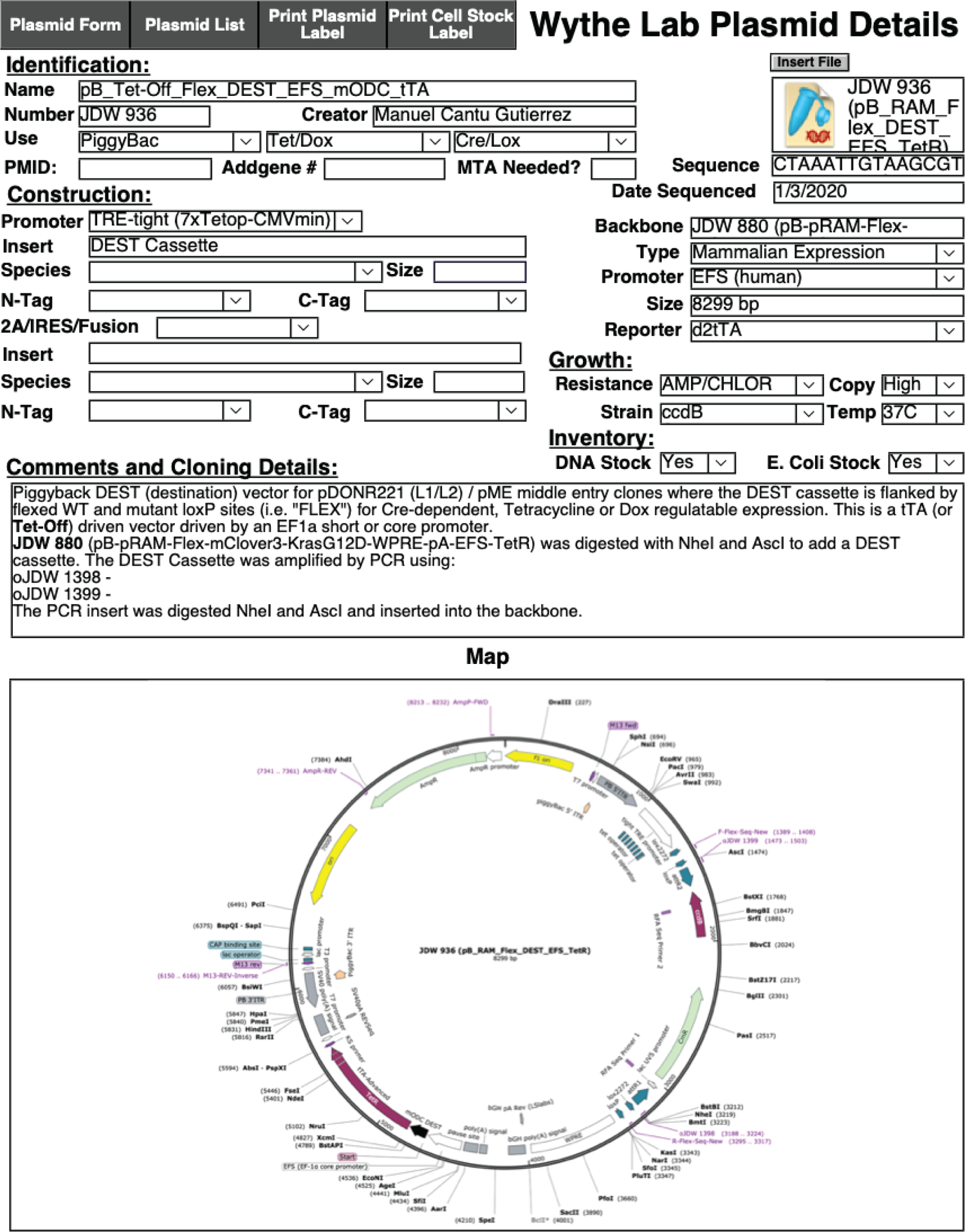

**Supplemental Figure 2.**
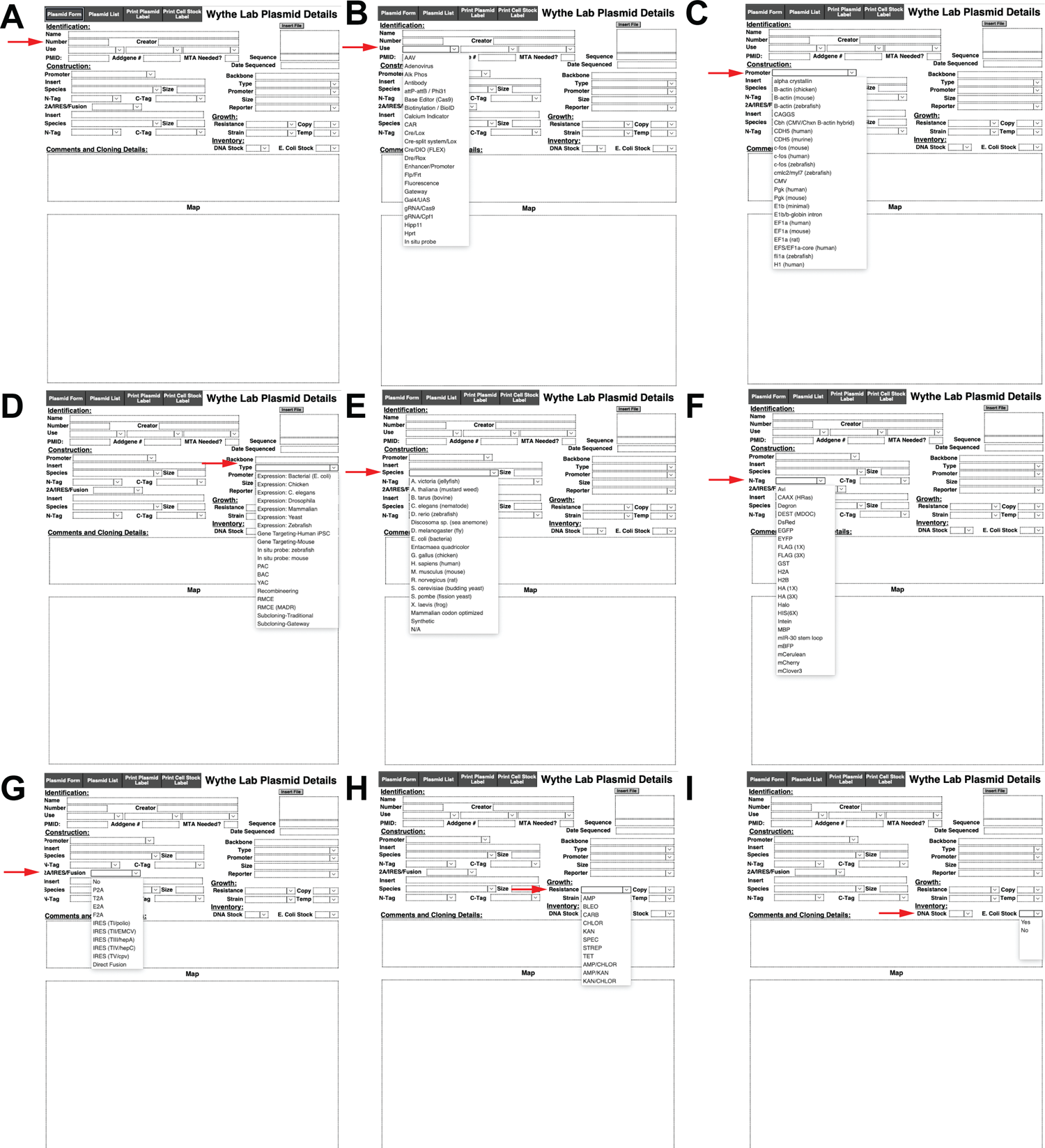

**Supplemental Figure 3.**
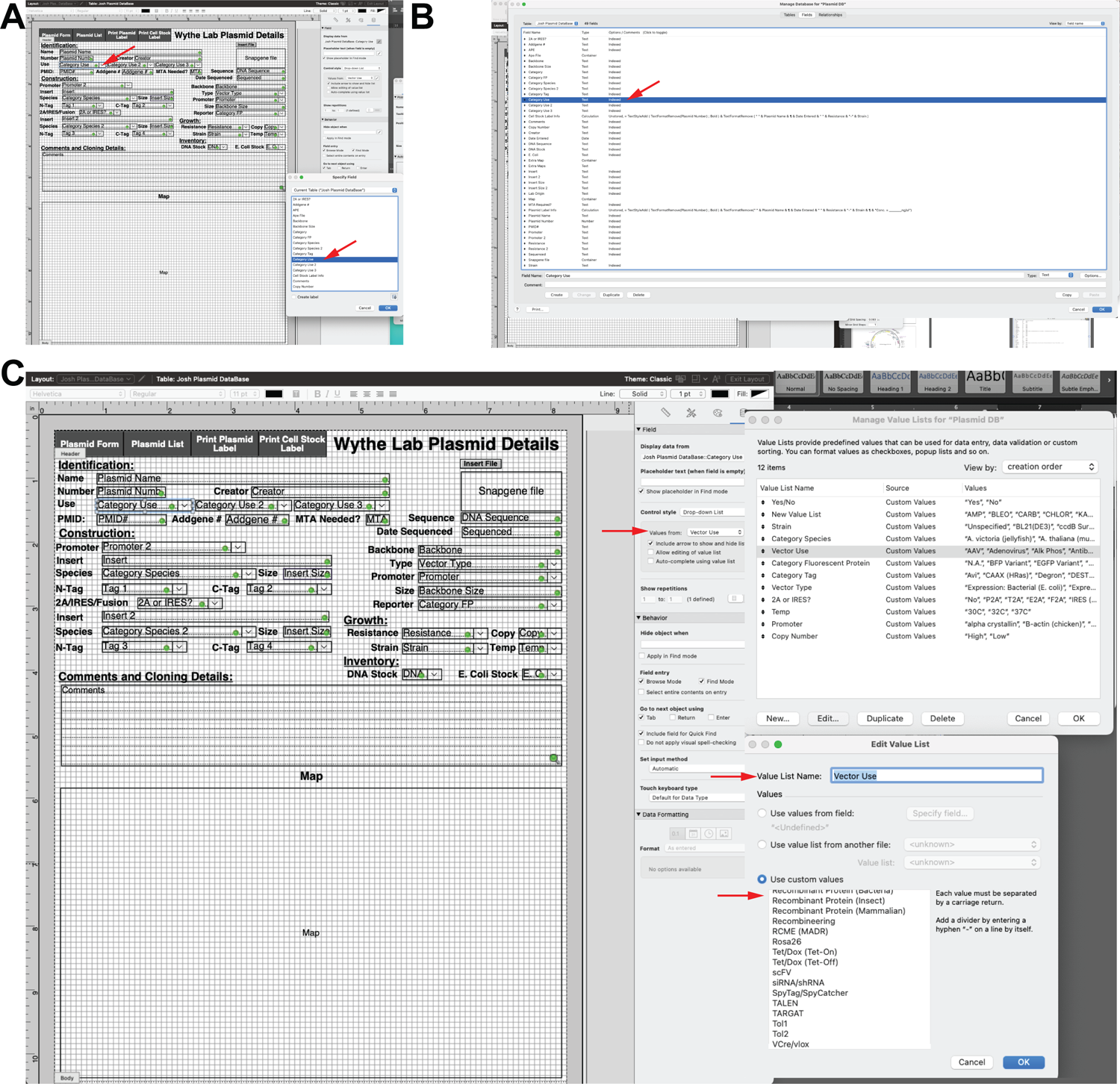

**Supplemental Figure 4.**
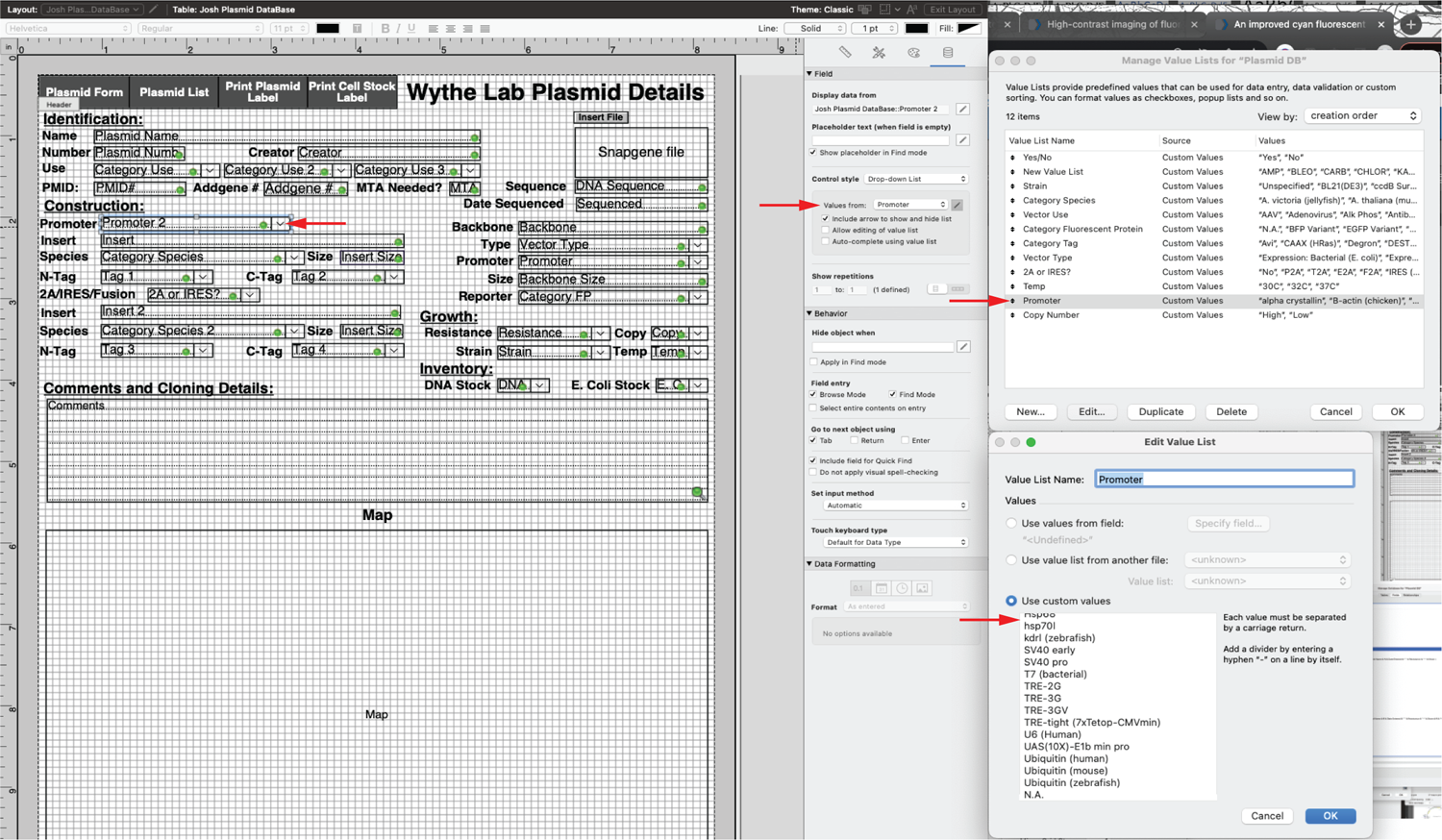

